# Formation of an Aminovinyl-Cysteine Residue in Thioviridamide Non-Lanthipeptides Occurs through a Path Independent of Known Lanthionine Synthetase Activity in *Streptomyces sp*. NRRL S-87

**DOI:** 10.1101/2020.08.21.260521

**Authors:** Yanping Qiu, Jingyu Liu, Yuqing Li, Yanqing Xue, Huan Wang, Wen Liu

## Abstract

2-<u>A</u>mino<u>vi</u>nyl-<u>cys</u>teine (AviCys) is an unusual thioether amino acid shared by a variety of ribosomally synthesized and posttranslationally modified peptides (RiPPs), as part of a macrocyclic ring system that contains the *C*-terminal 4 or 6 residues of a precursor peptide. This amino acid is nonproteinogenic and arises from processing the *C*-terminal Cys residue and an internal Ser/Thr residue to form an unsaturated thioether linkage. Enzyme activities for forming lanthionine (Lan), a distinct saturated thioether residue characteristic of lanthipeptide-related RiPPs, has long been speculated to be necessary for AviCys formation. Based on investigations into the biosynthesis of thioviridamide non-lanthipeptdes in *Streptomyces sp*. NRRL S-87, we here report an alternative path for AviCys formation that is independent of known Lan synthetase activity. This path relies on four dedicated enzymes for posttranslational modifications of the precursor peptide, in which TvaE_S-87_, a phosphotransferase homolog, plays a critical role. It works with LanD-like flavoprotein TvaF_S-87_ to form a minimum AviCys synthetase complex that follows the combined activity of TvaCD_S-87_ for Thr dehydration and catalyzes Cys oxidative decarboxylation and subsequent Michael addition of the resulting enethiol nucleophile onto the newly formed dehydrobutyrine residue for cyclization. With TvaE_S-87_, TvaF_S-87_ activity for Cys processing can be coordinated with TvaCD_S-87_ activity for minimizing competitive or unexpected spontaneous reactions and forming AviCys effectively.

## INTRODUCTION

Ribosomally synthesized and post-translationally modified peptides (RiPPs) are a large class of natural products that arise from post-translational modifications (PTMs) for precursor peptide maturation and the achievement of their biological activities.^1^ Using 20 proteinogenic amino acids as building blocks, nature creates stunningly diverse RiPPs largely because of unique enzymes for PTMs. The biosynthesis of thioviridamides (TVAs, **Figure 1A**),^2^ a growing group of structurally related thioamitide RiPPs with potent antitumor activity, is such a case. TVAs were initially isolated from *Streptomyces olivoviridis* and then produced in *S. lividans* where the TVA biosynthetic gene cluster (*tva*) was expressed heterologously.^3^ Recent survey of the published bacterial genome sequences revealed many *tva*-like gene clusters, and subsequent product mining in related microorganisms led to a rapid enrichment of the TVA family, in which several new members appear to be promising for cancer treatment.^4,5^ In addition to a characteristic thioamide peptide backbone, TVAs contain an unusual thioether residue, 2-<u>am</u>ino<u>vi</u>nyl-(3-methyl)-<u>cy</u>steine (AviCys),^6^ which is shared by a variety of RiPPs including linaridins (e.g., cypemycin^7^ and cacaoidin^8^) and certain lanthipeptide-related RiPPs (e.g., epidermin^9^ and microvionin lipolanthines^10^), as part of a macrocyclic ring system that contains the *C*-terminal 4 or 6 residues of a precursor peptide (**Figure 1A**).

**Figure 1.**
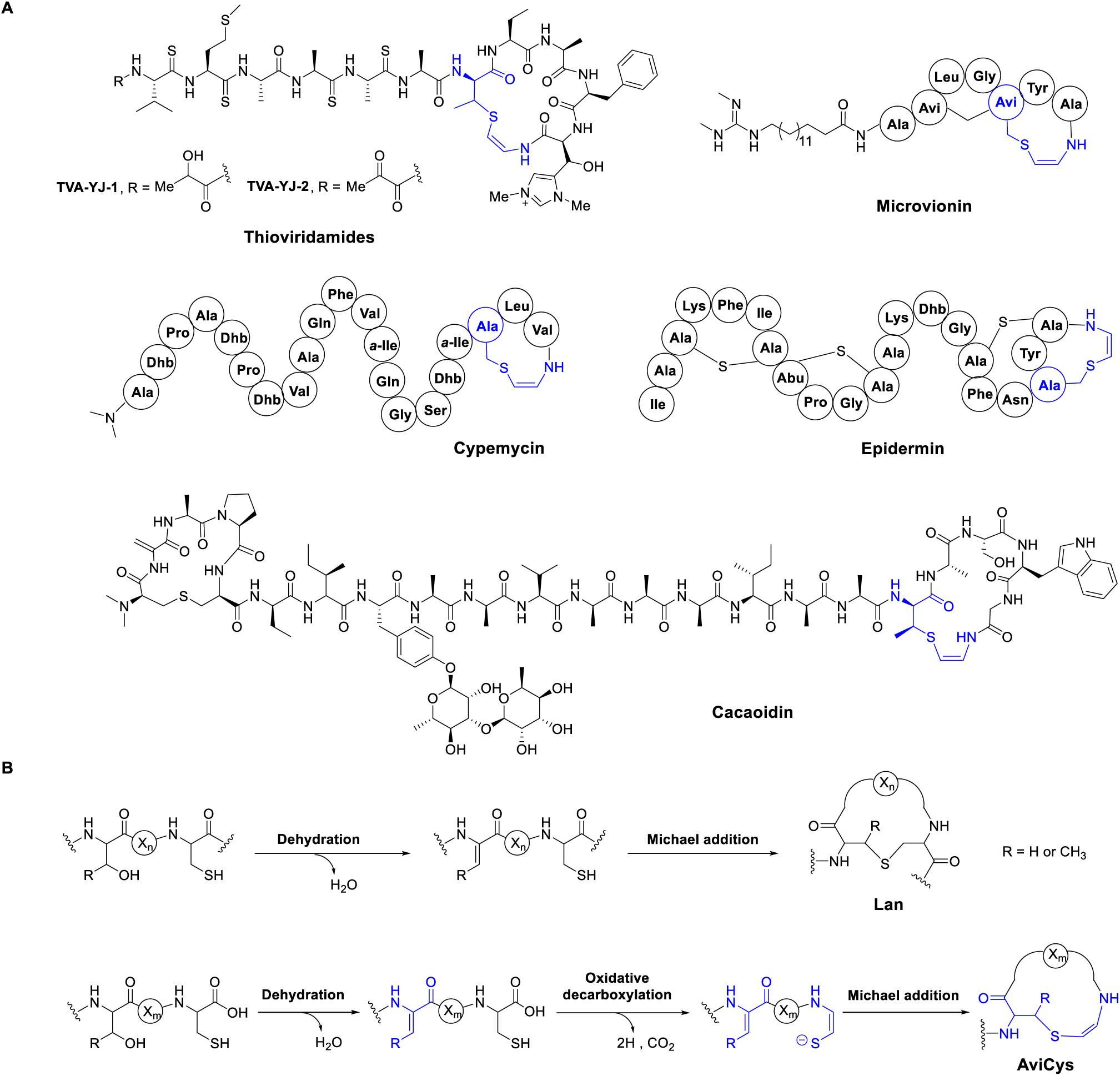
Thioether residue-associated structures and forming mechanisms. (**A**) Selected RiPPs containing an AviCys residue (blue). For the nonproteinogenic residues in addition to AviCys and Lan, Dhb, 2,3-didehydrobutyrine; Abu, α-aminobutyric acid; *a*-Ile, *allo*-isoleucine; and Avi, avionin. (**B**) Mechanistic comparison in the formation of thioether residues Lan (top) and AviCys (bottom). Modified residues during AviCys formation are labeled in blue.

The process through which an AviCys residue is formed remains poorly understood. Available insights have gained largely from the biosynthesis of lanthipeptide-related RiPPs, which, in fact, are characterized by a distinct thioether amino acid, (3-methyl-)<u>lan</u>thionine (Lan) (**Figure 1B**). The formation of a Lan residue has been well established over the past two decades.^11^ It results from dehydration of a Ser or Thr residue and subsequent cyclization by nucleophilic addition of a Cys residue onto the newly formed dehydroamino acid, 2,3-didehydroalanine (Dha) or 2,3-didehydrobutyrine (Dhb), to furnish a thioether linkage in the precursor peptide (**Figure 1B**). The two reactions, i.e., dehydration and cyclization that occur in tandem, can be attributed to the dedicated activities of a LanB dehydratase and a LanC cyclase for Class I lanthipeptides or to the combined activity of a multifunctional protein, i.e., LanM, LanKC or LanL for Class II, III or IV lanthipeptides, respectively (here collectively referred to as Lan synthetase).^11^ In comparison, the formation of an AviCys residue, which was proposed to share a similar biosynthetic route, relies on specifically processing the *C*-terminal Cys residue and the 2 or 4-aa-upstream, internal Ser/Thr residue of a precursor peptide (**Figure 1B**). It likely involves **1)** the dehydration of internal Ser or Thr and **2)** the oxidative decarboxylation of *C*-terminal Cys, followed by **3)** Michael addition of the resulting enethiol nucleophile onto the preceding Dha or Dhb residue to provide a distinct, unsaturated thioether linkage. In this route, processing *C*-terminal Cys to an enethiol, which is unique for AviCys formation, has been characterized to be dependent of the activity of LanD, a flavoprotein belonging to the family of homo-oligomeric, flavin-containing cysteine decarboxylases (HFCDs).^6^

Given mechanistic similarity in the formation of Lan and AviCys thioether residues, Lan synthetase activity for Ser/Thr dehydration and thiol addition/cyclization has long been speculated to be involved in AviCys formation.^6,12^ Recently, the combination of a LanKC-type, Class III multifunctional Lan synthetase with a LanD flavoprotein proved to be necessary for the formation of an AviCys-containing avionin motif in the biosynthesis of lipolanthines,^10^ supporting the relevance between these two distinct thioether residues in PTM enzymes. With great interest in how residue AviCys is formed and whether Lan synthetase activity participates in the biosynthesis of AviCys-containing non-lanthipeptides, we here investigated the biosynthetic pathway of TVA RiPPs in *S. sp*. NRRL S-87. Based on the characterization of a LanD-like, HFCD-fold flavoprotein and three previously functionally unknown, PTM-related enzymes, we demonstrated that in this *Streptomyces* strain, the formation of an AviCys residue does not require previously known Lan synthetase activity and instead occurs through a path biochemically similar to but phylogenetically different from that in the biosynthesis of lanthipeptide-related RiPPs.

## RESULTS

### TVA biosynthesis in *S. sp*. NRRL S-87 does not depend on Known Lan synthetase activity

We and others have recently validated the production of TVAs in *S. sp*. NRRL S-87, where a gene cluster (i.e., *tva*_*S-87*_) sharing nearly head-to-tail homology with *tva*, the prototype for TVA biosynthesis in *S. olivoviridis*, was identified (**Figures 2A** and **S2**).^2,4d^ Notably, the *tva*_*S-87*_ cluster does not encode any homologs of Lan synthetases, either monofunctional (i.e., LanB and LanC) or multifunctional (LanM, LanKC or LanL), consistent with the absence of residue Lan in the structure of TVAs. Whether TVA biosynthesis involves genes outside the *tva*_*S-87*_ cluster for Lan formation needed to be determined. In the genome of *S. sp*. NRRL S-87, we identified two genes potentially related to Lan formation, i.e., *lanKC1* and *lanKC2*, which are *tva*_*S-87*_-unclustered and encode two homologs of LanKC-type, Class III multifunctional Lan synthetases (**Table S2**). To exclude the participation of these two genes in TVA biosynthesis, we cloned and heterologously expressed the *tva*_*S-87*_ cluster in *S. laurentii*, a strain producing the thiopeptide antibiotic thiostrepton (**Figure 3A**).^13^ As expected, TVA-related compounds were observed, including TVA-YJ-1, the newly characterized member that shares an AviCys residue but differs from TVA-YJ-2 (previously identified in *S. sp*. NRRL S-87^4d^) by possessing a reduced lactyl unit instead of a pyruvyl unit at the *N*-terminus (**Figures 1A, S3** and **S25** and **Table S1**). Careful analysis of the *S. laurentii* genome revealed *lanL*_*SL*_, a gene coding for a homolog of LanL-type, Class IV multifunctional Lan synthetases (**Table S2**). We then inactivated *lanL*_*SL*_ in *S. laurentii* and introduced the *tva*_*S-87*_ cluster into the resulting mutant strain. As a result, the inactivation of *lanL*_*SL*_ had little effect on the production of TVAs in *S. laurentii* (**Figure 3A**), demonstrating that the *tva*_*S-87*_ cluster is sufficient for TVA biosynthesis and therefore contains all necessary genes for AviCys formation.

**Figure 2.**
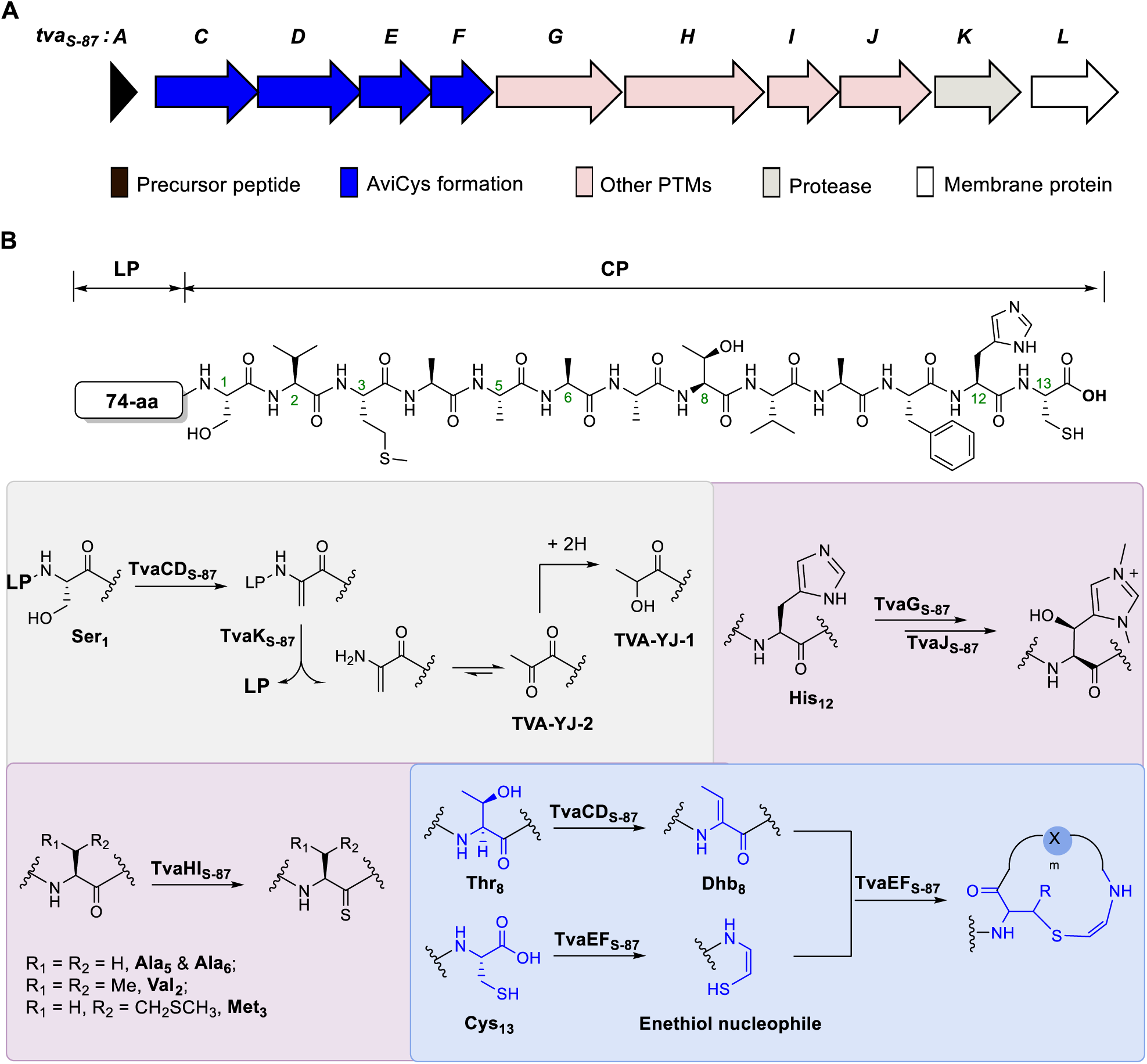
Gene cluster and proposed biosynthetic pathway of TVAs in *S. sp*. NRRL S-87. (**A**) Organization of genes responsible for the biosynthesis of TVAs. Gene functions are annotated by colored rectangular blocks. (**B**) Precursor peptide (TvaA_S-87_) and related PTMs in the biosynthesis of TVAs. Residues related to AviCys formation are highlighted in blue. The LP and CP parts of TvaA_S-87_ are indicated. The CP sequence is numbered. PTMs are highlighted by color, as grey for LP removal to form the *N*-terminal latyl and pyruvyl units of TVA-YJ-1 and TVA-YJ-2, respectively; light purple for His12 functionalization (top right) and thioamide formation (bottom left); and light blue for AviCys formation.

**Figure 3.**
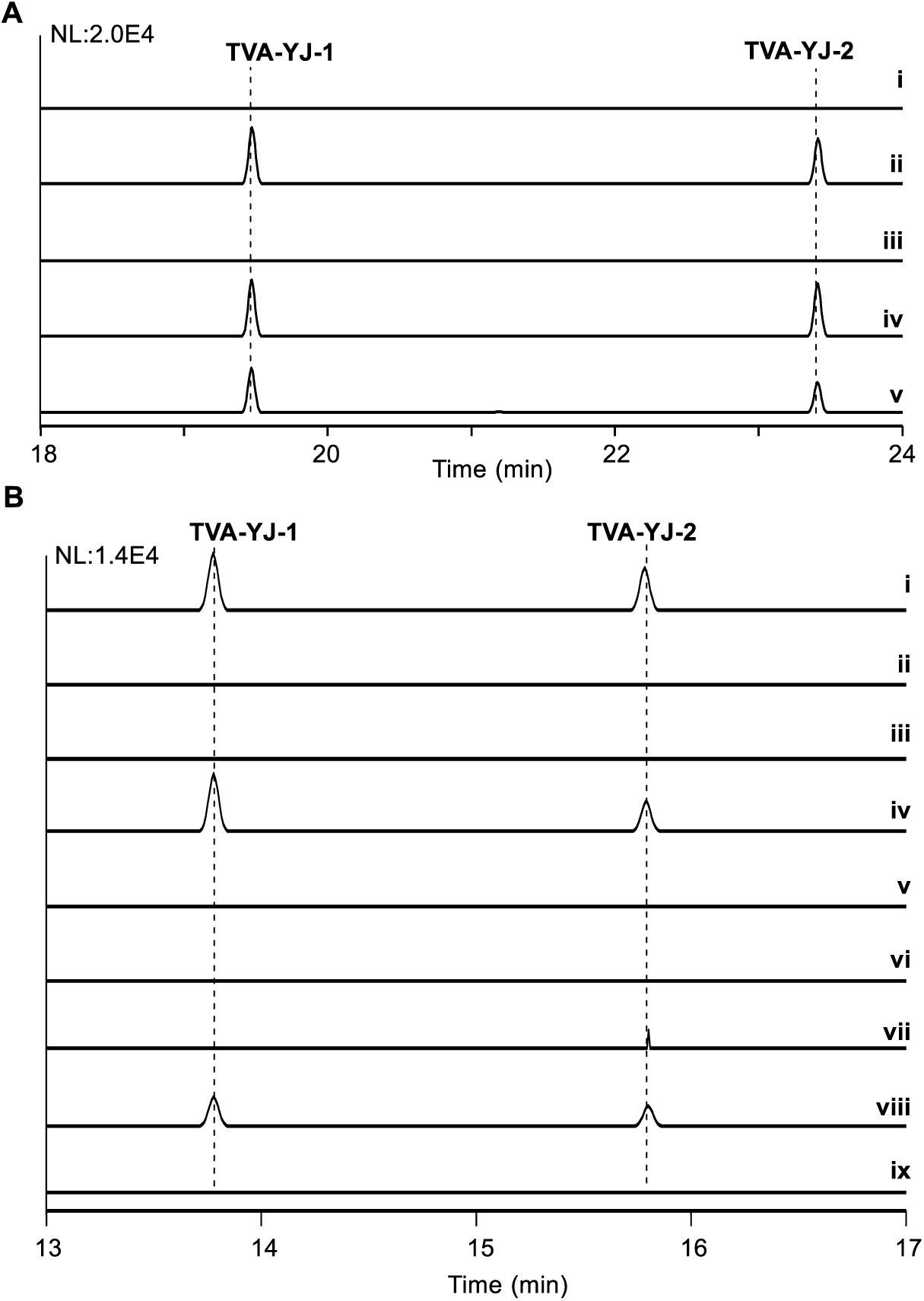
*In vivo* examination of TVA production by HPLC-MS. For extracted ion chromatograms (EICs), the ESI *m/z* M^+^ modes for TVA-YJ-1 and TVA-YJ-2 are 1305.5 and 1307.5, respectively. (**A**) Validation of the *tva*_*S-87*_ cluster (cloned from *S. sp*. NRRL S-87) by heterologous expression in *S. laurentii*. For HPLC-MS analysis, **Method I** was used (see **Supporting Information**). i, wild-type *S. laurentii*; ii, *S. laurentii* variant into which *tva*_*S-87*_ was introduced; iii, *ΔlanL*_*SL*_ *S. laurentii* mutant; iv, *ΔlanL*_*SL*_ *S. laurentii* mutant into which *tva*_*S-87*_ was introduced; and v, wild-type *S. sp*. NRRL S-87 (the TVA producer as a control). (**B**) Validation of the necessity of *tvaCDEF*_*S-87*_ for the biosynthesis of TVAs in *S. sp*. NRRL S-87. For HPLC-MS analysis, **Method II** was used (see **Supporting Information**). i, wild-type; ii, *ΔtvaA*_*S-87*_ mutant; iii, *ΔtvaC*_*S-87*_ mutant; iv, *ΔtvaC*_*S-87*_ mutant in which *tvaC*_*S-87*_ was expressed *in trans*; v, *ΔtvaC*_*S-87*_ mutant in which *tvaC*_*S-87*_*-N241A/D258A* was expressed *in trans*; vi, *ΔtvaD*_*S-87*_ mutant; vii, *ΔtvaE*_*S-87*_ mutant; viii, *ΔtvaE*_*S-87*_ mutant in which the mutated *tva*_*S-87*_ cluster was under the control of the promoter *PermE\**; and ix, *ΔtvaF*_*S-87*_ mutant.

### Identification of genes coding for AviCys formation in the *tva*_*S-87*_ cluster

In addition to *tvaA*_*S-87*_, the gene encoding the precursor peptide TvaA_S-87_ composed of a 74-aa leader peptide (LP) and the 13-aa core peptide (CP), **S**_**1**_**VMAAAAT**_**8**_**VAFHC**_**13**_ (numbering relates to residue position, as below), the *tva*_*S-87*_ cluster contains 10 functional genes, i.e., *tvaCDEFGHIKL*_S-87_, being organized into the same operon for co-expression to facilitate the coordination of their protein products in TVA biosynthesis (**Figure 2**). TvaL_S-87_ is a membrane protein potentially participating in TVA transportation, while the other 9 proteins were believed to be responsible for the transformation of the CP sequence of TvaA_S-87_ into mature molecules. During this PTM process, the LP sequence of TvaA_S-87_ was supposed to be necessary for the engagement of related enzyme catalysts and can be removed in the late biosynthetic stage as usual, assumedly by protease TvaK_S-87_.^4b^ During the maturation of TVAs, the specific PTMs include **1)** thioamidation of the peptidyl backbone, for which TvaH_S-87_ and TvaI_S-87_ might function together to form a complex composed of a discrete YcaO protein and a TfuA-fold protein;^14^ and **2)** functionalization within the AviMeCys-containing, *C*-terminal ring system, for which TvaG_S-87_ and TvaJ_S-87_ could work on the same residue, His12, for *N,N*-dimethylation and β-hydroxylation (**Figure 2B**).

Based on the above analysis, the PTM proteins for AviCys formation were narrowed to TvaF_S-87_, a LanD-like, HFCD-fold flavoprotein, along with remaining TvaC_S-87_, TvaD_S-87_ and TvaE_S-87_, all of which were functionally unassigned. The counterparts of TvaF_S-87_ were known in the biosynthesis of various AviCys-containing RiPPs. Either as a LanD protein (for lanthipeptides^10-12,15^) or as a LanD-like protein (for non-lanthipeptides^6,8,16^), these flavoproteins share the oxidative decarboxylation activity necessary for AviCys formation and can process the *C*-terminal Cys residue of a precursor peptide to an enethiol before Michael addition to the upstream Dha/Dhb residue. The homologs of TvaC_S- 87_ and TvaE_S-87_ were previously annotated as hypothetic proteins, with the exception in ref. 4b, where they were proposed to be phosphotransferases by Truman *et al*. Here, careful sequence analysis allowed the identification of a protein kinase domain in both TvaC_S-87_ and TvaE_S-87_, which share moderate sequence identity to each other (27.2 % identity). Different from TvaC_S-87_, which is likely an active phosphotransferase possessing the necessary residues for Ser/Thr activation and phosphorylation with adenosine triphosphate (ATP), TvaE_S-87_ appears to be an inactive homolog of phosphotransferases given the absence of these conserved residue in its sequence (**Figure S4**). TvaD_S-87_ is a hypothetic protein sharing no sequence homology with any proteins of known functions. Secondary structure prediction revealed that it resembles phosphothreonine lyases, the bacterial effector proteins capable of inactivating MAP kinase activity via phosphate elimination of a phosphorylated Thr residue, which leads to Dhb production, and thus attenuating the immune response of infected hosts (**Figure S5**).^17^ Consequently, while TvaE_S-87_ activity remained unclear based on sequence analysis alone, putative lyase TvaD_S-87_ likely associates with phosphotransferase TvaC_S- 87_ for the dehydration of the TvaA_S-87_ precursor peptide, specifically by targeting residues Ser1 and Thr8 (**Figure 2B**). Processing these two residues is a critical step in the proposed biosynthetic pathway of TVAs. Thr8 dehydration leads to a Dhb8-containing precursor to set the structural stage for AviMeCys formation, and Ser1 dehydration results in a Dha1-containing intermediate, where, following the hydrolysis of the LP sequence, the deamination of Dha1 can occur to form the *N*-terminal pyruvyl unit of TVA-YJ-2 (which might then undergo reduction to produce TVA-YJ-1).

### Necessity of *tvaCDEF*_*S-87*_ for the biosynthesis of TVAs

To determine the functions of *tvaC*_*S-87*_, *tvaD*_*S-87*_, *tvaE*_*S-87*_ and *tvaF*_*S-87*_ in TVA biosynthesis, we first inactivated these genes individually in *S. sp*. NRRL S-87 (**Figure 3B**). The precursor peptide-encoding gene *tvaA*_*S-87*_ was also inactivated, yielding the *ΔtvaA*_*S-87*_ mutant as a TVA non-producing, control strain. Compared to the wild-type strain, the *ΔtvaC*_*S-87*_, *ΔtvaD*_*S-87*_ and *ΔtvaF*_*S-87*_ mutants share a phenotype with the *ΔtvaA*_*S-87*_ mutant and failed to produce TVAs (**Figure 3B**). Phylogenetically, TvaC_S-87_ and TvaD_S-87_ share no sequence homology to any known LanM, LanKC or LanL-type, multifunctional Lan synthetases, suggesting that they evolve from different ancestors for dehydration activity development. Intriguingly, TVA-YJ-2 was identified in the *ΔtvaE*_*S-87*_ mutant, albeit with a significant decrease of yield. To ascertain TVA production, the *ΔtvaE*_*S-87*_ mutant was further engineered by introduction of a constitutive promoter, *PermE\**, to enhance the transcription and subsequent expression of the mutant *tva* cluster in which *tvaE*_*S-87*_ was deleted. Both TVA-YJ-2 and TVA-YJ-1 were observed, thereby indicating that *tvaE*_*S-87*_ is closely related to but not indispensable for the production of TVAs in *S. sp*. NRRL S-87.

### TvaC_S-87_ and TvaD_S-87_ activities in the dehydration of TvaA_S-87_

We utilized a heterologous expression strategy to examine dehydrated intermediates in *Escherichia coli*, where the precursor peptide-encoding gene *tvaA*_*S-87*_ was co-expressed with related PTM genes (**Figure 4A**). Expressing *tvaA*_*S-87*_ with *tvaCD*_*S-87*_ resulted in two dehydrated variants of TvaA_S-87_, i.e., **1** (− 18 Da, − 1 × H_2_O) and **2** (− 36 Da, − 2 × H_2_O), with a ratio of ∼3:1. Both variants were characterized by high-performance liquid chromatography with mass spectrometric (HPLC-MS) and high resolution (HR)-MS/MS detection (**Figures S6, S7** and **S8**). Compared to the major product **1**, which bears a Dhb residue derived from Thr8, the minor product **2** is di-dehydrated and has an additional Dha derived from the first Ser1 residue in the CP sequence of TvaA_S-87_. In contrast, expressing *tvaA*_*S-87*_ alone or co-expressing it with *tvaDE*_*S- 87*_ in *E. coli* only produced unmodified TvaA_S-87_. To access the dehydration mechanism, we took advantage of the above *E. coli* heterologous system that produces **1** and **2** and conducted site-specific mutagenesis to trace possible intermediate(s) (**Figures 4A** and **S9**). This approach can minimize unexpected effects resulting from the variety of constructions and thus facilitate product profile comparison. TvaD_S-87_ was subjected to the mutation of residues His24 and Arg26 to Ala at the proposed active site.^17^ This double mutation must have inactivated TvaD_S-87_, because the production of **1** and **2** was completely abolished. Instead, a new variant of TvaA_S-87_ (**3**), which likely arises from the TvaC_S-87_-catalyzed phosphorylation of residue Thr8, was accumulated in *E. coli*, consistent with the lack of lyase activity. We did not observe any putative TvaA_S-87_ variants in which Ser1 is phosphorylated. Most likely, Ser1 processing relies on completing the TvaD_S-87_-catalyzed conversion of **3** into **1** via phosphate elimination during **2** production.

**Figure 4.**
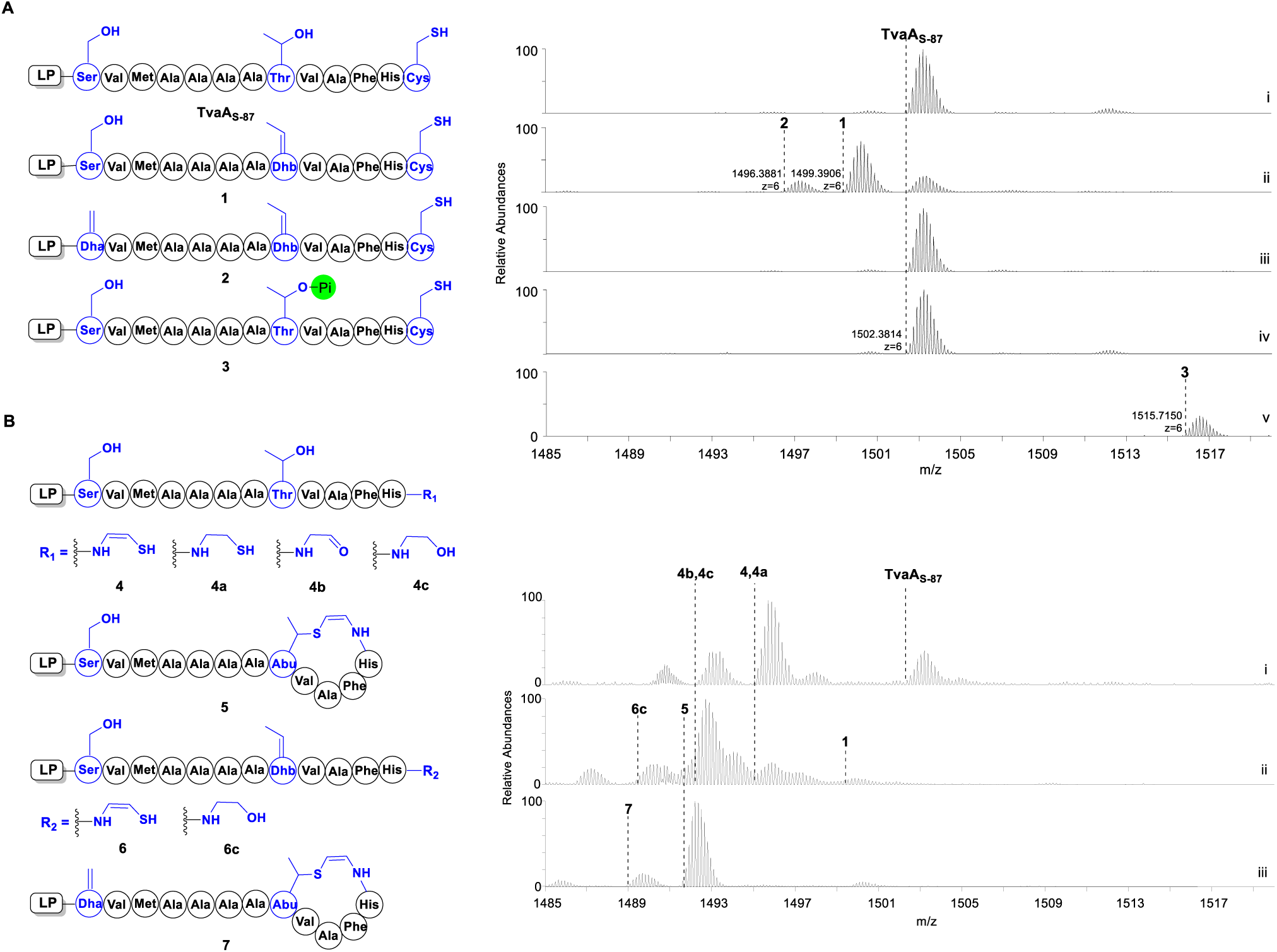
Heterologous expression of AviCys-forming genes in *E. coli*. Target residues for PTMs are labeled in blue. (**A**) Examination of TvaC_S-87_ and TvaD_S-87_ activities in the dehydration of TvaA_S-87_. The genes expressed in *E. coli* include precursor peptide-encoding gene *tvaA*_*S-87*_ alone (i), and the combination of *tvaA*_*S-87*_ with PTM-encoding genes *tvaCD*_*S-87*_ (ii); *tvaDE*_*S-87*_ (iii); mutant *tvaC*_*S-87*_*-N241A/D258A* and wild-type *tvaD*_*S-87*_ (iv); or wild-type *tvaC*_*S-87*_ and mutant *tvaD*_*S-87*_*-H24A/R26A* (v). (**B**) Examination of TvaE_S-87_ and TvaF_S-87_ activities in the formation of residue AviCys. The genes expressed in *E. coli* include the combination of *tvaA*_*S-87*_ with PTM-encoding genes *tvaF*_*S-87*_ (i); *tvaCDF*_*S-87*_ (ii); and *tvaCDEF*_*S-87*_ (iii).

Using the similar site-specific mutagenesis method, we examined the necessity of residues N241 and D258 in TvaC_S- 87_catalysis (**Figures 4A** and **S4**), where these residues are involved in the Mg^2+^-chelating of importance to kinase activity.^18^ Variants **1-3** were not produced by expressing *tvaA*_*S-87*_ with mutant *tvaC*_*S-87*_*-N241A/D258A* and wild-type *tvaD*_*S-87*_ in *E. coli*. Further, replacing wild-type *tvaC*_*S-87*_ with mutant *tvaC*_*S-87*_*-N241A/D258A* in the *tva* cluster abolished the production of TVAs in *S. sp*. NRRL S-87 (**Figure 3B**). These results supported the phosphotransferase activity of TvaC_S-87_ and the lyase activity of TvaD_S-87_. Considering expressing *tvaA*_*S-87*_ with *tvaC*_*S-87*_ alone failed to produce **3**, the two PTM enzymes might function together to form a two-component dehydratase complex, in which TvaD_S-87_ activity immediately follows TvaC_S-87_ activity to install Dhb8 first and Dha1 next in TvaA_S-87_.

### TvaF_S-87_ activity for the oxidative decarboxylation of TvaA_S-87_

We next expressed *tvaA*_*S-87*_ with *tvaF*_*S-87*_ in *E. coli*,to examine the LanD-like activity of TvaF_S-87_, i.e., for Cys13 oxidative decarboxylation in the treatment of the precursor peptide (**Figure 2B**). Analyzing the resulting product profile revealed a series of TvaA_S-87_-related peptides that differ at the *C*-terminus (**Figure 4B**). Among them, the − 46 Da (− [2H + CO_2_]) product was characterized to be variant **4** (**Figure S10**), which was believed to be the direct product of TvaF_S-87_ and arises from oxidative decarboxylation of the precursor peptide at the *C*-terminus. This enethiol variant is unstable and tends to undergo spontaneous conversions to various derivatives in *E. coli*, including thiol **4a** by hydrogenation, aldehyde **4b** by dethiolation and alcohol **4c** by further reduction (**Figures 4B** and **S10-S13**). Similar results were observed when assaying the activities of other LanD and LanD-like proteins involved in the biosynthesis of AviCys-containing RiPPs.^12,19^ Most likely, related enethiol intermediates are immediately transformed during AviCys formation, otherwise unexpected spontaneous conversions occur because of their high reactivity.

### TvaE_S-87_ activity is necessary for Effective AviMeCys formation

We then tested whether residue AviCys can be formed by combining TvaCD_S-87_ and TvaF_S-87_ activities together during the conversion of TvaA_S-87_ (**Figure 4B**). Compared with the combination of *tvaA*_*S-87*_ with *tvaF*_*S-87*_ alone in *E. coli*, expressing *tvaA*_*S-87*_ with *tvaCD*_*S-87*_ and *tvaF*_*S-87*_ further complicated the product profile, in which the individual products of both TvaCD_S-87_ (e.g., **1**) and TvaF_S-87_ (eg., **4** and its derivatives **4a-c**) were observed, indicating the competition of TvaCD_S-87_ and TvaF_S-87_ activities in the utilization of substrate TvaA_S-87_. Intriguingly, we observed a small amount of **5**, a − 64 Da (i.e., − [H_2_O + 2H + CO_2_]), relatively stable variant of the precursor peptide (**Figures 4B** and **S14**). Thus, dehydration and oxidative decarboxylation occurred. HR-MS/MS analysis narrowed the difference of this variant from TvaA_S-87_ to the 6-aa *C*-terminal sequence, **T_8_VAFHC_13_**. Unlike TvaA_S-87_ and the above linear variants, no fragment ions were observed within this sequence when analyzing **5**. It was resistant to thiol derivatization with iodoacetamide (IAA); in contrast, TvaA_S-87_ and its thiol-containing variants were sensitive to this treatment, yielding the derivatives conjugated by one IAA molecules (**Figures S15** and **S16**). Together, **5** was deduced to be the cyclized variant of TvaA_S-87_ that contains AviCys. This variant likely results from intramolecular Michael addition that follows TvaCD_S- 87_-catalyzed Thr8 dehydration and TvaF_S-87_-catalyzed, Cys13 oxidative decarboxylation for cyclizing the resulting linear intermediate (**6**) (**Figure 4B**). Similar to **4**, enethiol intermediate **6** appears to be highly reactive and thus was not observed in the *E. coli* system where *tvaACDF*_*S-87*_ were overexpressed. The presence of this linear intermediate was supported by the observation of **6c** (**Figures 4B** and **S17**), the Dhb8-containing alcohol derivative of **6**. Although residue AviCys can be formed, the accumulation of the above linear TvaA_S-87_ variants/derivatives indicates the lack of functional cooperation between TvaCD_S-87_ and TvaF_S-87_ activities and particularly the inefficiency of the subsequent cyclization step.

We further added *tvaE*_*S-87*_ into the above *E. coli* system in which *tvaA*_*S-87*_ was expressed with *tvaCD*_*S-87*_ and *tvaF*_*S-87*_, to examine whether the efficiency in AviCys formation can be improved. Remarkably, the product profile was significantly simplified, leaving **5** and **7**, the two AviCys-containing cyclized variants of TvaA_S-87_ with relatively high yields (**Figures 4B** and **S18**). Compared to **5, 7** contains an additional dehydrated residue, Dha1, which is derived from Ser1 by TvaCD_S-87_ activity. Clearly, the incorporation of TvaE_S-87_ activity greatly facilitates AviCys formation. Substrate TvaA_S-87_ and its linear variants/derivatives arising from TvaCD_S-87_ activity (e.g., **1** and **2**), TvaF_S-87_ activity (e.g., **4** and **4a-c**) or both of them (e.g., **6** and **6c**) were not observed or accumulated in the *E. coli* system where *tvaACDEF*_*S-87*_ were overexpressed (**Figure 4B**). This finding is important and indicates that TvaE_S-87_ activity not only accelerates the cyclization step but also coordinates TvaCD_S-87_ and TvaF_S-87_ activities down a path for effective AviCys formation.

### TvaE_S-87_ interacts with both TvaF_S-87_ and LP-containing substrate during AviCys formation

To examine whether dehydrated TvaA_S-87_ can serve as the precursor for AviCys formation, we overexpressed and purified TvaE_S- 87_ and TvaF_S-87_ from *E. coli* for *in vitro* assays (**Figure S1**). Similar to previously characterized LanD and LanD-like proteins, purified TvaF_S-87_ appeared light yellow and exhibited an absorbance spectrum characteristic of flavin in oxidized form. The non-covalently bound cofactor, which can be released from TvaF_S-87_ by heating, was confirmed to be flavin mononucleotide (FMN) (**Figure S19**). In terms of substrate preparation, dehydrated TvaA_S-87_ variants appeared to be highly unstable, and only Dhb8-containing **1** was isolated from the *E. coli* system in which *tvaA*_*S-87*_ was expressed with *tvaCD*_*S-87*_. Over a 2-hr incubation period at 30°C, this mono-dehydrated variant was completely degraded in buffer solution (**Figure 5**). When LanD-like flavoprotein TvaF_S-87_ was added into this solution, we did observe a distinct − 46 Da product (− [2H + CO_2_]) (**Figure 2**), which was then characterized to be **5** by HPLC-HR-MS, HR-MS/MS and IAA derivatization as aforementioned. Thus, **1** can be processed to the AviCys-containing, cyclized variant of TvaA_S-87_ in the presence of TvaF_S-87_, presumedly through the formation of linear enethiol intermediate **6**. Although TvaE_S-87_ alone had no effect on **1** conversion, the incubation of **1** with both TvaE_S-87_ and TvaF_S-87_ led to a remarkable improvement in the production of **5**, with a yield > 5-fold higher than that by incubating **1** with TvaF_S-87_ alone (**Figure 5**). Unambiguously, TvaE_S-87_ activity is not absolutely necessary but indeed facilitates AviCys formation, consistent with the above *in vivo* results obtained from gene inactivation in *S. sp*. NRRL S-87 and co-expression in *E. coli*.

**Figure 5.**
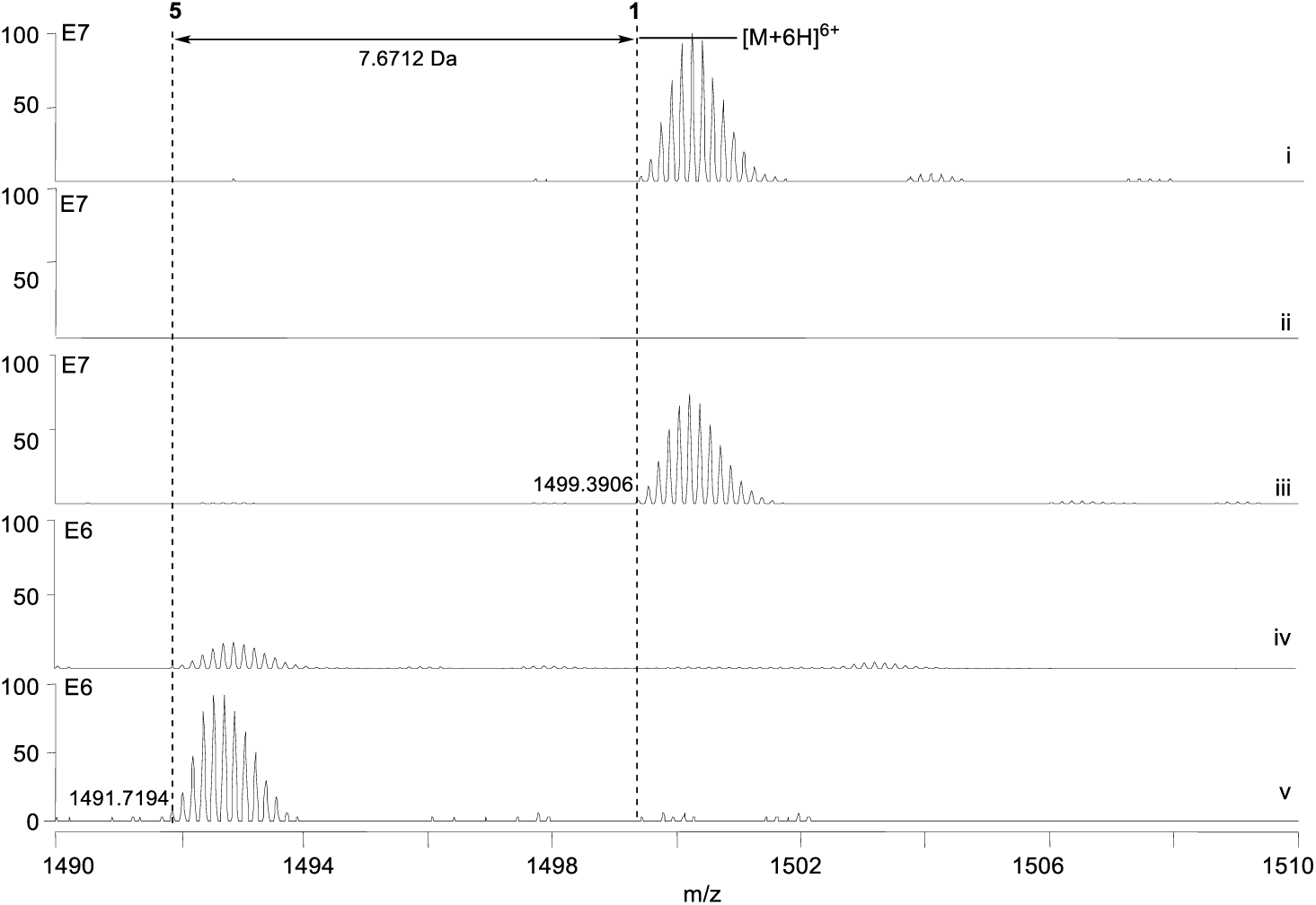
*In vitro* conversion of substrate **1** (in which residue Thr8 is dehydrated to Dhb8). For HPLC-MS analysis, the ESI *m/z* [M+H]^6+^ modes for **1** and cyclized AviCys-containing product **5** were shown. Tested samples include newly prepared **1** (i), the incubations of **1** alone at 30°C for 2-hr (ii) or with TvaE_S-87_ (iii), TvaF_S-87_ (iv) and TvaEF_S- 87_ (v), respectively, at 30°C for 2-hr.

During *in vitro* assays, we observed that TvaE_S-87_ can stabilize substrate **1**. With this protein, the rapid degradation of **1** was largely prevented over a 2-hr incubation period at 30°C. This observation led to the hypothesis that TvaE_S- 87_ interacts with the precursor peptide-derived substrate, assumedly by targeting its LP sequence part. To support this hypothesis, we chemically synthesized the LP and CP sequences of TvaA_S-87_ and measured their individual interactions with TvaE_S-87_ by microscale thermophoresis (MST) analysis (**Figure S20**).^20^ As expected, this analysis revealed a *K*_d_ value of 29.3 ± 5.1 nM between the LP sequence and TvaE_S-87_, indicating a strong interaction. In contrast, interaction with TvaE_S-87_ was not observed when using a 21-aa sequence containing CP (for solubilization, this sequence contains the additional 8 LP amino acids at the *N*-terminus of CP). In addition, we observed a moderate interaction between TvaE_S-87_ and LanD-like flavoprotein TvaF_S-87_ by MST, as supported by a *K*_d_ value of 1.1 ± 0.3 μM. These results suggested the notion that TvaE_S-87_ and TvaF_S-87_ function together by forming a heterologous complex, which can be engaged by binding tightly with the LP sequence of the precursor peptide to oxidatively decarboxylate the *C*-terminal Cys residue and immediately mediate subsequent Michael addition for macrocyclization and AviCys formation.

## DISCUSSION & CONCLUSION

Recent studies on the biosynthesis of Class III lanthipeptide-related RiPPs demonstrated that previously defined activities for Lan thioether formation is also involved in the formation of a distinct AviCys thioether residue.^10^ Based on *in vivo* and *in vitro* investigations into the biosynthesis of TVA non-lanthipeptides in *S. sp*. NRRL S-87, we here provided an alternative path for AviCys formation during the establishment of the *C*-terminal macrocyclic ring system of these thioamide RiPPs. This path is mechanistically similar to that for Lan formation; however, it does not involve any activities of known Lan synthetases, either monofunctional (i.e., LanB and LanC for Class I lanthipeptides) or multifunctional (LanM, LanKC or LanL for Class II, III or IV lanthipeptides).

In *S. sp*. NRRL S-87, effectively forming the AviCys residue of TVAs relies on the activities of four dedicated pathway-specific PTM enzymes. Specifically, phosphotransferase TvaC_S-87_ and lyase TvaD_S-87_ are functionally equivalent to but phylogenetically different from the kinase and lyase domains within Class II, III or IV Lan synthetases (i.e., LanM, LanKC or LanL multifunctional proteins). Most likely, they work together to form a two-component complex catalyzing the dehydration of TvaA_S-87_ (**Figure 6A**), by targeting Thr8 first and Ser1 next to install Dhb8 and Dha1 tandemly into the precursor peptide. Similar to previously characterized LanD-like HFCD-fold flavoproteins, TvaF_S-87_ has oxidative decarboxylation activity and is able to process the *C*-terminal Cys residue of TvaA_S-87_ to provide an enethiol group when functioning alone. We did observe AviCys formation by combining the activities of TvaCD_S-87_ and TvaF_S-87_ in the conversion of TvaA_S-87_, albeit with a poor yield. However, competition and a lack of coordination between Thr8 dehydration and Cys13 oxidative decarboxylation complicated the product profile. In particular, the cyclization of **6**, the precursor bearing Dhb8 and the enethiol group derived from Cys13, appears to occur in an inefficient manner, as the accumulation of this unstable intermediate led to spontaneous conversions to produce various derivatives including alcohol **6c**.

**Figure 6.**
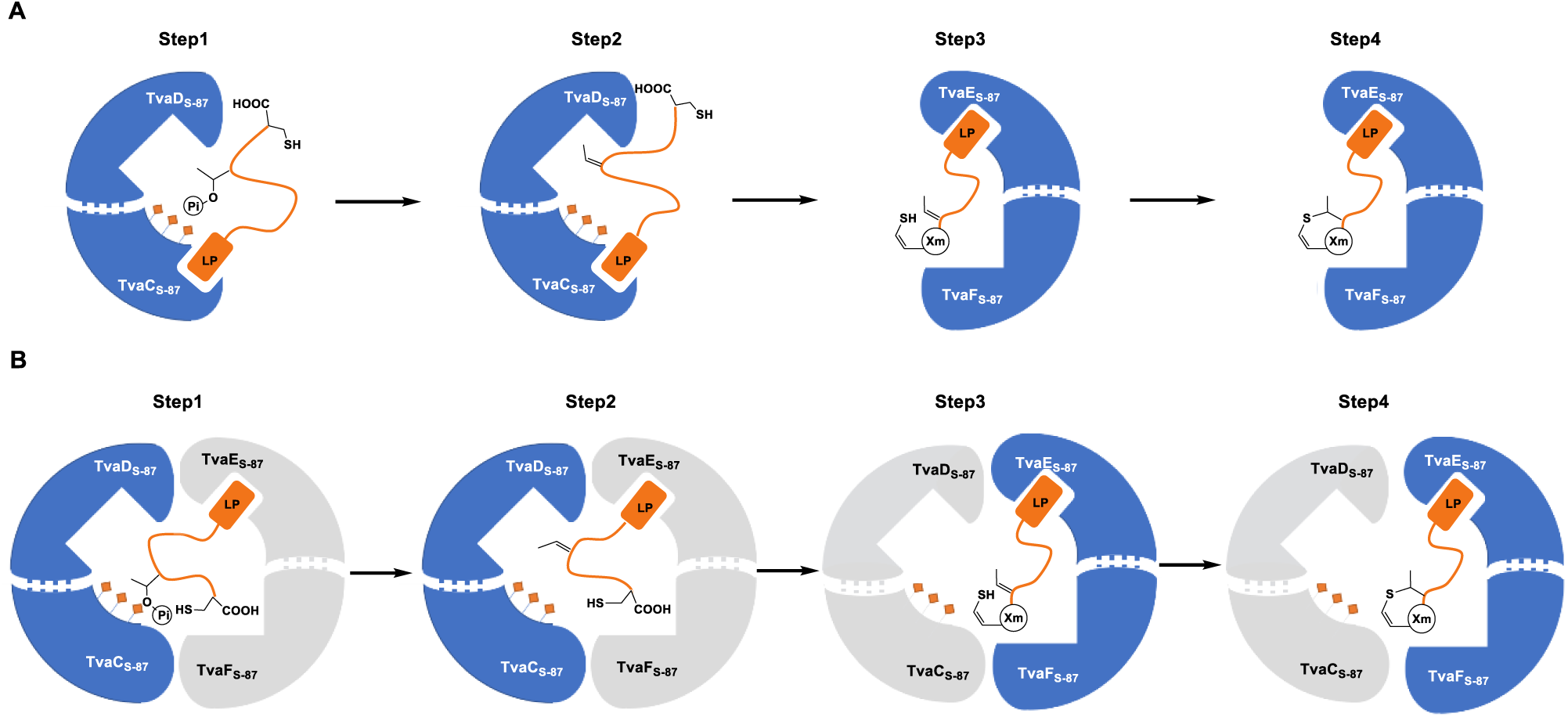
Proposed modes for AviCys formation during the biosynthesis of TVAs in *S. Sp*. NRRL S-87. Step 1, Thr phosphorylation; Step 2, H2O elimination; Step 3, Cys oxidative decarboxylation; and Step 4, cyclization. (**A**) Two component, TvaCD_S-87_ dehydratase complex and minimum TvaEF_S-87_ AviCys synthetase complex. In this mode, phosphotransferase TvaC_S-87_ is proposed to share with phosphotransferase homolog TvaE_S-87_ the ability for LP binding according to ref. 22. (**B**) Four component TvaCDEF_S-87_ AviCys synthetase super-complex.

The phosphotransferase homolog TvaE_S-87_ plays a critical role in effective AviCys formation. This protein, which does not have phosphotransferase activity, can interact with both TvaF_S-87_ and particularly the LP sequence of the precursor peptide. Benefiting from these interactions for trapping unstable substrate/intermediate and minimizing unexpected conversions, it could function with TvaF_S-87_ to form a two-component, minimum AviCys synthetase complex (**Figure 6**). In this complex, TvaE_S-87_ can be a dedicated cyclase catalyzing Michael addition between Dhb8 and the *C*-terminal enethiol group; alternatively, it serves as a noncatalytic engaging protein to mediate the putative dual activity of TvaF_S-87_ (i.e., for Cys13 processing and cyclization). Intriguingly, this phosphotransferase homolog can coordinate TvaCD_S-87_-catalyzed Thr8 dehydration and TvaF_S-87_-catalyzed Cys13 processing to avoid their competition in substrate, leading to the further hypothesis that similar to the MicKC-MicD and SpaKC-SpaF complexes in the biosynthesis of microvionin lipolanthines and TVA-like thiosparsoamide, respectively,^10,21^ TvaCD_S- 87_ and TvaEF_S-87_ may form a four component super-complex for introducing an AviCys residue into an unmodified precursor peptide (**Figure 6B**). Indeed, the SpaKC-encoding gene proved to be capable of compensating the loss of *tvaC*_*S-87*_, *tvaD*_*S-87*_ or *tvaE*_*S-87*_ in *S. sp*. NRRL S-87, supporting that this LanKC-type Lan synthetase is functionally equivalent to the combination of TvaC_S-87_, TvaD_S-87_ and TvaE_S-87_ (**Figure S21**). This observation indicates that nature employs a convergent evolution strategy to develop AviCys synthetase activity from different protein ancestors. At the current stage, we do not exclude the possibility that Cys processing precedes Thr dehydration during AviCys formation in this proposed super-complex because enethiol intermediate is unstable and cannot be isolated for *in vitro* assays.

Recently, Tao *et al*. identified lexapeptide, a new RiPP containing both Lan and AviCys thioether residues, and the associated biosynthetic gene cluster in *S. rochei* Sal35.^16^ Using a similar heterologous co-expression method in *E. coli*, four dedicated PTM enzymes, i.e., LxmKYXD, were indicated to be necessary for the formation of the AviCys residue in lexapeptide. Remarkably, despite poor homology in primary sequence, secondary structure prediction revealed that LxmKYXD are the counterparts of TvaCDEF_S-87_ in this study. It should be noted that we here reassigned the functions of LxmY and LxmX based on the above analysis in both primary sequence and secondary structure (**Figure S22**). LxmY can be a lyase functioning with phosphotransferase LxmK, which appears to be active and possesses all necessary residues for Ser/Thr activation and phosphorylation with ATP; in contrast, LxmX, which lacks conserved active site residues, is a phosphotransferase homolog proposed to be functionally equivalent to TvaE_S-87_ in *S. sp*. NRRL S-87. Whether the activities associated with LxmKYX are involved in the formation of the Lan residue occurring in lexapeptide remains to be determined. If so, LxmX and its counterpart TvaE_S-87_ in TVA biosynthesis more likely works as a new type, dedicated cyclase functionally corresponding to the cyclase domain of the LanKC-like proteins MicKC and SpaKC (**Figures S23**). However, whether Lan synthetase-encoding gene(s) outside the *lxm* cluster is the involved in the formation of the Lan residue in lexapeptide needs to be clarified. Determining the role of the TvaE_S-87_/LxmX-represented component, either as a cyclase or as an engaging protein, in related AviCys synthetase complex would be extremely interesting. In addition, genome survey revealed a number of genes coding for *tvaE*_*S-87*_/*lxmKYXD* homologs, which appear to be clustered with different PTM-encoding genes (**Figures S24**), suggesting that the path demonstrated here is widely employed in bacteria for the biosynthesis of diverse AviCys-containing RiPPs. The study presented here provides insights into the biosynthesis of TVA non-lanthipeptides that remains poorly understood. Using chemoenzymatic or synthetic biology strategies, enzymatic AviCys formation has potential for peptide drug development by application in peptide macrocyclization and functionalization, which remain a challenge to current synthesis approaches.

## MATRIALS & METHODS

For details of the materials and methods used in this study, please see **Supporting Information**.

## ASSOCIATED CONTENT

### Supporting Information

The Supporting Information is available free of charge on the ACS Publications website at DOI: xxxxx Supplementary materials and methods, results, **Figures S1−S25**, and **Tables S1−S4**.

## AUTHOR INFORMATION

\# These authors contributed equally to this work.

### Corresponding Author

*

### Notes

The authors declare no competing financial interest.

## ACKNOWLEDGMENT

We thank Zhijun Tang for discussion during manuscript preparation and Dr. Hao-Yang Wang at the laboratory of Mass Spectrometry Analysis for helps in the mass data collection. This work was supported in part by grants from MST (2019YFA0905400, 2018YFA090103 and 2018ZX091711001-006-010), NSFC (21520102004, 21750004 and 21621002), STCSM (17JC1405200), CAS (XDB20020200) (for L.P.); and NSFC (21750004 and 21520102004), CAS (QYZDJ-SSW-SLH037 and XDB20020200), STCSM (17JC1405100) and K. C. Wang Education Foundation.

## Notes

### Competing Interest Statement

The authors have declared no competing interest.

