## Supplementary Materials for "Formation of an Aminovinyl-Cysteine Residue in Thioviridamide Non-Lanthipeptides Occurs through a Path Independent of Known Lanthionine Synthetase Activity in *Streptomyces sp*. NRRL S-87"

**1. Supplementary Methods**

**1.1 General Materials and Methods**

**1.2 Gene Inactivation and Complementation**

**1.3 Gene Cluster Engineering and Heterologous Expression**

**1.4 Production and Analysis of TVAs**

**1.5 Protein Expression and Purification.**

**1.6 *In vitro* Assays of TvaE$_{S\text{-}87}$ and TvaF$_{S\text{-}87}$ Activities**

**1.7 Measurement of Protein-Protein Interactions**

**2. Supplementary Figures**

**Figure S1.** Purified recombinant enzymes analysis.

**Figure S2.** Alignment of the *tva*$_{S\text{-}87}$ cluster.

**Figure S3.** Characterization of the production of TVA-YJ-1.

**Figure S4.** Sequence alignment.

**Figure S5.** Structural homology.

**Figure S6.** Characterization of TvaA$_{S\text{-}87}$.

**Figure S7.** Characterization of **1**.

**Figure S8.** Characterization of **2**.

**Figure S9.** Characterization of **3**.

**Figure S10.** Characterization of **4**.

**Figure S11.** Characterization of **4a**.

**Figure S12.** Characterization of **4b**.

**Figure S13.** Characterization of **4c**.

**Figure S14.** Characterization of **5**.

**Figure S15.** Characterization of **TvaA$_{S\text{-}87}$-IAA**.

**Figure S16.** Characterization of **4a-IAA**.

**Figure S17.** Characterization of **6c**.

**Figure S18.** Characterization of **7**.

**Figure S19.** Characterization of the FMN-binding manner of the TvaF$_{S\text{-}87}$.

**Figure S20.** MST assay.

**Figure S21.** *In vivo* examination of TVA production.

**Figure S22.** Structural homology.

**Figure S23.** Functional alignment of known AviCys synthetase-encoding genes.

**Figure S24.** Selected gene clusters containing the homologs of *tvaCDEF*$_{S\text{-}87}$ and their associated bacterial strains.

**Figure S25.** NMR spectra of TVA-YJ-1.

### 3. Supplementary Tables

**Table S1.** NMR data (600MHz, DMSO-d6) for TVA-YJ-1.

**Table S2.** Lan synthase homologs in the genome of *Streptomyces sp.* NRRL S-87 and *Streptomyces laurentii*.

**Table S3.** Related bacterial strains and plasmids used in this study.

**Table S4.** Primers used in this study.

**References**

### 1. Supplementary Methods

#### 1.1 General Materials and Methods

**Materials, bacteria strains and plasmids.** Biochemicals and media were purchased from Sinopharm Chemical Reagent Co., Ltd. (China), Oxoid Ltd. (U.K.) or Sigma-Aldrich Corporation (USA) unless otherwise stated. Restriction endonucleases were purchased from Thermo Fisher Scientific Co. Ltd. (USA). Chemical reagents were purchased from standard commercial sources. Synthetic peptides were purchased from Genscript Biotech (Nanjing, China). Related bacterial strains and plasmids are summarized in **Table S3**. Primers used in this study are listed in **Table S4**.

**DNA isolation, manipulation, and sequencing.** DNA isolation and manipulation in *Escherichia coli* or *Streptomyces* strains were carried out according to standard methods.[1] PCR amplifications were carried out on an Applied Biosystems Veriti™ Thermal Cycler either using Taq DNA polymerase (Vazyme Biotech Co. Ltd, China) for routine genotype verification or PrimeSTAR HS DNA polymerase (Takara Biotechnology Co., Ltd. Japan) for high fidelity amplification. The synthesis of primers and genes were performed at Shanghai Sangon Biotech Co., Ltd. (China). DNA sequencing was performed at Shanghai Biosune Biotech Co., Ltd. (China).

**Sequence analysis.** Biosynthetic gene clusters (BGCc) were mined from microbial genomes using the AntiSMASH web tool.[2] Open reading frames (ORFs) were identified using the FramePlot 4.0beta program (http://nocardia.nih.go.jp/fp4/).[3] The deduced proteins were compared with other known proteins in the databases using available BLAST methods (http://blast.ncbi.nlm.nih.gov/Blast.cgi).[4] Amino acid sequence alignments were performed using Vector NT1 and ESPript 3.0 (http://espript.ibcp.fr/ESPript/ESPript/).[5] Gene cluster diagrams were made with MultiGeneBlast (http://multigeneblast.sourceforge.net/).[6]

**Chemical analysis.** Analysis by High Performance Liquid Chromatography (HPLC) were carried out on an Agilent 1260 HPLC system (Agilent Technologies Inc., USA). Analyses by HPLC-associated Electrospray ionization Mass Spectrometer (ESI-MS) and ESI-high resolution MS (ESI-HR-MS) were performed on a Thermo Fisher LTQ XL ESI-MS spectrometer and a Q Exactive™ Plus Mass Spectrometer (Thermo Fisher Scientific Inc., USA), respectively. Related data were processed using Thermo Xcalibur software. NMR data were recorded on a Bruker AV500 spectrometers (Bruker Co. Ltd., Germany) or on an Agilent PremiumCompact+ 500MHz NMR spectrometer (Agilent Technologies Inc., USA).

#### 1.2 Gene Inactivation and Complementation

**Gene Inactivation.** The gene *tvaA*$_{S-87}$ was inactivated in *Streptomyces sp.* NRRL S-87 through a double-crossover recombinant event using a method described previously.[7] The 1.92-kb fragment obtained using the primers *tvaA*$_{S-87}$-R-for/*tvaA*$_{S-87}$-R-rev and the 2.04-kb fragment obtained using the primers *tvaA*$_{S-87}$-L-for/ *tvaA*$_{S-87}$-L-rev were cloned into pMD19-T to yield pYJ1001 and pYJ1002, respectively. These two fragments were recovered from pYJ1001 and pYJ1002 and then cloned into pOJ260, yielding pYJ1003, in which a 180-bp in-frame coding region of *tvaA*$_{S-87}$ was deleted. This recombinant plasmid was introduced into *S. sp.* NRRL S-87 by conjugation between *E. coli* ET12567-Streptomyces. The colonies that were apramycin-resistant at 30°C were identified as integrating mutants, in which a single-crossover homologous recombination event took place. These mutants were cultured for several rounds in the absence of apramycin, and the resulting apramycin-sensitive isolates were subjected to PCR amplification to examine the genotype of the *S. sp.* NRRL S-87 mutant strain YJ101.

The gene *tvaC*$_{S-87}$, *tvaD*$_{S-87}$, *tvaE*$_{S-87}$ and *tvaF*$_{S-87}$ were inactivated individually in *S. sp.* NRRL S-87 using a modified CRISPR-Editing method.[8] As exemplified by the inactivation of *tvaC*$_{S-87}$, a 20-bp sg sequence was synthesized and cloned into the plasmid pWHU2653 through homologous recombination among the two ~140-bp PCR products amplified from pWHU2653 using the two primer pairs sg-for/*tvaC*$_{S-87}$-sg-rev and *tvaC*$_{S-87}$-sg-for/sg-rev, respectively, and NheI/XbaI-digested pWHU2653, yielding pYJ1004. A truncated *tvaC*$_{S-87}$ gene was then constructed into pYJ1004 through homologous recombination among the two 2-kb PCR products amplified from the genome of *S. sp.* NRRL S-87 using the two primer pairs *tvaC*$_{S-87}$-L-for/ *tvaC*$_{S-87}$-L-rev and *tva C*$_{S-87}$-R-for/ *tva C*$_{S-87}$-R-rev, respectively, and HindIII-digested pYJ1004, leading to pYJ1008. This recombinant plasmid was then introduced into *S. sp.* NRRL S-87 by conjugation, and colonies that were apramycin-resistant at 30°C were identified as integrating mutants. These mutants were cultured with 800 μg/mL 5-fluorocytocine, resulting in 5-fluorocytocine-resistant isolates, which were subjected to PCR amplification for examination of the genotype of the *S. sp.* NRRL S-87 mutant strain YJ102, in which *tvaC*$_{S-87}$ was deleted in frame. The above CRISPR-Editing method was applied to the construction of the mutant strains YJ103, YJ104 and YJ105 for inactivating *tvaD*$_{S-87}$, *tvaE*$_{S-87}$ and *tvaF*$_{S-87}$, respectively, in *S. sp.* NRRL S-87.

For the inactivation of $lanL_{SL}$ in *S. laurentii*, a truncated $lanL_{SL}$ gene was constructed into pOJ260 through homologous recombination among the two 2-kb PCR products amplified from the genome of *S. laurentii* using the primer pairs $lanL_{SL}$-L-for/$lanL_{SL}$-L-rev and $lanL_{SL}$ -R-for/$lanL_{SL}$-R-rev, respectively, and EcoRI/BamHI-digested POJ260, leading to pYJ1017. This recombinant plasmid was introduced into *S. laurentii* by conjugation. The colonies that were apramycin-resistant at 30°C were identified as integrating mutants, in which a single-crossover homologous recombination event took place. These mutants were cultured for several rounds in the absence of apramycin, and the resulting apramycin-sensitive isolates were subjected to PCR amplification to examine the genotype of the *S. laurentii* mutant strain YJ113.

**Gene complementation.** For homologous complementation, the $tvaC_{S\text{-}87}$-containing fragment was amplified by PCR using primer pairs $tvaC_{S\text{-}87}$-C-for/$tvaC_{S\text{-}87}$-C-rev, and then cloned into the *PermE**-containing pSET152 derivative,[9] yielding pYJ1012, in which $tvaC_{S\text{-}87}$ was under the control of *PermE**, a constitutive promoter responsible for expressing the erythromycin-resistance gene *ermE* in *Saccharopolyspora erythraea*. pYJ1012 was then introduced into YJ102 (the △$tvaC_{S\text{-}87}$ mutant) by conjugation, generating the corresponding recombinant strain YJ106 in which $tvaC_{S\text{-}87}$ is expressed *in trans*. For complementation with a $tvaC_{S\text{-}87}$ double mutant, a two-step PCR amplification was conducted using pYJ1012 as the template, with the primer pairs $tvaC_{S\text{-}87}$-N241A-for/$tvaC_{S\text{-}87}$-N241A-rev and $tvaC_{S\text{-}87}$-D258A-for/$tvaC_{S\text{-}87}$-D258A-rev to produce pYJ1013, in which $tvaC_{S\text{-}87}$ was mutated to $tvaC_{S\text{-}87}$-*N241A/D258A*. After sequencing to conform the identity, this plasmid was introduced into YJ102 by conjugation, generating the corresponding recombinant strain YJ107 in which $tvaC_{S\text{-}87}$-N241A/D258A is expressed *in trans*.

For heterologous complementation, the *spaKC*-containing fragment was amplified by PCR using primer *spaKC*-for/*spaKC*-rev, and then cloned into the *PermE**-containing pSET152 derivative,[9] yielding pYJ1015, in which s*paKC* was under the control of *PermE**. pYJ1015 was then introduced into YJ102 (the △$tvaC_{S\text{-}87}$ mutant), YJ103 (the △$tvaD_{S\text{-}87}$ mutant) or YJ104 (the △$tvaE_{S\text{-}87}$ mutant) by conjugation, generating the corresponding recombinant strain YJ109, YJ110 or YJ111 in which *spaKC* is expressed *in trans*.

#### 1.3 Gene Cluster Engineering and Heterologous Expression

**Engineering of the mutant *tva*$_{S\text{-}87}$ cluster.** To enhance the expression of the *tva*$_{S\text{-}87}$ cluster (the △*tvaE*$_{S\text{-}87}$ mutant) in *S. sp.* NRRL S-87, the constitutive promoter *PermE** was inserted at the upstream of this cluster. The *PermE** promoter-containing fragment was firstly amplified by PCR from cosmid pYJ1016 using primer pairs *permE-for/2kb-rev*, and then cloned into EcoRI/HindIII-digested pOJ260 to obtain cosmid pYJ1014. pYJ1014 was introduced into strain YJ104 (the △*tvaE*$_{S\text{-}87}$ mutant) by conjugation to obtain the corresponding recombinant strain YJ108 (flowing a protocol described above).

**Heterologous Expression of the *tva*$_{S\text{-}87}$ cluster.** The 10-kb biosynthetic gene cluster of TVAs was assembled and cloned into the plasmid pJTU2554, yielding the recombinant plasmid pYJ1016 via homologous recombination among the two 5-kb PCR products amplified from the genome of *S. sp.* NRRL S-87 using the primer pairs YJ-1-for/YJ-1-rev and YJ-2-for/YJ-2-rev, respectively, the 0.5-kb PCR product amplified from the *PermE**-containing pSET152 derivative[9] using the primer pair PermE2-for/PermE2-rev, and the 5-kb PCR products amplified from pJTU2554 using the primer pair YJ-3-for/YJ-3-rev. The resulting plasmid pYJ1016, in which the biosynthetic genes in the *tva* cluster are under the control of *PermE**, was introduced into the thiostrepton-producing strain *S. laurentii* or YJ113 (the △*lanL*$_{SL}$ mutant) by conjugation, and colonies that were apramycin-resistant at 30°C were identified as the recombinant strain YJ112 or YJ114. The production and analysis of TVA-YJ-1 were conducted using methods described below.

#### 1.4 Production and Analysis of TVAs

**Culture and Fermentation.** The *S. sp.* NRRL S-87 wild type strain or its mutant derivative was spread on PS5 agar plates that contain the medium composed of 20 g of soluble starch, 5 g of pharmamedia, and 20 g of agar per liter (pH 7.0), and incubated at 28°C for sporulation and growth. Approximately 1 $cm^2$ of the sporulated agar of *S. sp.* NRRL S-87 was cut, chopped, and inoculated into 100 mL of the fermentation medium, which was composed of 25 g of glucose, 15 g of soybean meal, 2 g of dry yeast, and 4 g of $CaCO_3$ per liter (pH7.0).[10] After incubation at 28°C and 220 rpm for 96 hr, bacterial cells were collected by centrifugation (3000 × g for 45 min), and was stored at -80 °C before acetone extraction. The *S. laurentii* wild type strain or its mutant derivative was spread on ISP2 agar plates that contain the medium composed of 10 g of malt extract, 4 g of yeast extract, 4 g of glucose and 20 g of agar per liter (pH 7.0), and incubated at 28°C for sporulation and growth. Approximately 1 $cm^2$ of the sporulated agar of *S. laurentii* or its mutant derivative was cut, chopped, and inoculated into 100 mL of the fermentation medium, which was composed of 25 g of glucose, 15 g of soybean meal, 2 g of dry yeast, and 4 g of $CaCO_3$ per liter (pH7.0).[10] After incubation at 28°C and 220 rpm for 96 hr, bacterial cells were collected by centrifugation (3000 × g for 45 min), and was stored at -80 °C before methonal/acetone extraction.

**Product Examination.** 10 mL of each fermentation broth was centrifuged, and collected mycelia were soaked by 3 mL of acetone for 5 min. After centrifugation to remove the residue, the supernatant was evaporated before re-dissolution in 200 μL of methanol. The methanol sample was subjected to HPLC-MS analysis on an Agilent Zorbax column (SB-C18, 4.6×250 mm, 5 µm, Agilent Technologies Inc., USA) by gradient elution of solvent A ($H_2O$ + 0.1% formic acid) and solvent B (acetonitrile + 0.1% formic acid) with a flow rate of 1 mL/min over a 35 min period. For **Method I**, T = 0 min, 35% B; T = 5 min, 35% B; T = 20 min, 70% B; T = 25 min, 100% B; T = 30 min, 100% B; and T = 35 min, 35% B. For **Method II**, T = 0 min, 15% B; T = 20 min, 70% B; T = 25 min, 100% B; T = 28 min, 100% B; T = 30min, 15% B; and T = 35 min, 15% B. Related data were analyzed using Thermo Xcalibur software.

**Product Isolation.** 7 L of the $tva_{S\text{-}87}$-containing *S. laurentii* fermentation broth was centrifuged, and pelleted mycelial cake was extracted with 1.5 L of methanol three times. After filtration and concentration, the extract was loaded onto a silica gel column, which was pre-treated with dichloromethane–methanol (0, 5, 8, 10, 15 and 20% MeOH), and then eluted with the same solvent system (25 and 30% MeOH). After evaporation and re-dissolution in 3 mL of methanol, the crude sample was eluted on a Sephadex column (Ø = 1.5 cm, l = 200 cm, Pharmacia Co., Ltd., Sweden) with MeOH. The fraction containing TVAs was combined and concentrated to ~85 mg of dried extract. Further purification by RP-HPLC on an Xselect CSH C18 column (250 × 10 mm, 5 µm, Waters Technology Co., Ltd., USA) by gradient elution of solvent A ($H_2O$ + 0.1% formic acid) and solvent B (acetonitrile + 0.1% formic acid) with a flow rate of 3 mL/min over a 35 min period as follows: T = 0 min, 37% B; T = 5 min, 37% B; T = 20 min, 55% B; T = 25 min, 65% B; T = 30 min, 100% B; and T = 35 min, 37% B.

**1.5 Protein Expression and Purification.**

**Constructs for co-expression in *E. coli.*** The related $tva_{S\text{-}87}$ genes were amplified using corresponding primers. While the PCR product containing $tvaA_{S\text{-}87}$ was cloned into the plasmid pRSFduet-1, the products containing $tvaC_{S\text{-}87}$, $trxA$-$tvaD_{S\text{-}87}$, $tvaE_{S\text{-}87}$ and $tvaF_{S\text{-}87}$, were cloned individually into pETduet-1 and CDFDuet-1.

To prepare the variants of *tvaC*$_{S\text{-}87}$ and *tvaD*$_{S\text{-}87}$ by Site-specific mutagenesis, Rolling-cycle PCR amplification followed by subsequent DpnI digestion was performed according to the standard procedure of the QuickChange Site-Directed Mutagenesis Kit purchasedfrom Stratagene (GE Healthcare, USA) or Mut ExpressTM II (Vazyme Biotech Co. Ltd., China). Each mutation was confirmed by sequencing.

**Constructs for *in vitro* assays after purification from *E. coli*.** The related *tva*$_{S\text{-}87}$ genes were amplified using corresponding primers, and then cloned individually into pET37b(+) or pET28a(+) for the expression of the recombinant proteins TvaE$_{S\text{-}87}$, TvaF$_{S\text{-}87}$ and the LP sequence of precursor peptide TvaA$_{S\text{-}87}$.

**Protein expression.** *E. coli* BL21(DE3) served as a general host for heterologous expression. The culture of each recombinant *E. coli* strain was incubated in Luria-Bertani (LB) medium (5 g of yeast extract, 10 g of tryptone and 10 g of NaCl per liter) containing 50 μg/mL kanamycin, 100 μg/mL ampicillin and 50 μg/mL streptomycin, at 37°C and 250 rpm until the cell density reached 0.6-0.8 at $OD_{600}$. Protein expression was induced by the addition of isopropyl-β-D-thiogalactopyranoside (IPTG) to a final concentration of 0.1-0.3 mM, followed by further incubation for 25-30 hr at 25°C or 16°C. The cells were harvested by centrifugation at 3000 × g for 20 min, flash-frozen and then stored at -80°C.

**Protein purification.** *E. coli* cells were re-suspended in lysis buffer (137 mM NaCl, 2.7 mM KCl, 10 mM $Na_2HPO_4$, 1.8 mM $KH_2PO_4$, 10% glycerol and 5 mM imidazole, pH 8.0). After disruption by FB-110X Low Temperature Ultra-Pressure Continuous Flow Cell Disrupter (Shanghai Litu Mechanical Equipment Engineering Co., Ltd, China), soluble fractions were collected by centrifugation. Recombinant proteins that contain a 6xHis-tag were purified on a HisTrap HP column (GE Healthcare, USA), which was pre-treated with 10 column volumes (CVs) of lysis buffer followed by 10 CVs of wash buffer (137 mM NaCl, 2.7 mM KCl, 10 mM $Na_2HPO_4$, 1.8 mM $KH_2PO_4$, 10% glycerol and 40 mM imidazole, pH 7.4), using elution buffer (137 mM NaCl, 2.7 mM KCl, 10 mM $Na_2HPO_4$, 1.8 mM $KH_2PO_4$, 10% glycerol and 250 mM imidazole, pH 7.4). Desired protein fractions were concentrated (to 500 μM-1 mM) using Amicon® Ultra-15 Centrifugal Filter Devices (MILLIPORE, USA) and desalted using a PD-10 Desalting Column (GE Healthcare, USA) according to the manufacturer's protocols, and then quantified in concentration by Bradford assay using bovine serum albumin as the standard.

The purity of recombinant proteins was determined by sodium dodecyl sulfate polyacrylamide gel electrophoresis (SDS-PAGE). For the determination of the flavin cofactor associated with $TvaF_{S\text{-}87}$, The UV spectra of recombinant proteins were recorded at a concentration of 23 mg/mL on a DeNovix DS-11 UV/Vis spectrophotometer (DeNovix Inc. Wilmington, DE 19810 USA). Each protein solution was incubated at 100˚C for 5 min for denaturation and then subjected to HR-ESI-MS analysis on a 6230B Accurate Mass TOF LC/MS System (Agilent Technologies Inc., USA) to examine the presence of FMN (with the *m/z* $[M+H]^+$ mode calcd. 457.1124; observed 457.1139) in purified $TvaF_{S\text{-}87}$.

For *N*-terminal tag removal, the protease-encoding gene *3c* was synthesized by Genscript Biotech (Nanjing, China), and cloned into pET28a(+) for the expression of the recombinant 3C protein. For the removal of Small Ubiquitin-related Modification (SUMO)-tag, proteolysis was conducted at 30°C 15-30 min in 30 μL of the reaction mixture that contained 15 μM SUMO Protease (Ulp-1), 100 μM SUMO-tagged $TvaA_{S\text{-}87}$ or its derivative, 50 mM NaCl, 100 mM HEPES (pH 7.5). Purified products containing precursor peptide $TvaA_{S\text{-}87}$ and/or its variants were subjected to HPLC-HR-MS and HR-MS/MS analyses under conditions as described below. The derivatization of (ene)thiol-containing $TvaA_{S\text{-}87}$ and/or its variants was conducted using a method described below.

To prepare mono-dehydrated **1** for *in vitro* assays, the above product resulting from the co-expression of *$tvaA_{S\text{-}87}$* with *$tvaCD_{S\text{-}87}$* in *E. coli* was applied to further purification on a MonoSpin C18 column (GL Sciences Inc., Japan). This column was equilibrated with acetonitrile (ACN) and the water containing 0.1% formic acid. After washing with the ACN solution (5%) containing 0.1% formic acid to remove polar contaminants and then with the ACN solution (35%) containing 0.1% formic acid, the sample was eluted with the ACN solution (60%) containing 0.1% formic acid. The collection was freeze-dried using a freeze dryer (Martin Christ Inc., Germany). After lyophilization, the collection was subject to HPLC-HR-MS and HR-MS/MS analyses under conditions as described below.

To concentrate **3** for phosphorylation examination, the above product resulting from the co-expression of *$tvaA_{S\text{-}87}$* with wild-type *$tvaC_{S\text{-}87}$* and mutant *$tvaD_{S\text{-}87}$-H24A/R26A* in *E. coli* was applied to further purification on a High-Select™ $TiO_2$ Phosphopeptide Enrichment kit (Thermo Fisher Scientific Co. Ltd., USA) by following the manufacturer's procedure. The collection was subjected to HPLC-HR-MS and HR-MS/MS analysis under conditions as described below.

#### 1.6 *In vitro* Assays of $TvaE_{S\text{-}87}$ and $TvaF_{S\text{-}87}$ Activities

***In vitro* Examination of AviCys formation.** Conversions were conducted at 30°C for 2 hr in 30 μL of the reaction mixture that contained 10-100 μM **1**, 20 μM $TvaF_{S-87}$, 20 μM $TvaE_{S-87,}$ or both $TvaE_{S-87}$ and 20 μM $TvaE_{S-87}$ along with 50 mM Tris-HCl (pH 8.0), 1 mM TCEP, 5 μM FMN, 10 mM KCl, 10 mM $MgCl_2$ and 5 μM 3C. Reactions were quenched by adding equal volumes of acetonitrile, and after centrifugation, reaction mixtures were subjected to HPLC-HR-MS and HR-MS/MS analyses. For (ene)thiol derivatization, 20 μL of each quenched reaction mixture was treated with iodoacetamide (IAA) in dark at room temperature for 15-30 min before 6% (v/v) formic acid was added to eliminate excessive IAA and prevent side reactions.

**HPLC-HR-MS and HR-MS/MS analyses**. An Agilent ZORBAX column (300SB-C18, 2.1 mm × 100 mm, 3.5 μm, Agilent Technologies Inc., USA) or an Agilent 300Extend-C18 column (2.1 mm × 100 mm, 3.5 μm, Agilent Technologies Inc., USA) was used on an UltiMate 3000 UHPLC system coupled to a Thermo Scientific Q Exactive Plus Orbitrap mass spectrometer. Gradient elution was conducted using solvent A ($H_2O$ + 0.1% formic acid) and solvent B (ACN + 0.1% formic acid) with a flow rate of 0.3 mL/min over a 43 min period as follows: T = 0 min, 10% B; T = 2 min, 10% B; T = 20 min, 30% B; T = 25 min, 60% B; T = 30 min, 100% B; T = 35 min, 100% B; T = 38 min, 10% B; and T = 43 min, 10% B. Unless otherwise stated, ESI-MS was performed in positive ion mode, with a spray voltage of 3800 V, a capillary temperature of 375°C, aux gas heater temperature 350°C and an S-lens level 60. Full MS was examined at a resolution of 70,000 (AGC target 2e5, maximum IT 50 ms, range 300–1000 or 400-1800 m/z). Parallel reaction monitoring (PRM) or data-dependent MS/MS was performed at a resolution of 35,000 (AGC target between 1e5 and 1e6, maximum IT between 100 ms and 250 ms, isolation windows in the range of 1.0 to 2.0 m/z) using a stepped NCE of 18, 20 and 28 or an NCE of 25. Scan ranges, inclusion lists, charge exclusions, and dynamic exclusions were adjusted as needed.

#### 1.7 Measurement of Protein-Protein Interactions

**MicroScale Thermophoresis (MST) assay.** The affinity of the purified $TvaF_{S-87}$ and $TvaA_{LP(S-87)}$ to $TvaE_{S-87}$ was measured using the Monolith NT.115Pico (Nanotemper Technologies). $TvaE_{S-87}$ were site-specifically fluorescently labelled according to the protein labeling procedure described in The Monolith His-tag Labeling Kit RED-tris-NTA 2$^{nd}$ Generation. Then, the labelled proteins were premixed with 0.07% Tween 20 before the binding assay. The same volume (10 μL) labelled $TvaE_{S-87}$ was mixed thoroughly with same volume unlabeled ligands ($TvaF_{S-87}$ and $TvaA_{LP(S-87)}$, respectively) of 16 different serial concentrations in storage buffer supplemented with 0.07% Tween 20. The mixture was then loaded into 16 silica capillaries and measured at 25°C by using standard method set in Monolith NT.115$^{pico}$. Each assay was repeated three times and data analyses were performed using Origin 2020b software and Nanotemper analysis software.

### 2. Supplementary Figures

**Figure S1.** Purified recombinant enzymes analysis. (**A**) The TvaA$_{LP(S\text{-}87)}$ (red rectangular) was purified in Superdex 75 16/60 column coupled with an AKTA pure system (GE Healthcare) at 8°C (λ = 220 nm). (**B**) Coomassie-stained SDS-PAGE analysis of TvaA$_{LP(S\text{-}87)}$ (Line 1, red rectangular), TvaE$_{S\text{-}87}$ (Line 2), TvaF$_{S\text{-}87}$ (Line 3) and protein standard (Line M).

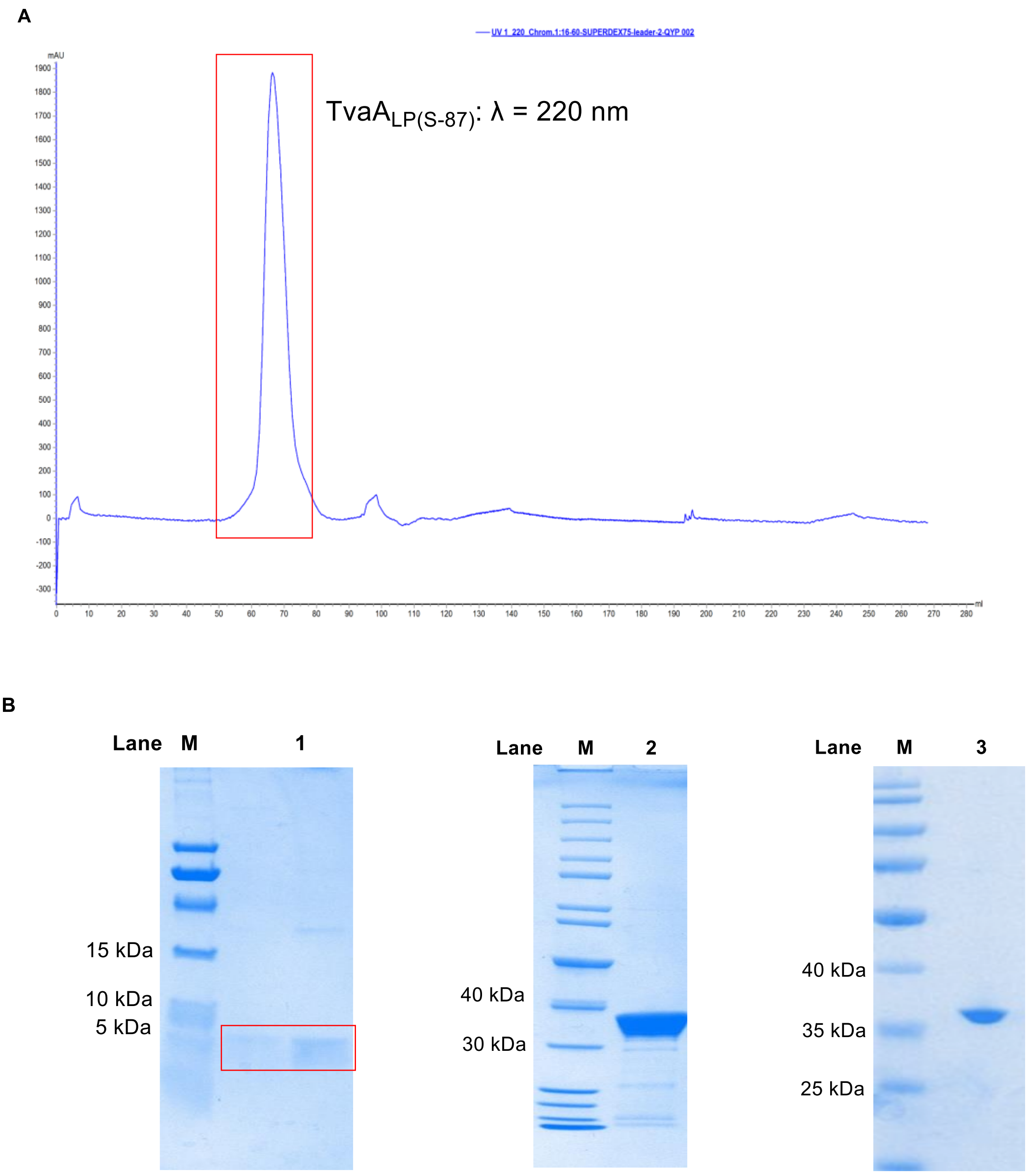

**Figure S2.** Alignment of the $tva_{S\text{-}87}$ cluster from *S. sp.* NRRL S-87 with the prototypical gene cluster identified from *S.olivoviridis* NA05001.[10]

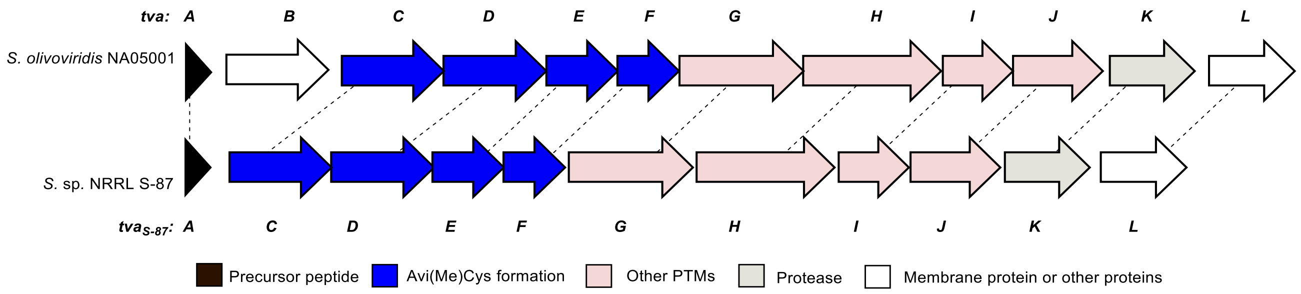

**Figure S3.** Characterization of the production of TVA-YJ-1. (**A**) UV spectrum of TVA-YJ-1. (**B**) COSY, HMBC and NOESY correlations for the structural characterization of TVA-YJ-1. (**C**) HR-MS and HR-MS/MS spectra of TVA-YJ-1.

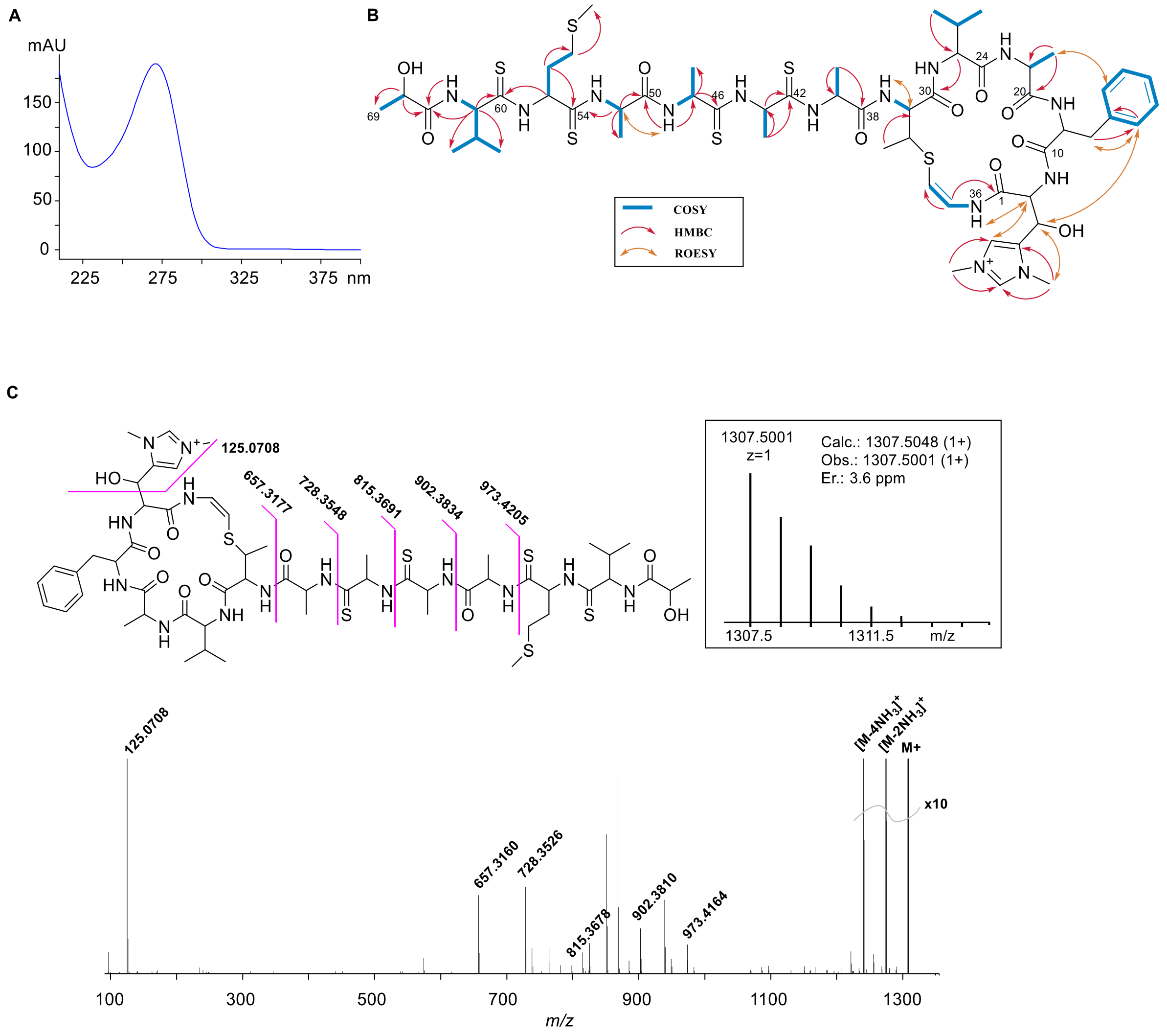

**Figure S4.** Sequence alignment of TvaC$_{S-87}$, TvaE$_{S-87}$, LxmX, LxmK,[11] MicKC (kinase domain),[12] and SpaKC[13] (kinase domain) with known APH family proteins 1J7I_A, 2BKK_A, 3Q2J_A, 4DCA_A, 4FEX_A and 6IY9_A. The conserved residues D236, N241 and D258 are indicated by arrows.

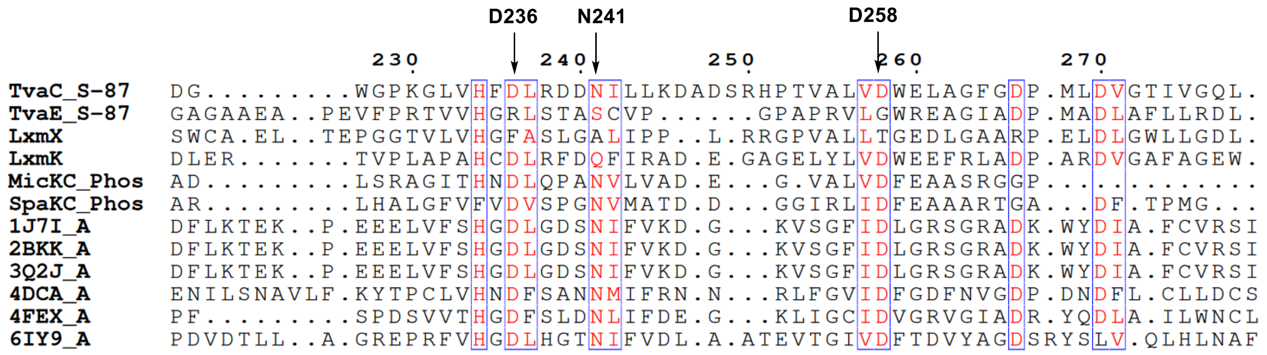

**Figure S5.** Structural homology shared by various characterized proteins with TvaD$_{S-87}$ based on the prediction of its secondary structure using HHpred program (https://toolkit.tuebingen.mpg.de/tools/hhpred).[14] Related phosphothreonine lyase OspF-catalyzed reaction is shown below.

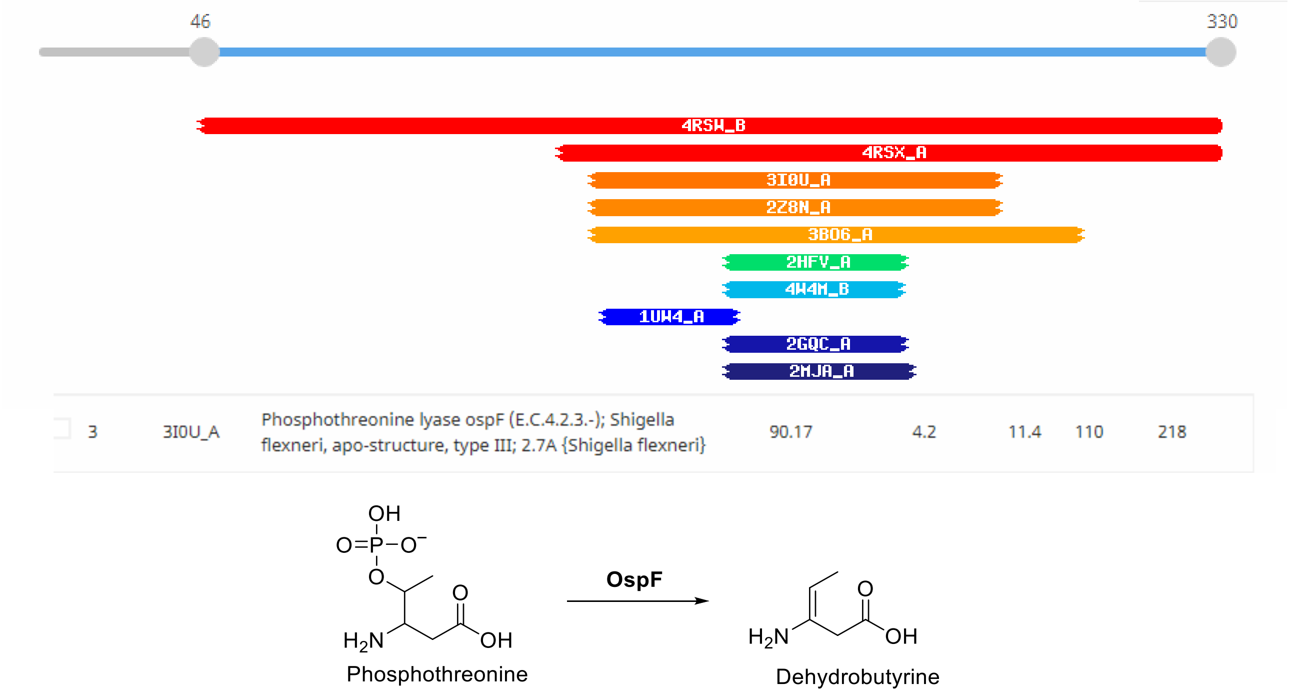

**Figure S6.** Characterization of TvaA$_{S-87}$. (**A**) HPLC-HR-MS analysis ($C_{384}H_{612}N_{100}O_{139}S_5$ $[M+6H]^{6+}$ m/z: calculated 1502.3822, observed 1502.3879 and error 3.8 ppm). (**B**) HR-MS/MS analysis. The HCD fragments and the MS/MS spectrum are shown.

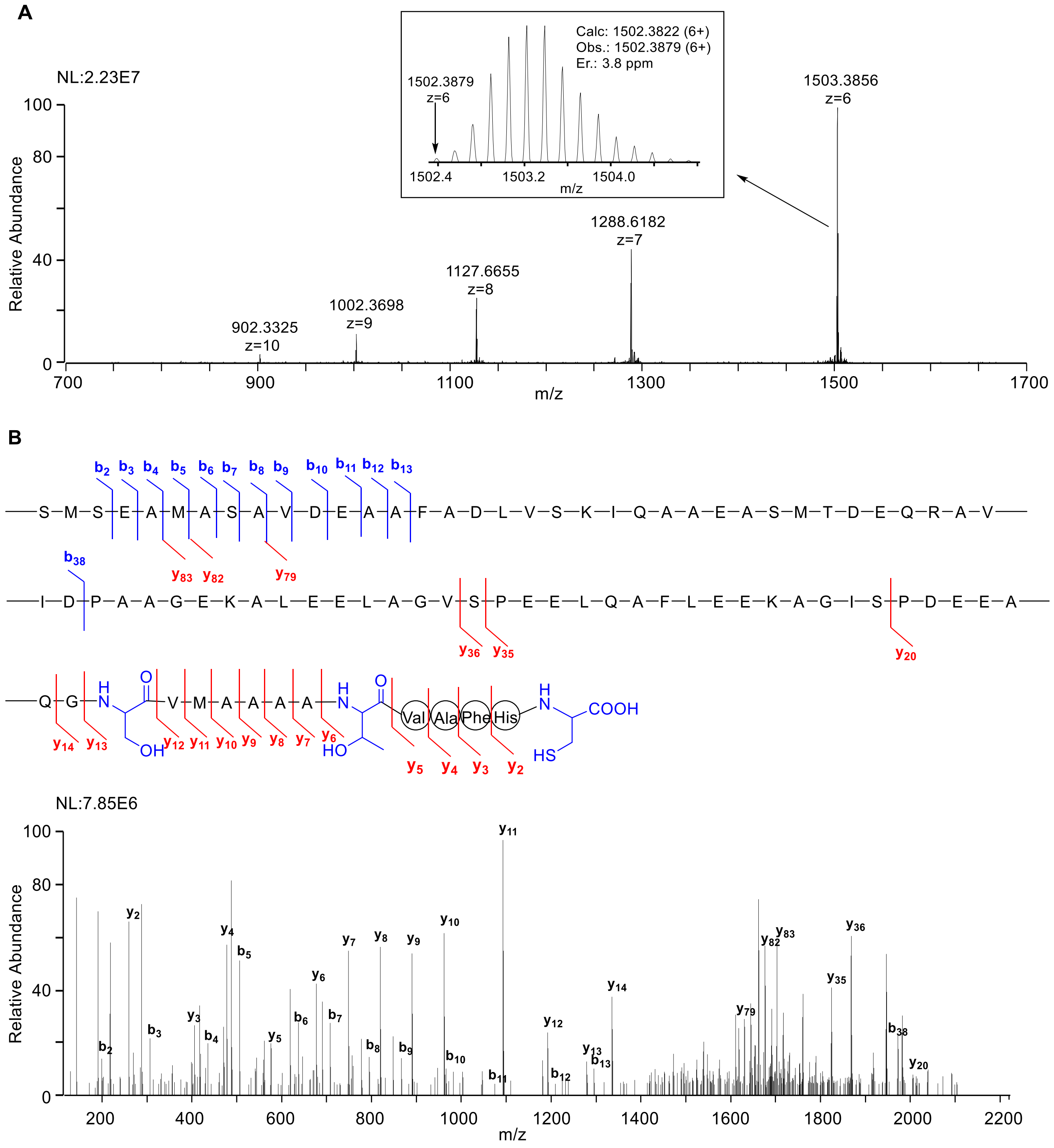

| Ions | Calc. | Obs. | Er. (ppm) | Ions | Calc. | Obs. | Er. (ppm) |
|---|---|---|---|---|---|---|---|
| **$b_2^+$** | 219.0803 | 219.0798 | 2.3 | **$y_2^+$** | 259.0854 | 259.0860 | 2.3 |
| **$b_3^+$** | 306.1124 | 306.1118 | 1.9 | **$y_3^+$** | 406.1538 | 406.1543 | 1.2 |
| **$b_4^+$** | 435.1549 | 435.1545 | 0.9 | **$y_4^+$** | 477.1909 | 477.1917 | 1.7 |
| **$b_5^+$** | 506.1921 | 506.1918 | 0.6 | **$y_5^+$** | 576.2593 | 576.2599 | 1.0 |
| **$b_6^+$** | 637.2325 | 637.2325 | 0.0 | **$y_6^+$** | 677.3070 | 677.3078 | 1.2 |
| **$b_7^+$** | 708.2697 | 708.2694 | 0.4 | **$y_7^+$** | 748.3441 | 748.3446 | 0.7 |
| **$b_8^+$** | 795.3017 | 795.3008 | 1.1 | **$y_8^+$** | 819.3812 | 819.3821 | 1.1 |
| **$b_9^+$** | 866.3388 | 866.3406 | 2.1 | **$y_9^+$** | 890.4183 | 890.4186 | 0.3 |
| **$b_{10}^+$** | 965.4072 | 965.4072 | 0.0 | **$y_{10}^+$** | 961.4554 | 961.4562 | 0.8 |
| **$b_{11}^+$** | 1080.4342 | 1080.4344 | 0.2 | **$y_{11}^+$** | 1092.4960 | 1092.4966 | 0.5 |
| **$b_{12}^+$** | 1209.4767 | 1209.4755 | 1.0 | **$y_{12}^+$** | 1191.5643 | 1191.5657 | 1.2 |
| **$b_{13}^+$** | 1280.5138 | 1280.5103 | 2.7 | **$y_{13}^+$** | 1278.5963 | 1278.5972 | 0.7 |
| **$b_{38}^{2+}$** | 1970.9164 | 1970.9158 | 0.3 | **$y_{14}^+$** | 1335.6178 | 1335.6189 | 0.8 |
| | | | | **$y_{20}^+$** | 2004.8784 | 2004.8729 | 2.7 |
| | | | | **$y_{35}^{2+}$** | 1823.8553 | 1823.8549 | 0.2 |
| | | | | **$y_{36}^{2+}$** | 1867.3713 | 1867.3730 | 0.9 |
| | | | | **$y_{79}^{5+}$** | 1629.5913 | 1629.5789 | 7.6 |
| | | | | **$y_{82}^{5+}$** | 1675.4125 | 1675.4177 | 3.1 |
| | | | | **$y_{83}^{5+}$** | 1701.6217 | 1701.6388 | 10.0 |

**Figure S7.** Characterization of **1**. (**A**) HPLC-HR-MS analysis ($C_{384}H_{610}N_{100}O_{138}S_5$ $[M+6H]^{6+}$ m/z: calculated 1499.3805, observed 1499.3906 and error 6.7 ppm). (**B**) HR-MS/MS analysis. The HCD fragments and the MS/MS spectrum are shown.

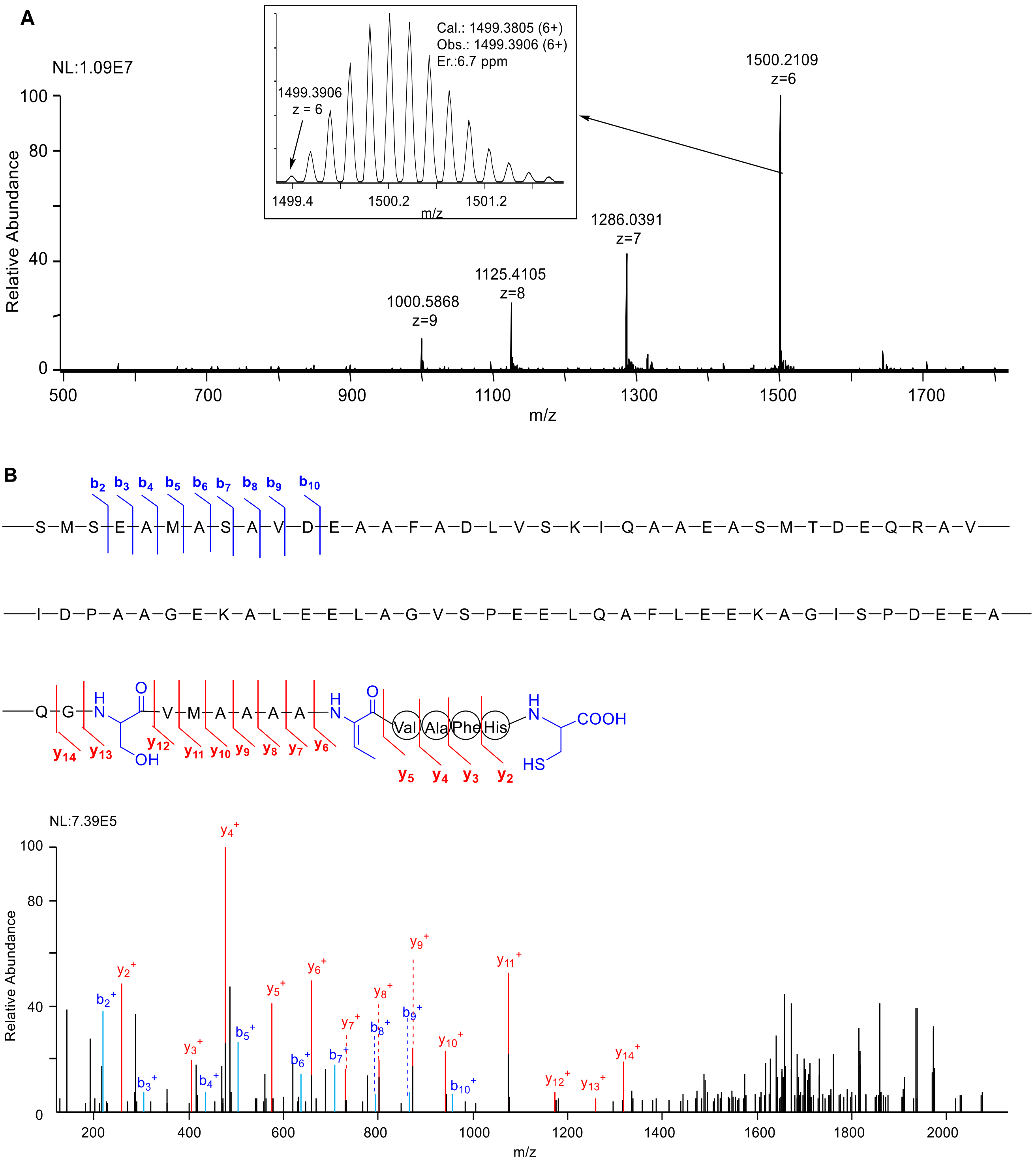

| Ions | Calc. | Obs. | Er. (ppm) | Ions | Calc. | Obs. | Er. (ppm) |
|---|---|---|---|---|---|---|---|
| **$b_2^+$** | 219.0803 | 219.0791 | 5.5 | **$y_2^+$** | 259.0854 | 259.0851 | 1.2 |
| **$b_3^+$** | 306.1124 | 306.1107 | 5.5 | **$y_{3+}$** | 406.1537 | 406.1527 | 2.5 |
| **$b_4^+$** | 435.1549 | 435.1528 | 4.8 | **$y_4^+$** | 477.1909 | 477.1899 | 2.1 |
| **$b_5^+$** | 506.1921 | 506.1914 | 1.4 | **$y_5^+$** | 576.2593 | 576.2585 | 1.4 |
| **$b_6^+$** | 637.2325 | 637.2311 | 2.2 | **$y_6^+$** | 659.2964 | 659.2941 | 3.5 |
| **$b_7^+$** | 708.2697 | 708.2665 | 4.5 | **$y_7^+$** | 730.3336 | 730.3353 | 2.3 |
| **$b_8^+$** | 795.3017 | 795.2980 | 4.7 | **$y_8^+$** | 801.3707 | 801.3706 | 0.1 |
| **$b_9^+$** | 866.3388 | 866.3345 | 5.0 | **$y_9^+$** | 872.4078 | 872.4044 | 3.9 |
| **$b_{10}^+$** | 965.4072 | 965.4000 | 7.4 | **$y_{10}^+$** | 943.4449 | 943.4402 | 5.0 |
| | | | | **$y_{11}^+$** | 1074.4854 | 1074.4819 | 3.3 |
| | | | | **$y_{12}^+$** | 1173.5538 | 1173.5494 | 3.7 |
| | | | | **$y_{13}^+$** | 1260.5858 | 1260.5825 | 2.6 |
| | | | | **$y_{14}^+$** | 1317.6073 | 1317.6047 | 2.0 |

**Figure S8.** Characterization of **2**. (**A**) HPLC-HR-MS analysis. ($C_{384}H_{608}N_{100}O_{137}S_5$ $[M+6H]^{6+}$ m/z: calculated 1496.3787, observed 1496.3881 and error 6.2 ppm). (**B**) HR-MS/MS analysis. The HCD fragments and the MS/MS spectrum are shown.

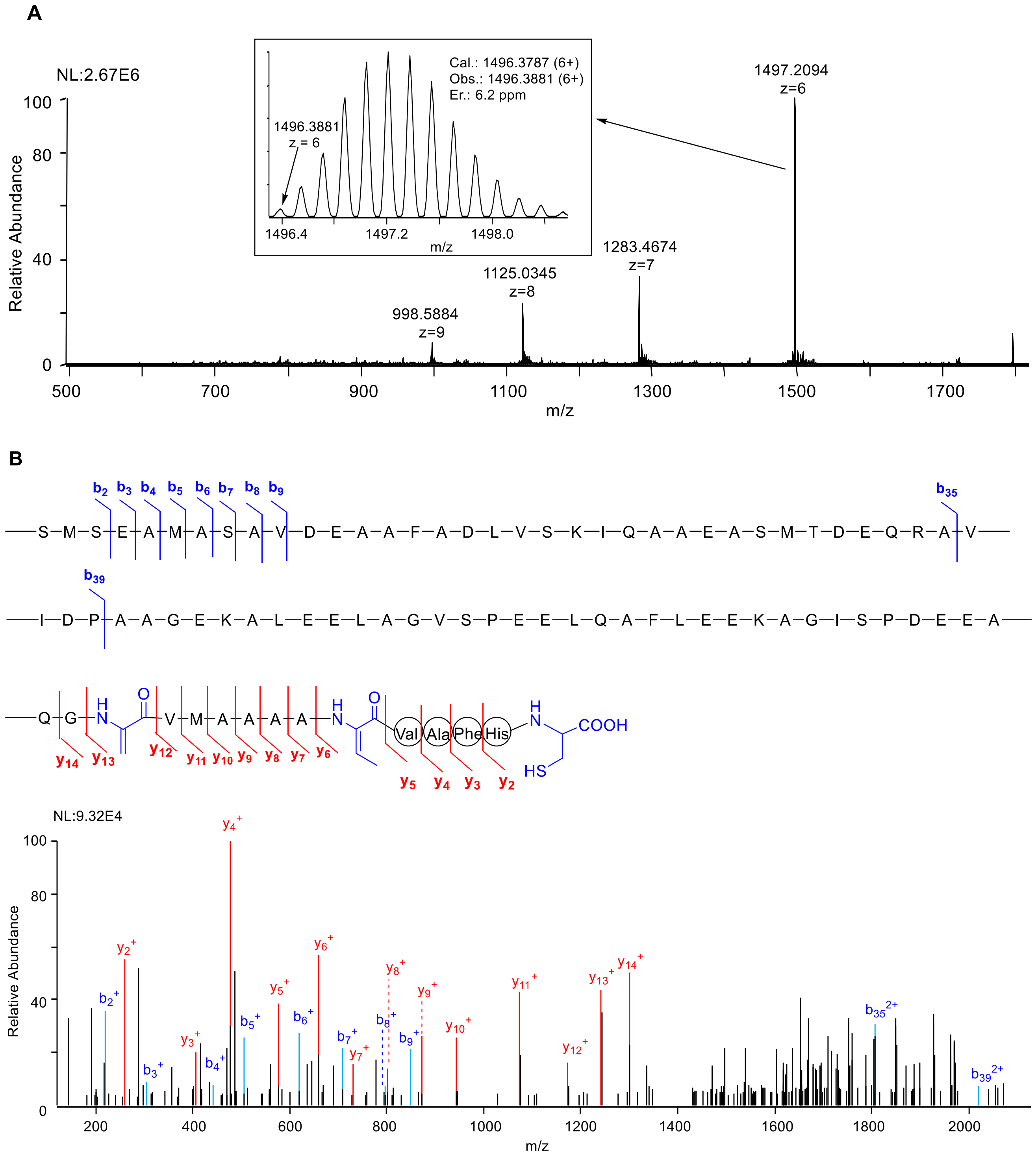

| Ions | Calc. | Obs. | Er. (ppm) | Ions | Calc. | Obs. | Er. (ppm) |
|---|---|---|---|---|---|---|---|
| **$b_2^+$** | 219.0803 | 219.0791 | 5.5 | **$y_2^+$** | 259.0854 | 259.0850 | 1.5 |
| **$b_3^+$** | 306.1124 | 306.1099 | 8.2 | **$y_{3+}$** | 406.1537 | 406.1532 | 1.2 |
| **$b_4^+$** | 435.1549 | 435.1527 | 5.0 | **$y_4^+$** | 477.1909 | 477.1901 | 1.7 |
| **$b_5^+$** | 506.1921 | 506.1896 | 4.9 | **$y_5^+$** | 576.2593 | 576.2589 | 0.7 |
| **$b_6^+$** | 637.2325 | 637.2328 | 0.5 | **$y_6^+$** | 659.2964 | 659.2951 | 2.0 |
| **$b_7^+$** | 708.2697 | 708.2667 | 4.2 | **$y_7^+$** | 730.3336 | 730.3333 | 0.4 |
| **$b_8^+$** | 795.3017 | 795.2976 | 5.2 | **$y_8^+$** | 801.3707 | 801.3690 | 2.1 |
| **$b_9^+$** | 866.3388 | 866.3376 | 1.4 | **$y_9^+$** | 872.4078 | 872.4028 | 5.7 |
| **$b_{35}^{2+}$** | 1807.3267 | 1807.3388 | 6.7 | **$y_{10}^+$** | 943.4449 | 943.4387 | 6.6 |
| **$b_{39}^{2+}$** | 2019.4427 | 2019.4587 | 8.0 | **$y_{11}^+$** | 1074.4854 | 1074.4840 | 1.3 |
| | | | | **$y_{12}^+$** | 1173.5538 | 1173.5542 | 0.3 |
| | | | | **$y_{13}^+$** | 1242.5753 | 1242.5721 | 2.6 |
| | | | | **$y_{14}^+$** | 1299.5967 | 1299.5970 | 0.2 |

**Figure S9.** Characterization of **3**. (**A**) HPLC-HR-MS analysis. ($C_{384}H_{613}N_{100}O_{142}PS_5$ $[M+6]^{6+}$ m/z: calculated 1515.7100 observed 1515.7150 and error 3.3 ppm). (**B**) HR-MS/MS analysis. The HCD fragments and the MS/MS spectrum are shown.

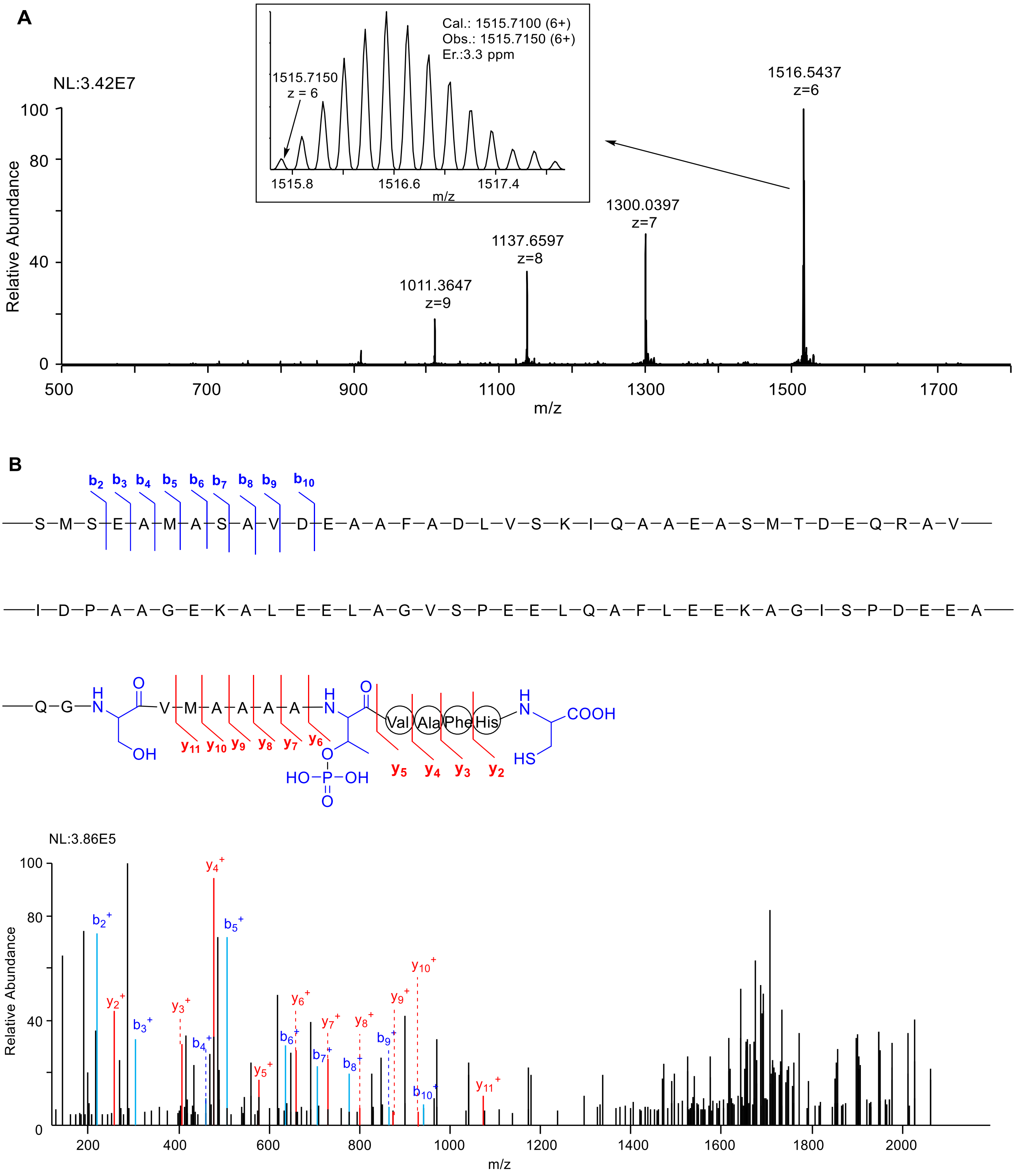

| Ions | Calc. | Obs. | Er. (ppm) | Ions | Calc. | Obs. | Er. (ppm) |
|---|---|---|---|---|---|---|---|
| **$b_2^+$** | 219.0803 | 219.0793 | 4.6 | **$y_2^+$** | 259.0864 | 259.0854 | 3.9 |
| **$b_3^+$** | 306.1124 | 306.1112 | 3.9 | **$y_{3+}$** | 406.1548 | 406.1548 | 0.0 |
| **$b_4^+$** | 435.1549 | 435.1547 | 0.6 | **$y_4^+$** | 477.1920 | 477.1907 | 2.7 |
| **$b_5^+$** | 506.1921 | 506.1903 | 3.6 | **$y_5^+$** | 576.2604 | 576.2611 | 1.2 |
| **$b_6^+$** | 637.2325 | 637.2323 | 0.3 | **$y_6^+$-$H_2PO_4$** | 659.2975 | 659.2973 | 0.3 |
| **$b_7^+$** | 708.2697 | 708.2706 | 1.3 | **$y_7^+$** | 730.3346 | 730.3325 | 2.9 |
| **$b_8^+$** | 795.3017 | 795.3024 | 0.9 | **$y_8^+$** | 801.3717 | 801.3718 | 0.1 |
| **$b_9^+$** | 866.3388 | 866.3384 | 0.5 | **$y_9^+$** | 872.4088 | 872.4051 | 4.2 |
| **$b_{10}^+$** | 965.4072 | 965.4044 | 2.9 | **$y_{10}^+$** | 943.4459 | 943.4477 | 1.9 |
| | | | | **$y_{11}^+$** | 1074.4864 | 1074.4897 | 3.1 |

**Figure S10.** Characterization of **4**. (**A**) HPLC-HR-MS analysis. ($C_{383}H_{610}N_{100}O_{137}S_5$ $[M+6]^{6+}$ m/z: calculated 1494.7147 observed 1494.7272 and error 8.3 ppm). (**B**) HR-MS/MS analysis. The HCD fragments and the MS/MS spectrum are shown.

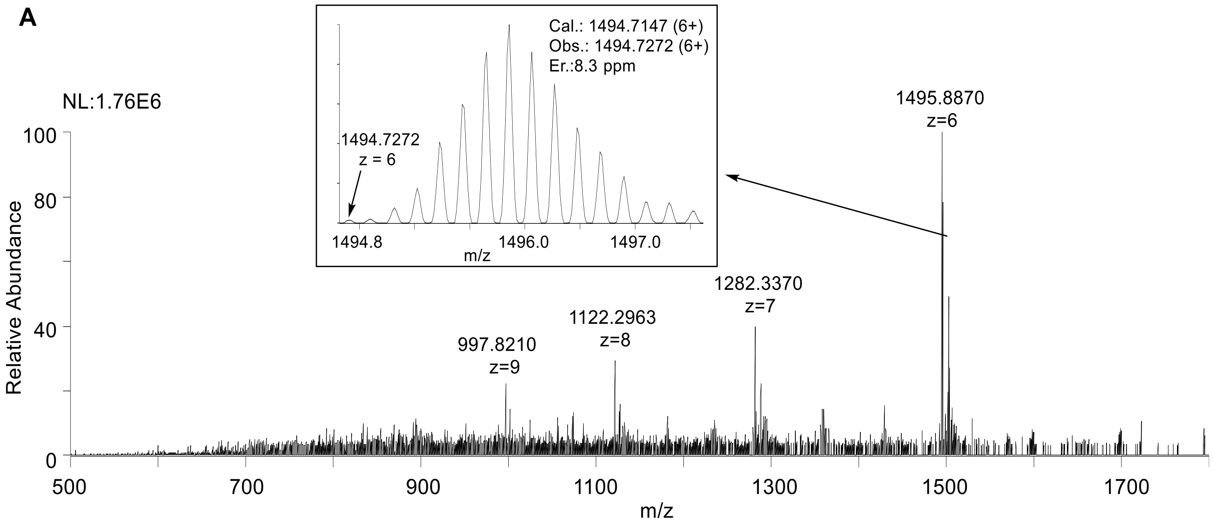


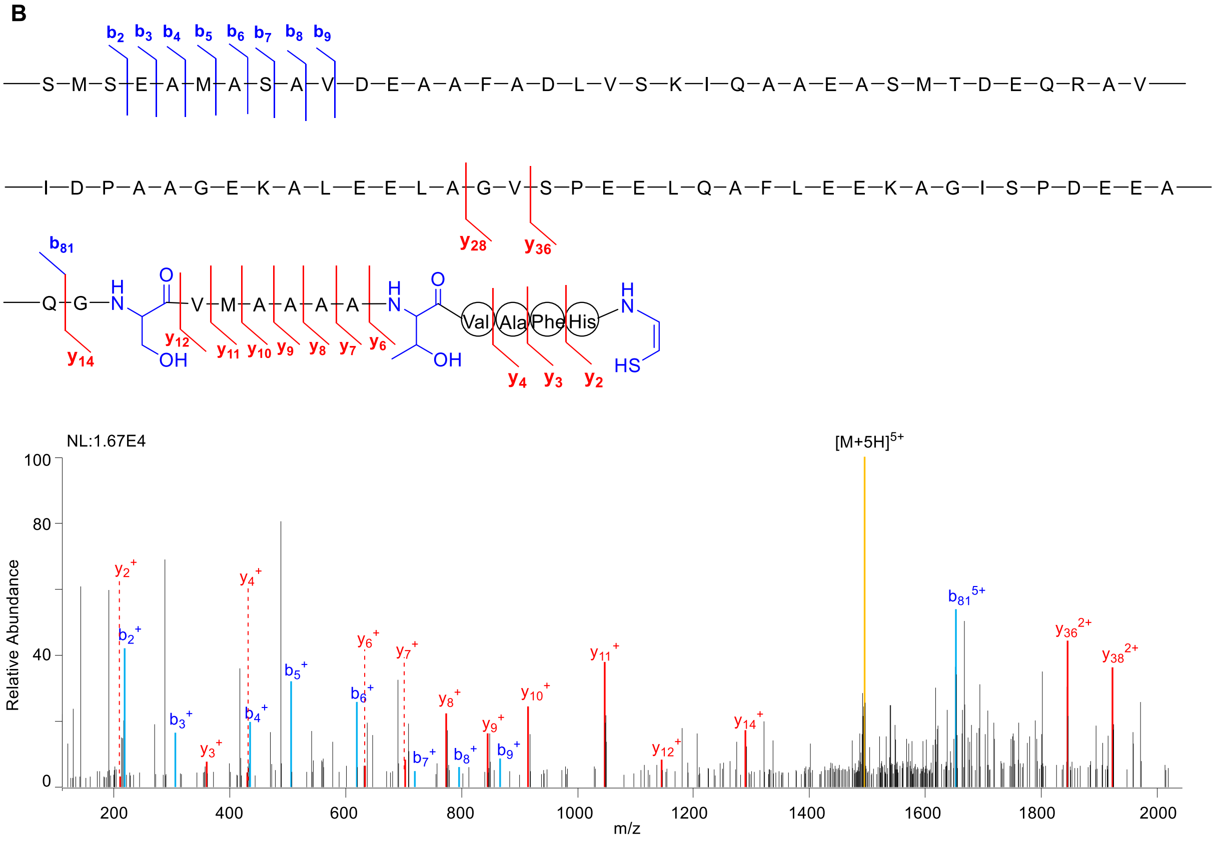

| Ions | Calc. | Obs. | Er. (ppm) | Ions | Calc. | Obs. | Er. (ppm) |
|---|---|---|---|---|---|---|---|
| $b_2$ | 219.0803 | 219.0797 | 2.7 | $y_2$ | 213.0810 | 213.0804 | 2.8 |
| $b_3$ | 306.1124 | 306.1119 | 1.6 | $y_3$ | 360.1494 | 360.1487 | 1.9 |
| $b_4$ | 435.1549 | 435.1552 | 0.7 | $y_4$ | 431.1865 | 431.1852 | 3.0 |
| $b_5$ | 506.1921 | 506.1919 | 0.4 | $y_6$ | 631.3026 | 631.3033 | 1.1 |
| $b_6$ | 637.2325 | 637.2328 | 0.5 | $y_7$ | 702.3397 | 702.3392 | 0.7 |
| $b_7$ | 708.2697 | 708.2703 | 0.8 | $y_8$ | 773.3768 | 773.3771 | 0.4 |
| $b_8$ | 795.3017 | 795.3024 | 0.9 | $y_9$ | 844.4139 | 844.4152 | 1.5 |
| $b_9$ | 866.3388 | 866.3392 | 0.5 | $y_{10}$ | 915.4510 | 915.4500 | 1.1 |
| $b_{81}^{5+}$ | 1653.1902 | 1653.2036 | 8.1 | $y_{11}$ | 1046.4915 | 1046.4918 | 0.3 |
| | | | | $y_{12}$ | 1145.5599 | 1145.5620 | 1.8 |
| | | | | $y_{14}$ | 1289.6134 | 1289.6133 | 0.1 |
| | | | | $y_{36}^{2+}$ | 1844.3694 | 1844.3816 | 6.6 |
| | | | | $y_{38}^{2+}$ | 1922.4144 | 1922.4240 | 5.0 |

**Figure S11.** Characterization of **4a**. (**A**) HPLC-HR-MS analysis. ($C_{383}H_{612}N_{100}O_{137}S_5$ $[M+6]^{6+}$ m/z: calculated 1495.0506 observed 1495.0551 and error 3.0 ppm). (**B**) HR-MS/MS analysis. The HCD fragments and the MS/MS spectrum are shown.

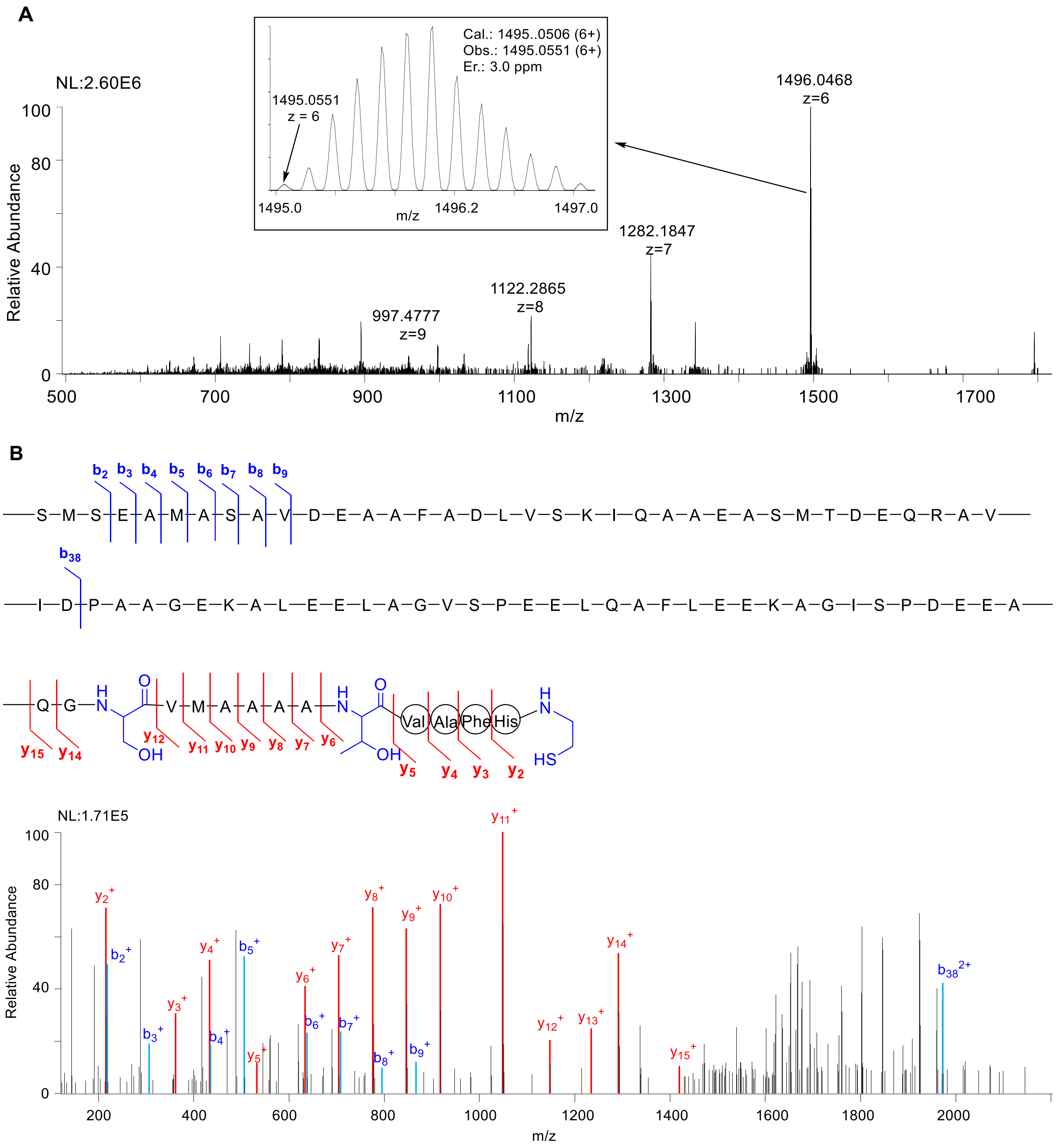

| Ions | Calc. | Obs. | Er. (ppm) | Ions | Calc. | Obs. | Er. (ppm) |
|---|---|---|---|---|---|---|---|
| $b_2$ | 219.0803 | 219.0795 | 3.7 | $y_2$ | 215.0956 | 215.0964 | 3.7 |
| $b_3$ | 306.1124 | 306.1118 | 2.0 | $y_3$ | 362.1640 | 362.1641 | 0.3 |
| $b_4$ | 435.1549 | 435.1540 | 2.1 | $y_4$ | 433.2011 | 433.2013 | 0.5 |
| $b_5$ | 506.1921 | 506.1963 | 8.3 | $y_5$ | 532.2695 | 532.2699 | 0.8 |
| $b_6$ | 637.2325 | 637.2321 | 0.6 | $y_6$ | 633.3172 | 633.3173 | 0.2 |
| $b_7$ | 708.2697 | 708.2700 | 0.4 | $y_7$ | 704.3543 | 704.3561 | 2.6 |
| $b_8$ | 795.3017 | 795.3038 | 2.6 | $y_8$ | 775.3914 | 775.3915 | 0.1 |
| $b_9$ | 866.3388 | 866.3367 | 2.4 | $y_9$ | 846.4269 | 846.4286 | 2.1 |
| $b_{38}^{2+}$ | 1970.9164 | 1970.9052 | 5.7 | $y_{10}$ | 917.4656 | 917.4653 | 0.3 |
| | | | | $y_{11}$ | 1048.5061 | 1048.5065 | 0.4 |
| | | | | $y_{12}$ | 1147.5745 | 1147.5736 | 0.8 |
| | | | | $y_{13}$ | 1234.6066 | 1234.6111 | 3.6 |
| | | | | $y_{14}$ | 1291.6280 | 1291.6274 | 0.5 |
| | | | | $y_{15}$ | 1419.6866 | 1419.6971 | 7.4 |

**Figure S12.** Characterization of **4b**. (**A**) HPLC-HR-MS analysis. ($C_{383}H_{610}N_{100}O_{138}S_4$ $[M+6]^{6+}$ m/z: calculated 1492.0518 observed 1492.0566 and error 3.2 ppm). (**B**) HR-MS/MS analysis. The HCD fragments and the MS/MS spectrum are shown.

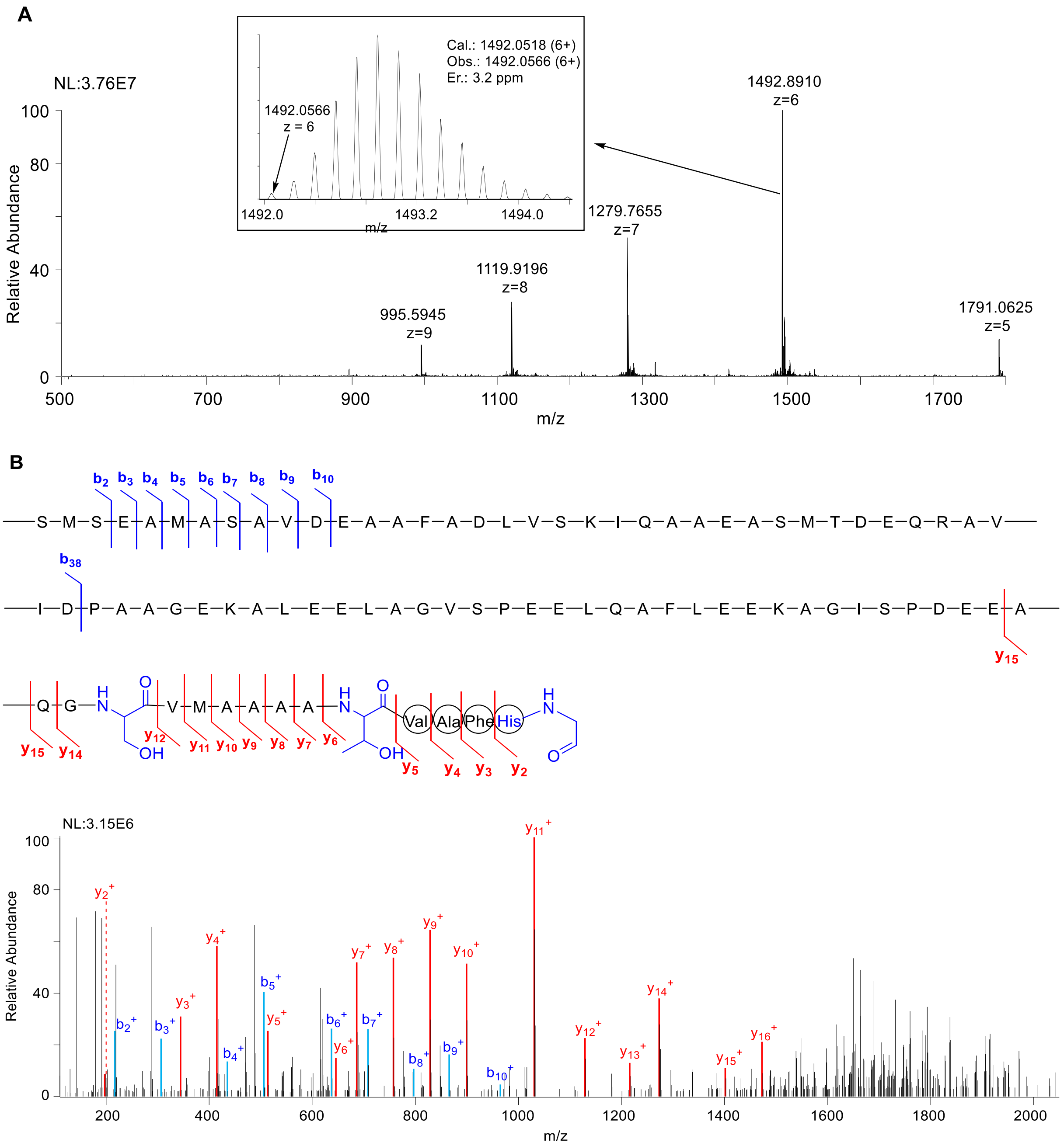

| Ions | Calc. | Obs. | Er. (ppm) | Ions | Calc. | Obs. | Er. (ppm) |
|---|---|---|---|---|---|---|---|
| **$b_2$** | 219.0803 | 219.0800 | 1.4 | **$y_2$** | 197.1028 | 197.1032 | 2.1 |
| **$b_3$** | 306.1124 | 306.1122 | 0.7 | **$y_3$** | 344.1712 | 344.1720 | 2.3 |
| **$b_4$** | 435.1549 | 435.1545 | 0.9 | **$y_4$** | 415.2083 | 415.2089 | 1.4 |
| **$b_5$** | 506.1921 | 506.1923 | 0.4 | **$y_5$** | 514.2767 | 514.2781 | 2.7 |
| **$b_6$** | 637.2325 | 637.2325 | 0.0 | **$y_6$** | 615.3244 | 615.3259 | 2.4 |
| **$b_7$** | 708.2697 | 708.2692 | 0.7 | **$y_7$** | 686.3615 | 686.3627 | 1.7 |
| **$b_8$** | 795.3017 | 795.3037 | 2.5 | **$y_8$** | 757.3986 | 757.3995 | 1.2 |
| **$b_9$** | 866.3388 | 866.3411 | 2.7 | **$y_9$** | 828.4357 | 828.4370 | 1.6 |
| **$b_{10}$** | 965.4072 | 965.4049 | 2.4 | **$y_{10}$** | 899.4728 | 899.4742 | 1.6 |
| | | | | **$y_{11}$** | 1030.5133 | 1030.5143 | 1.0 |
| | | | | **$y_{12}$** | 1129.5817 | 1129.5829 | 1.1 |
| | | | | **$y_{13}$** | 1216.6138 | 1216.6118 | 1.6 |
| | | | | **$y_{14}$** | 1273.6352 | 1273.6388 | 2.8 |
| | | | | **$y_{15}$** | 1401.6938 | 1401.6987 | 3.5 |
| | | | | **$y_{16}$** | 1472.7309 | 1472.7336 | 1.8 |

**Figure S13.** Characterization of **4c**. (**A**) HPLC-HR-MS analysis ($C_{383}H_{612}N_{100}O_{138}S_4$ $[M+6]^{6+}$ m/z: calculated 1492.3877 observed 1492.3975 and error 6.6 ppm). (**B**) HR-MS/MS analysis. The HCD fragments and the MS/MS spectrum are shown.

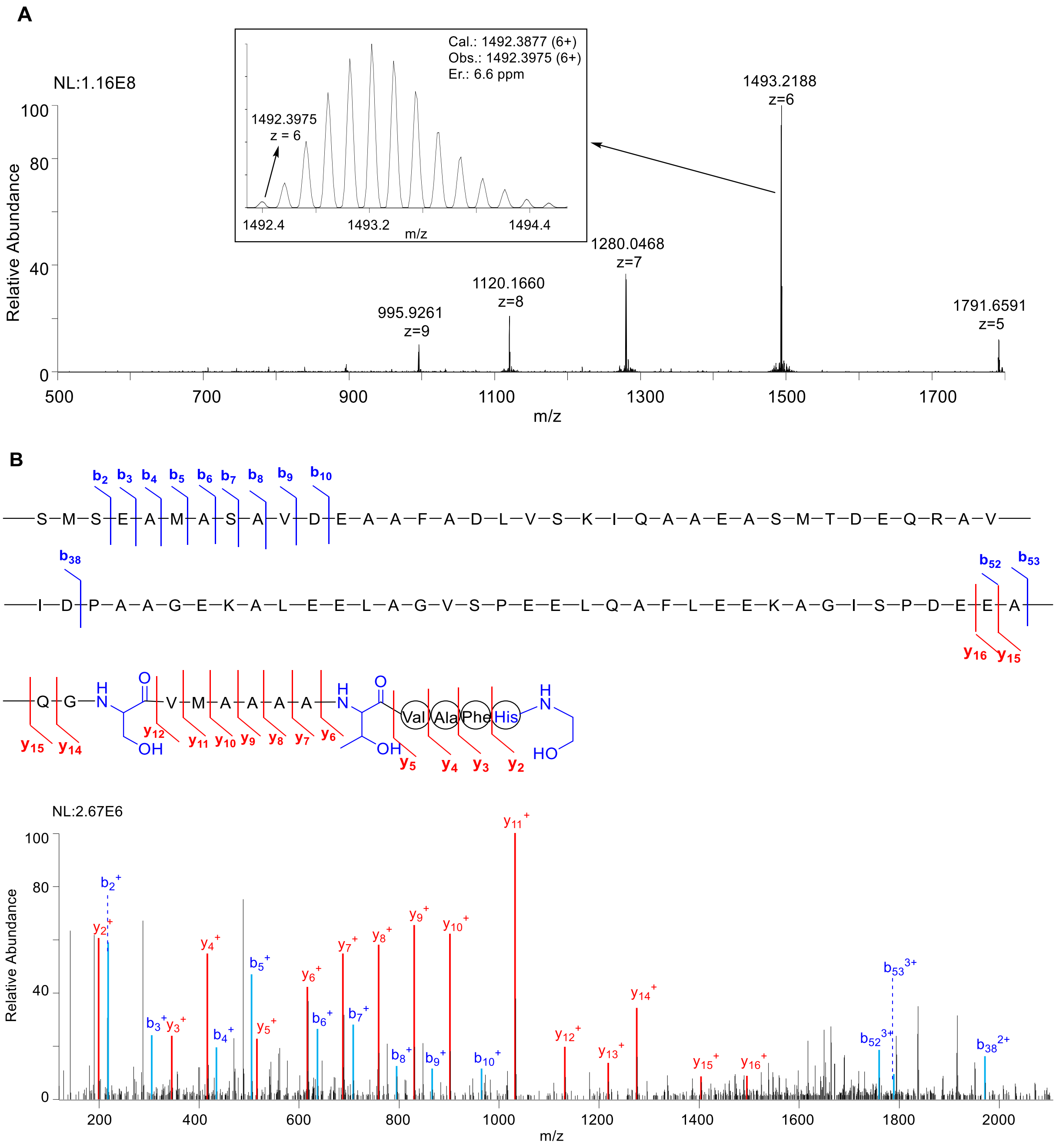

| Ions | Calc. | Obs. | Er. (ppm) | Ions | Calc. | Obs. | Er. (ppm) |
|---|---|---|---|---|---|---|---|
| $b_2$ | 219.0803 | 219.0790 | 1.4 | $y_2$ | 199.1184 | 199.1183 | 0.5 |
| $b_3$ | 306.1124 | 306.1107 | 0.7 | $y_3$ | 346.1868 | 346.1860 | 2.3 |
| $b_4$ | 435.1549 | 435.1529 | 0.9 | $y_4$ | 417.2239 | 417.2227 | 2.9 |
| $b_5$ | 506.1921 | 506.1898 | 0.4 | $y_5$ | 516.2923 | 516.2912 | 2.1 |
| $b_6$ | 637.2325 | 637.2294 | 0.0 | $y_6$ | 617.3400 | 617.3383 | 2.8 |
| $b_7$ | 708.2697 | 708.2661 | 0.7 | $y_7$ | 688.3771 | 688.3749 | 3.2 |
| $b_8$ | 795.3017 | 795.2976 | 2.5 | $y_8$ | 759.4143 | 759.4114 | 3.8 |
| $b_9$ | 866.3388 | 866.3339 | 2.7 | $y_9$ | 830.4514 | 830.4481 | 4.0 |
| $b_{10}$ | 965.4072 | 965.4005 | 2.4 | $y_{10}$ | 901.4885 | 901.4850 | 3.9 |
| $b_{52}^{3+}$ | 1759.5144 | 1759.5203 | 3.4 | $y_{11}$ | 1032.5290 | 1032.5250 | 3.9 |
| $b_{53}^{3+}$ | 1788.5251 | 1788.5249 | 0.1 | $y_{12}$ | 1131.5974 | 1131.5922 | 4.6 |
| $b_{38}^{2+}$ | 1314.2800 | 1970.9202 | 1.9 | $y_{13}$ | 1218.6294 | 1218.6245 | 4.0 |
| | | | | $y_{14}$ | 1275.6509 | 1275.6469 | 3.1 |
| | | | | $y_{15}$ | 1403.7094 | 1403.7053 | 2.9 |
| | | | | $y_{16}$ | 1474.7466 | 1474.7396 | 4.7 |

**Figure S14.** Characterization of **5**. (**A**) HPLC-HR-MS analysis ($C_{383}H_{608}N_{100}O_{136}S_5$ $[M+6H]^{6+}$ m/z: calculated 1491.7129, observed 1491.7194 and error 4.4 ppm). (**B**) HR-MS/MS analysis. The HCD fragments and the MS/MS spectrum are shown.

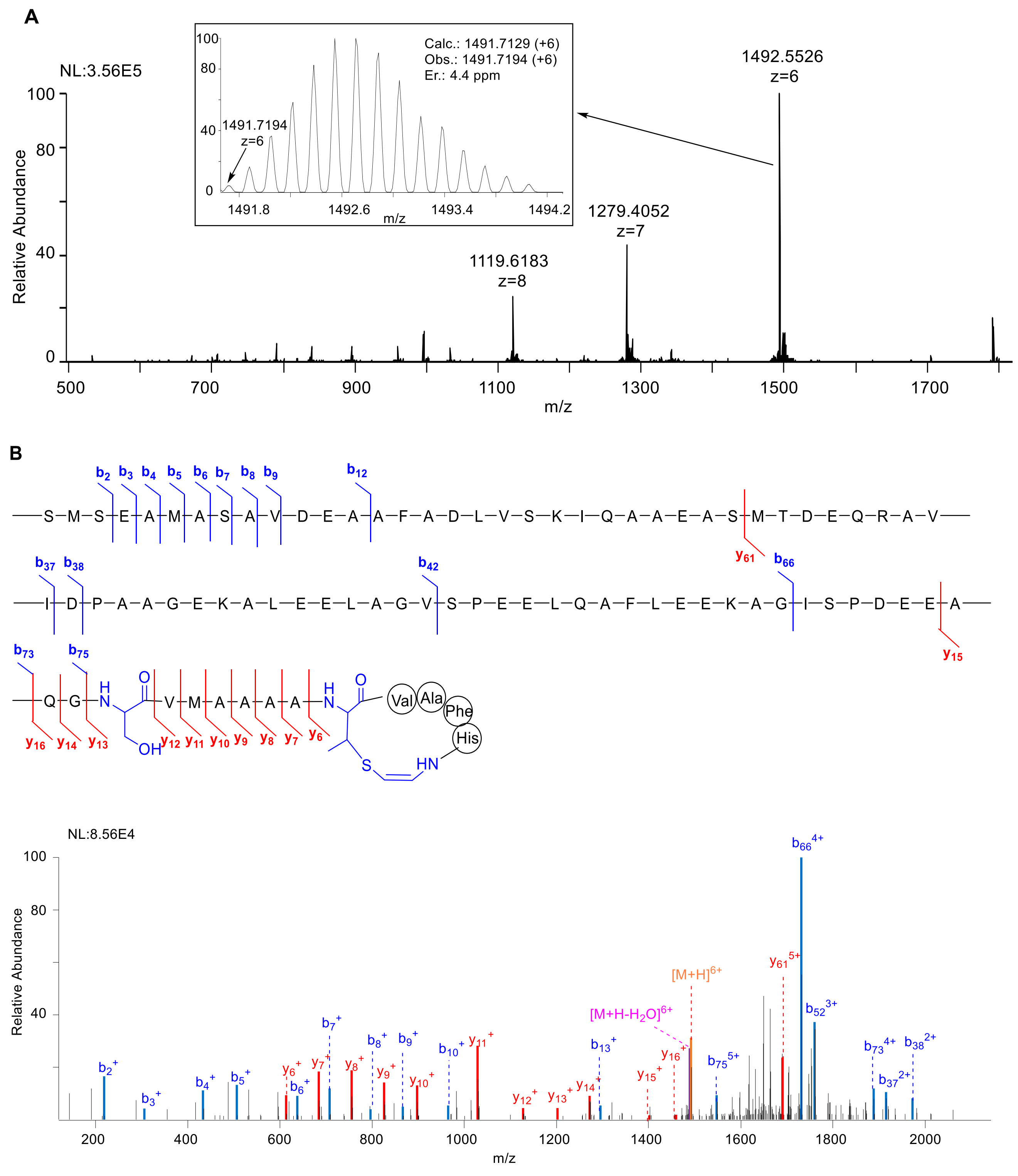

| Ions | Calc. | Obs. | Er. (ppm) | Ions | Calc. | Obs. | Er. (ppm) |
|---|---|---|---|---|---|---|---|
| $b_2^+$ | 219.0803 | 219.0794 | 4.1 | $y_2^+$ | 213.081 | - | - |
| $b_3^+$ | 306.1124 | 306.1111 | 4.2 | $y_3^+$ | 360.1494 | - | - |
| $b_4^+$ | 435.1549 | 435.1537 | 2.8 | $y_4^+$ | 431.1865 | - | - |
| $b_5^+$ | 506.1921 | 506.1903 | 3.6 | $y_5^+$ | 530.2549 | - | - |
| $b_6^+$ | 637.2325 | 637.2304 | 3.3 | $y_6^+$ | 613.292 | 613.2892 | 4.6 |
| $b_7^+$ | 708.2697 | 708.2679 | 2.5 | $y_7^+$ | 684.3291 | 684.3272 | 2.8 |
| $b_8^+$ | 795.3017 | 795.2989 | 3.5 | $y_8^+$ | 755.3662 | 755.363 | 4.2 |
| $b_9^+$ | 866.3388 | 866.335 | 4.4 | $y_9^+$ | 826.4034 | 826.4003 | 3.8 |
| $b_{12}^+$ | 965.4072 | 965.4033 | 4.0 | $y_{10}^+$ | 897.4405 | 897.4391 | 1.6 |
| $b_{13}^+$ | 1280.5138 | 1280.5135 | 0.2 | $y_{11}^+$ | 1028.481 | 1028.4781 | 2.8 |
| $b_{37}^{2+}$ | 1913.4028 | 1913.4078 | 2.6 | $y_{12}^+$ | 1127.5494 | 1127.5447 | 4.2 |
| $b_{38}^{2+}$ | 1970.9164 | 1970.9158 | 0.3 | $y_{13}^+$ | 1214.5814 | 1214.588 | 5.4 |
| $b_{52}^{3+}$ | 1759.5144 | 1759.5173 | 1.6 | $y_{14}^+$ | 1271.6029 | 1271.5985 | 3.4 |
| | | | | $y_{15}^+$ | 1399.6614 | 1399.6605 | 0.6 |
| | | | | $y_{16}^+$ | 1470.6985 | 1470.7014 | 2.0 |
| | | | | $y_{61}^{5+}$ | 1688.8177 | 1688.8082 | 5.6 |

**Figure S15.** Characterization of **TvaA$_{S\text{-}87}$-IAA**. (**A**) HPLC-HR-MS analysis ($C_{386}H_{615}N_{101}O_{140}S_5$ $[M+6H]^{6+}$ m/z: calculated 1511.8858, observed 1511.8867 and error 0.6 ppm). (**B**) HR-MS/MS analysis. The HCD fragments and the MS/MS spectrum are shown.

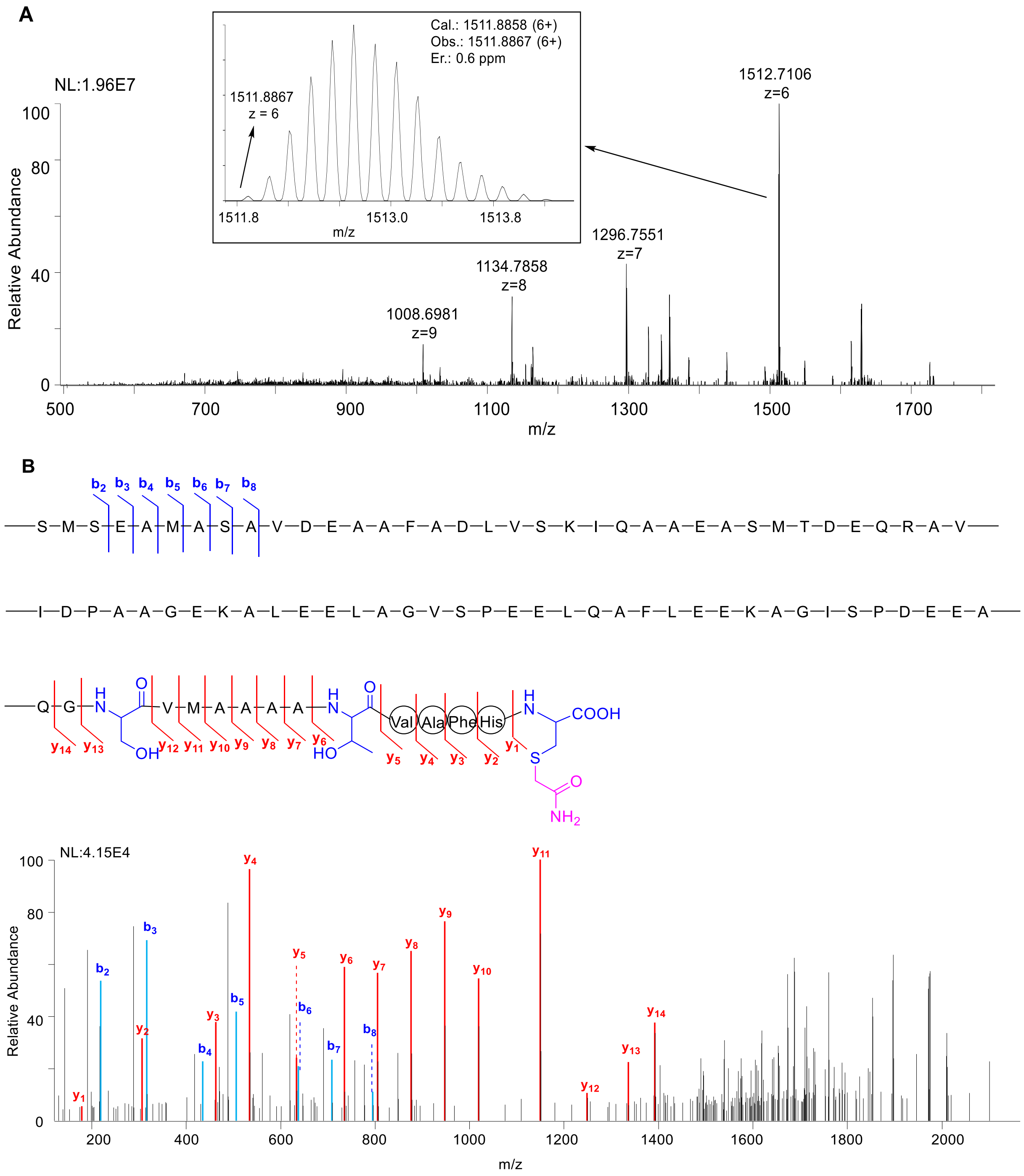

| Ions | Calc. | Obs. | Er.(ppm) | Ions | Calc. | Obs. | Er.(ppm) |
|---|---|---|---|---|---|---|---|
| $b_2^+$ | 219.0803 | 219.0798 | 2.3 | $y_1^+$ | 179.0480 | 179.0485 | 2.8 |
| $b_3^+$ | 306.1124 | 306.1122 | 0.7 | $y_2^+$ | 316.1069 | 316.1070 | 0.3 |
| $b_4^+$ | 435.1549 | 435.1547 | 0.5 | $y_3^+$ | 463.1753 | 463.1759 | 1.3 |
| $b_5^+$ | 506.1921 | 506.1922 | 0.2 | $y_4^+$ | 534.2124 | 534.2136 | 2.2 |
| $b_6^+$ | 637.2325 | 637.2338 | 2.0 | $y_5^+$ | 633.2808 | 633.2814 | 0.9 |
| $b_7^+$ | 708.2697 | 708.2715 | 2.5 | $y_6^+$ | 734.3285 | 734.3272 | 1.8 |
| $b_8^+$ | 795.3017 | 795.3029 | 1.5 | $y_7^+$ | 805.3656 | 805.3652 | 1.8 |
| | | | | $y_8^+$ | 876.4027 | 876.4012 | 1.7 |
| | | | | $y_9^+$ | 947.4398 | 947.4398 | 0.0 |
| | | | | $y_{10}^+$ | 1018.4769 | 1018.4757 | 1.2 |
| | | | | $y_{11}^+$ | 1149.5175 | 1149.5164 | 0.9 |
| | | | | $y_{12}^+$ | 1248.5858 | 1248.5969 | 8.8 |
| | | | | $y_{13}^+$ | 1335.6178 | 1335.6244 | 4.9 |
| | | | | $y_{14}^+$ | 1392.6393 | 1392.6409 | 1.1 |
| | | | | $y_{15}^+$ | 1479.0480 | 1479.0485 | 2.8 |

**Figure S16.** Characterization of **4a-IAA**. (**A**) HPLC-HR-MS analysis ($C_{385}H_{615}N_{101}O_{138}S_5$ $[M+6H]^{6+}$ m/z: calculated 1504.5542, observed 1504.5501 and error 2.7 ppm). (**B**) HR-MS/MS analysis. The HCD fragments and the MS/MS spectrum are shown.

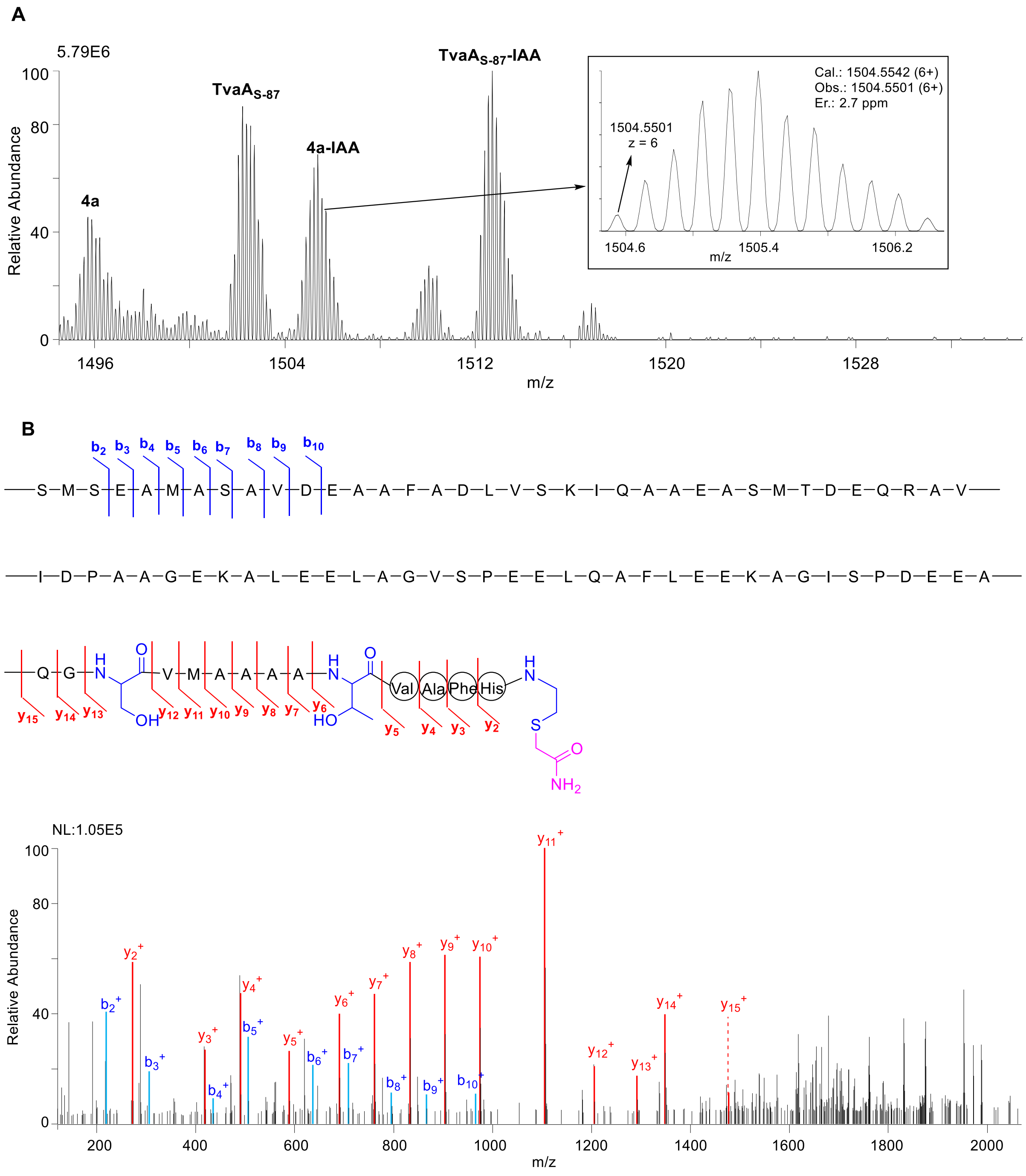

| Ions | Calc. | Obs. | Er. (ppm) | Ions | Calc. | Obs. | Er. (ppm) |
|---|---|---|---|---|---|---|---|
| $b_2$ | 219.0803 | 219.0800 | 1.4 | $y_2$ | 272.1170 | 272.1177 | 2.6 |
| $b_3$ | 306.1124 | 306.1121 | 1.0 | $y_3$ | 419.1854 | 419.1860 | 1.4 |
| $b_4$ | 435.1549 | 435.1547 | 0.5 | $y_4$ | 490.2226 | 490.2236 | 2.0 |
| $b_5$ | 506.1921 | 506.1920 | 0.2 | $y_5$ | 589.2910 | 589.2921 | 1.9 |
| $b_6$ | 637.2325 | 637.2316 | 1.4 | $y_6$ | 690.3386 | 690.3396 | 1.4 |
| $b_7$ | 708.2697 | 708.2712 | 2.1 | $y_7$ | 761.3758 | 761.3771 | 1.7 |
| $b_8$ | 795.3017 | 795.3023 | 0.8 | $y_8$ | 832.4129 | 832.4142 | 1.6 |
| $b_9$ | 866.3388 | 866.3394 | 0.7 | $y_9$ | 903.4500 | 903.4509 | 1.0 |
| $b_{10}$ | 965.4072 | 965.4066 | 0.6 | $y_{10}$ | 974.4871 | 974.4890 | 1.9 |
| | | | | $y_{11}$ | 1105.5276 | 1105.5286 | 0.9 |
| | | | | $y_{12}$ | 1204.5960 | 1204.5972 | 1.0 |
| | | | | $y_{13}$ | 1291.6280 | 1291.6333 | 4.1 |
| | | | | $y_{14}$ | 1348.6495 | 1348.6501 | 0.4 |
| | | | | $y_{15}$ | 1476.7081 | 1476.7041 | 2.7 |

**Figure S17.** Characterization of **6c**. (**A**) HPLC-HR-MS analysis ($C_{383}H_{610}N_{100}O_{137}S_4$ $[M+6H]^{6+}$ m/z: calculated 1489.3860, observed 1489.3868 and error 0.5 ppm). (**B**) HR-MS/MS analysis. The HCD fragments and the MS/MS spectrum are shown.

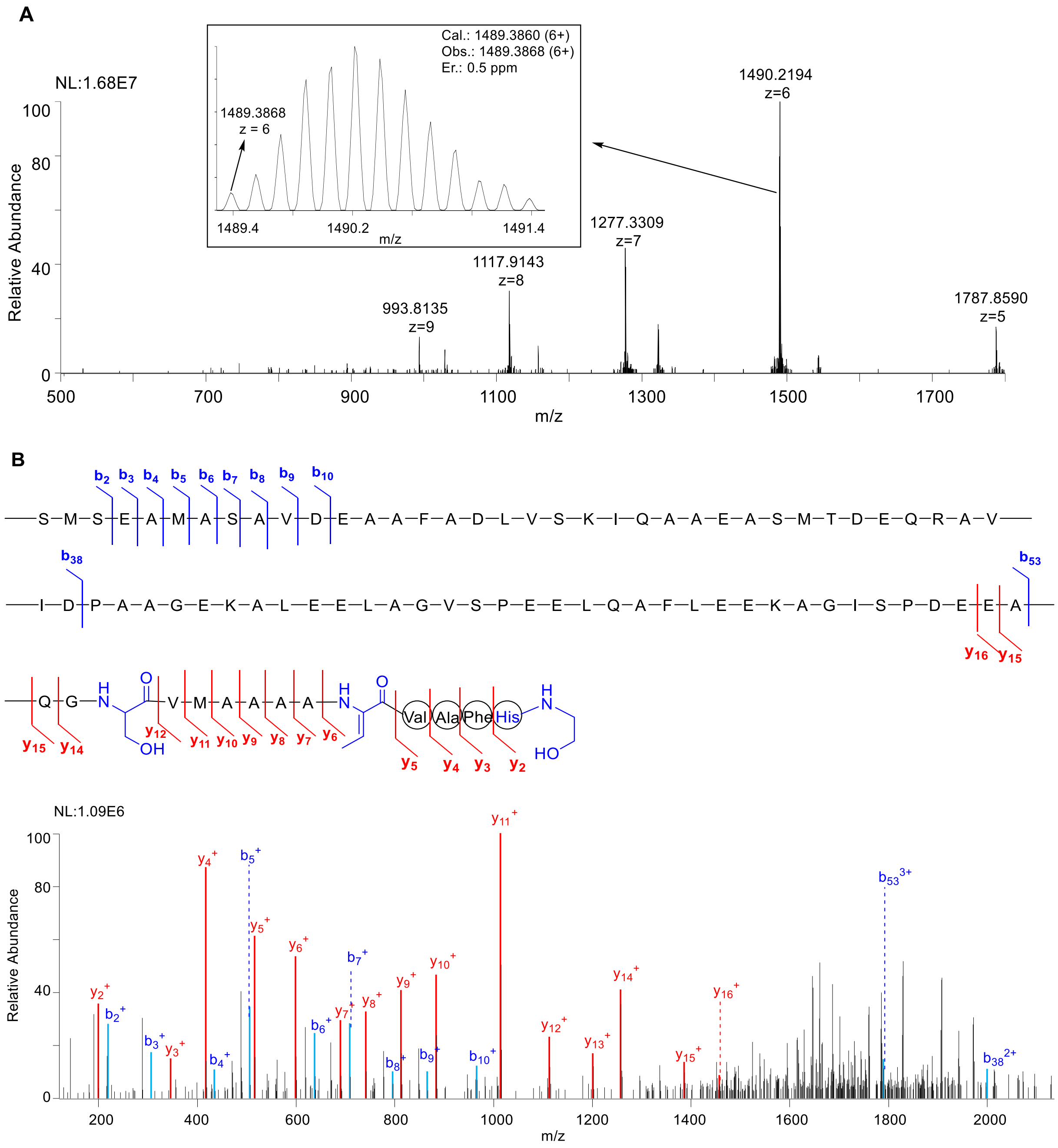

| Ions | Calc. | Obs. | Er. (ppm) | Ions | Calc. | Obs. | Er. (ppm) |
|---|---|---|---|---|---|---|---|
| $b_2$ | 219.0803 | 219.0797 | 2.7 | $y_2$ | 199.1184 | 199.1189 | 2.5 |
| $b_3$ | 306.1124 | 306.1113 | 3.6 | $y_3$ | 346.1868 | 346.1866 | 0.6 |
| $b_4$ | 435.1549 | 435.1545 | 0.9 | $y_4$ | 417.2239 | 417.2240 | 0.2 |
| $b_5$ | 506.1921 | 506.1911 | 2.0 | $y_5$ | 516.2923 | 516.2924 | 0.2 |
| $b_6$ | 637.2325 | 637.2314 | 1.7 | $y_6$ | 599.3295 | 599.3295 | 0.0 |
| $b_7$ | 708.2697 | 708.2685 | 1.7 | $y_7$ | 670.3666 | 670.3664 | 0.3 |
| $b_8$ | 795.3017 | 795.3011 | 0.8 | $y_8$ | 741.4037 | 741.4035 | 0.3 |
| $b_9$ | 866.3388 | 866.3378 | 1.2 | $y_9$ | 812.4408 | 812.4412 | 0.5 |
| $b_{10}$ | 965.4072 | 965.4077 | 0.5 | $y_{10}$ | 883.4779 | 883.4777 | 0.2 |
| $b_{53}^{3+}$ | 1788.5251 | 1788.5249 | 3.5 | $y_{11}$ | 1014.5184 | 1014.5179 | 0.5 |
| $b_{38}^{2+}$ | 1314.2800 | 1970.9202 | 0.9 | $y_{12}$ | 1113.5868 | 1113.5847 | 1.9 |
| | | | | $y_{13}$ | 1200.6188 | 1200.6188 | 0.0 |
| | | | | $y_{14}$ | 1257.6403 | 1257.6401 | 0.2 |
| | | | | $y_{15}$ | 1385.6989 | 1385.7004 | 1.1 |
| | | | | $y_{16}$ | 1456.7360 | 1456.7316 | 3.0 |

**Figure S18.** Characterization of **7**. (**A**) HPLC-HR-MS analysis ($C_{383}H_{606}N_{100}O_{135}S_5$ $[M+6H]^{6+}$ m/z: calculated 1488.7111, observed 1488.7216 and error 7.1 ppm). (**B**) HR-MS/MS analysis. The HCD fragments and the MS/MS spectrum are shown.

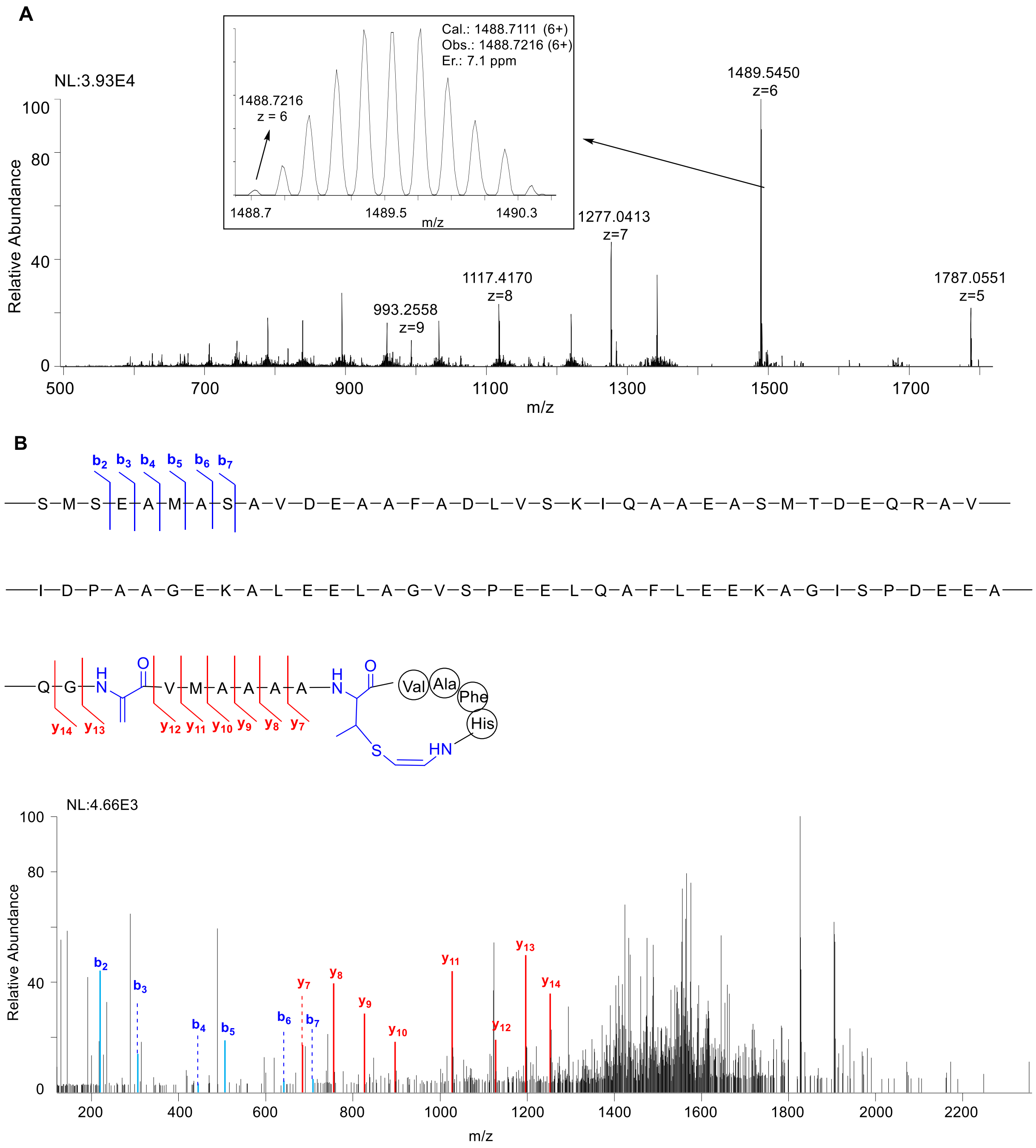

| Ions | Calc. | Obs. | Er.(ppm) | Ions | Calc. | Obs. | Er.(ppm) |
|---|---|---|---|---|---|---|---|
| $b_2^+$ | 219.0803 | 219.0799 | 1.8 | $y_2^+$ | 213.0810 | - | - |
| $b_3^+$ | 306.1124 | 306.1115 | 2.9 | $y_3^+$ | 360.1494 | - | - |
| $b_4^+$ | 435.1549 | 435.1544 | 1.1 | $y_4^+$ | 431.1865 | - | - |
| $b_5^+$ | 506.1921 | 506.1920 | 0.2 | $y_5^+$ | 530.2549 | - | - |
| $b_6^+$ | 637.2325 | 637.2308 | 2.7 | $y_7^+$ | 684.3291 | 684.3298 | 1.0 |
| $b_7^+$ | 708.2697 | 708.2693 | 0.6 | $y_8^+$ | 755.3662 | 755.3648 | 1.9 |
| | | | | $y_9^+$ | 826.4034 | 826.4042 | 1.0 |
| | | | | $y_{10}^+$ | 897.4405 | 897.4398 | 0.8 |
| | | | | $y_{11}^+$ | 1028.4810 | 1028.4830 | 1.9 |
| | | | | $y_{12}^+$ | 1127.5494 | 1127.5456 | 3.4 |
| | | | | $y_{13}^+$ | 1196.5698 | 1196.5746 | 4.0 |
| | | | | $y_{14}^+$ | 1253.5912 | 1253.5965 | 4.2 |

**Figure S19.** Characterization of the FMN-binding manner of the TvaF$_{S\text{-}87}$. (**A**) Purified TvaF$_{S\text{-}87}$. (**B**) UV spectrometry of TvaF$_{S\text{-}87}$ (**C**) HPLC-MS analysis after heating the TvaF$_{S\text{-}87}$ (i), with FMN as the standard (ii). For FMN, $[M+H]^+$ m/z calculated 457.1124; observed 457.1139.

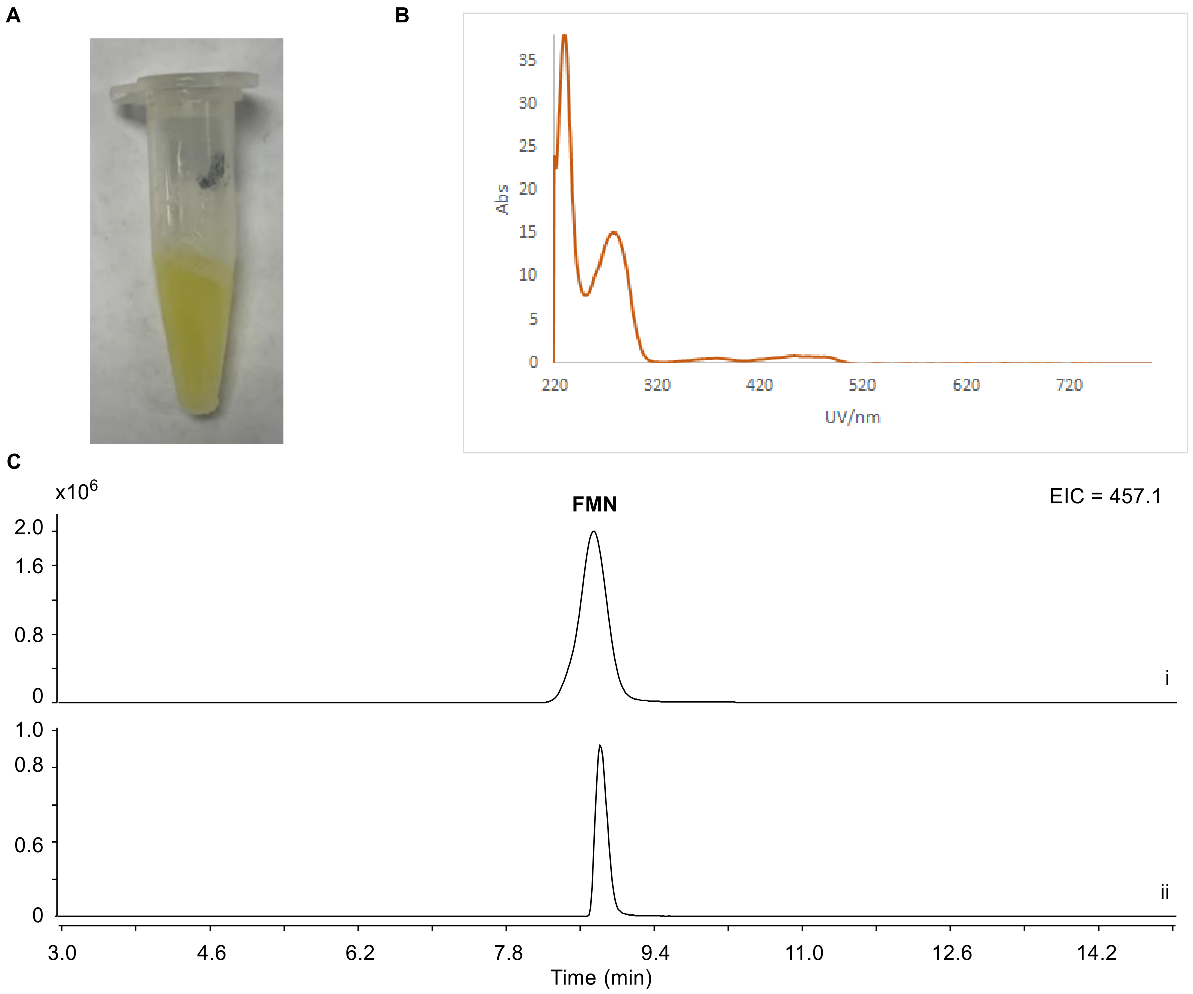

**Figure S20.** MST assay of the binding affinity of TvaE$_{S-87}$ with the LP sequence of TvaA$_{S-87}$ (i.e., TvaA$_{LP(S-87)}$, **A**), TvaF$_{S-87}$ (**B**) or the CP sequence of TvaA$_{S-87}$ (i.e., TvaA$_{core(S-87)}$, **C**). Both TvaA$_{LP(S-87)}$ and TvaA$_{core(S-87)}$ were synthesized by Genscript Biotech (Nanjing, China). Error is reported as standard deviation of three independent measurements. The fluorescence polarization was plotted on a log-scale with the substrate peptide.

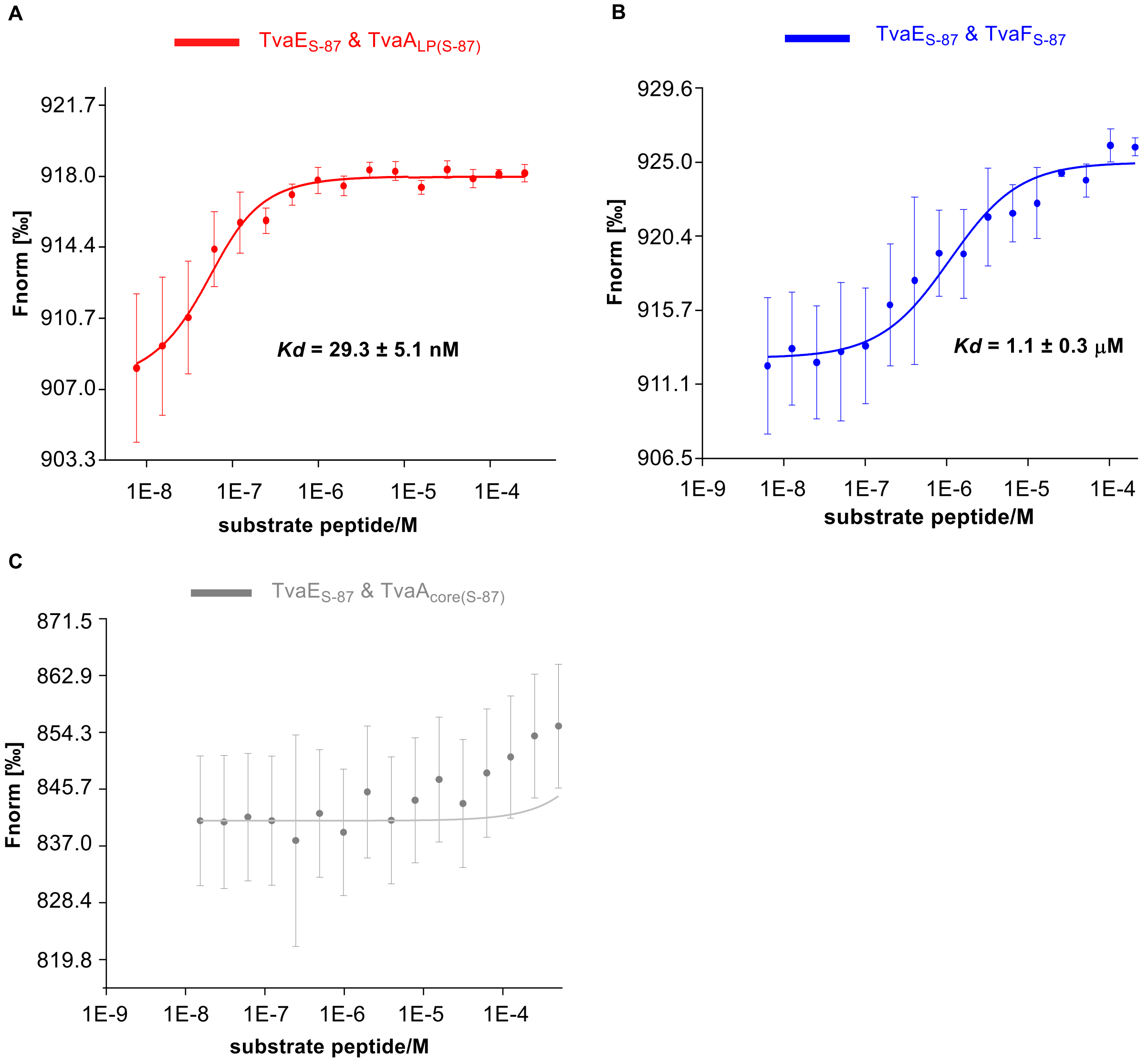

**Figure S21.** *In vivo* examination of TVA production in *S. sp.* NRRL S-87 by HPLC-MS (using **Method II**). For extracted ion chromatograms (EICs), the ESI *m/z* $M^+$ modes for TVA-YJ-1 and TVA-YJ-2 are 1305.5 and 1307.5, respectively. i, *ΔtvaC*$_{S\text{-}87}$ mutant; ii, *ΔtvaC*$_{S\text{-}87}$ mutant in which *spaKC* was expressed *in trans*; iii, *ΔtvaD*$_{S\text{-}87}$ mutant; iv *ΔtvaD*$_{S\text{-}87}$ mutant in which *spaKC* was expressed *in trans*; v, *ΔtvaE*$_{S\text{-}87}$ mutant; vi, *ΔtvaC*$_{S\text{-}87}$ mutant in which *spaKC* was expressed *in trans*; and vii, wild-type *S. sp.* NRRL S-87.

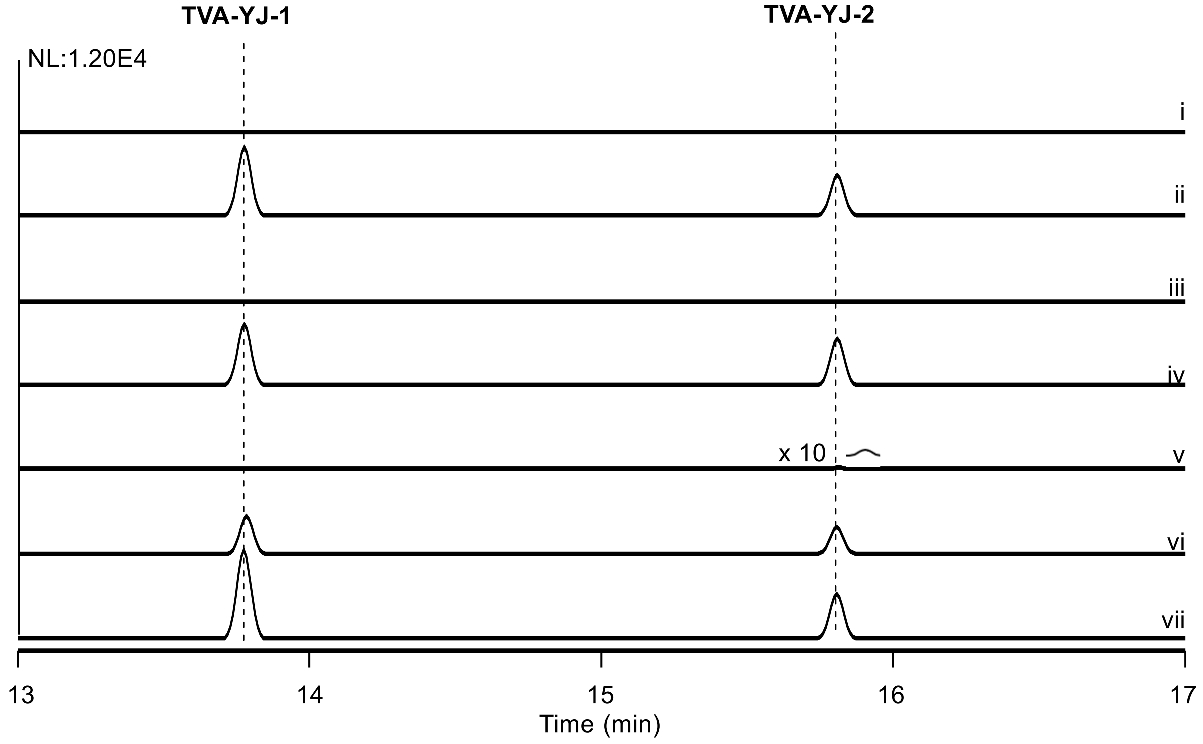

**Figure S22.** Structural homology shared by various characterized proteins with LxmX (top) or LxmY[11] (bottom) based on the prediction of their secondary structure using HHpred program (https://toolkit.tuebingen.mpg.de/tools/hhpred).[14]

**LxmX**

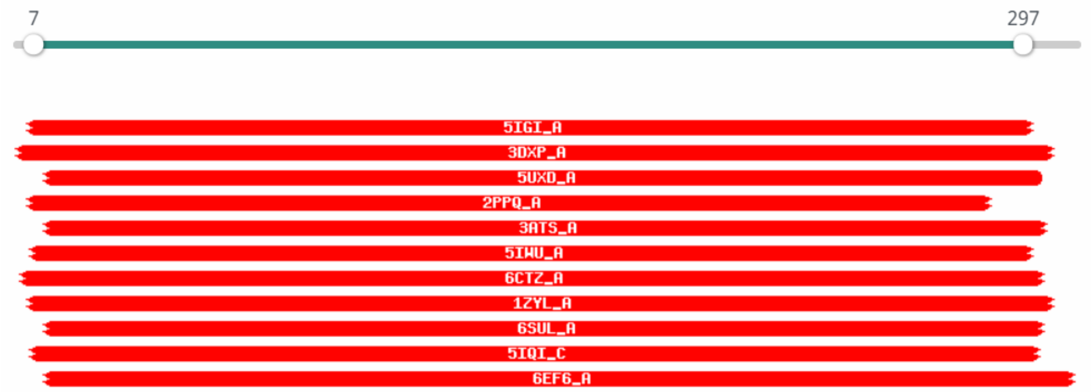


| Nr | Hit | Name | Probability | E-value | SS | Cols | Target Length |
|---|---|---|---|---|---|---|---|
| 1 | 5IGI_A | Macrolide 2'-phosphotransferase; macrolide phosphotransferase, kinase, TRANSFERASE-ANTIBIOTIC complex; HET: IPA, GMP, ZIT; 1.2A {Escherichia coli}; Related PDB entries: 5IGH_A 5IGJ_A 5IGP_A 5IGR_A 5IGS_A 5IGT_A | 99.85 | 1.5e-18 | 26 | 272 | 301 |
| 2 | 3DXP_A | putative acyl-CoA dehydrogenase; Protein kinase-like fold, structural genomics; HET: MSE; 2.32A {Ralstonia eutropha JMP134} | 99.84 | 1.3e-18 | 23.4 | 290 | 359 |

**LxmY**

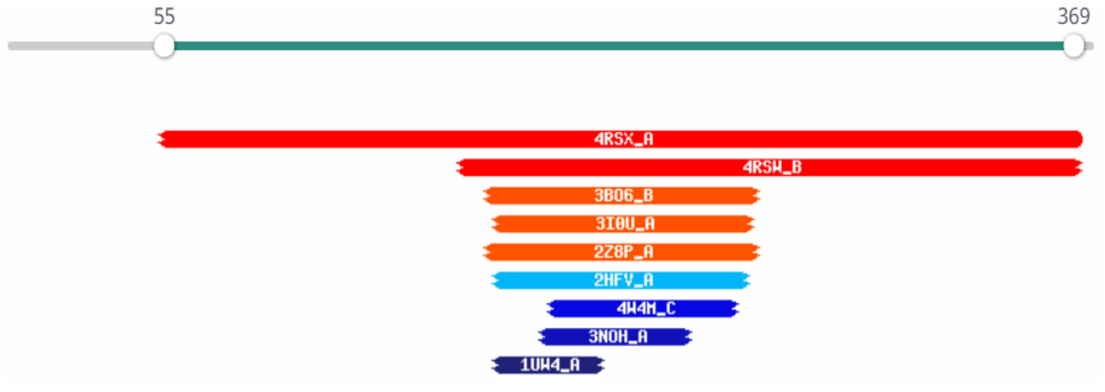


| | | | | | | | |
|---|---|---|---|---|---|---|---|
| 4 | 3I0U_A | Phosphothreonine lyase ospF (E.C.4.2.3.-); Shigella flexneri, apo-structure, type III; 2.7A {Shigella flexneri}; Related PDB entries: 3I0U_B | 93.26 | 1.1 | 10.7 | 87 | 218 |

**Figure S23.** Functional alignment of known AviCys synthetase-encoding genes. Gene functions are annotated by colored rectangular blocks.

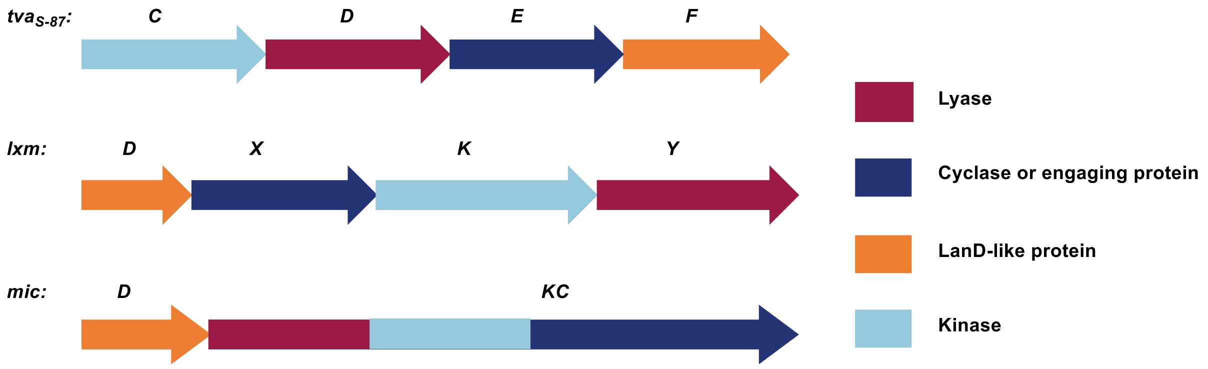

**Figure S24.** Selected gene clusters containing the homologs of *tvaCDEF*$_{S\text{-}87}$ and their associated bacterial strains.

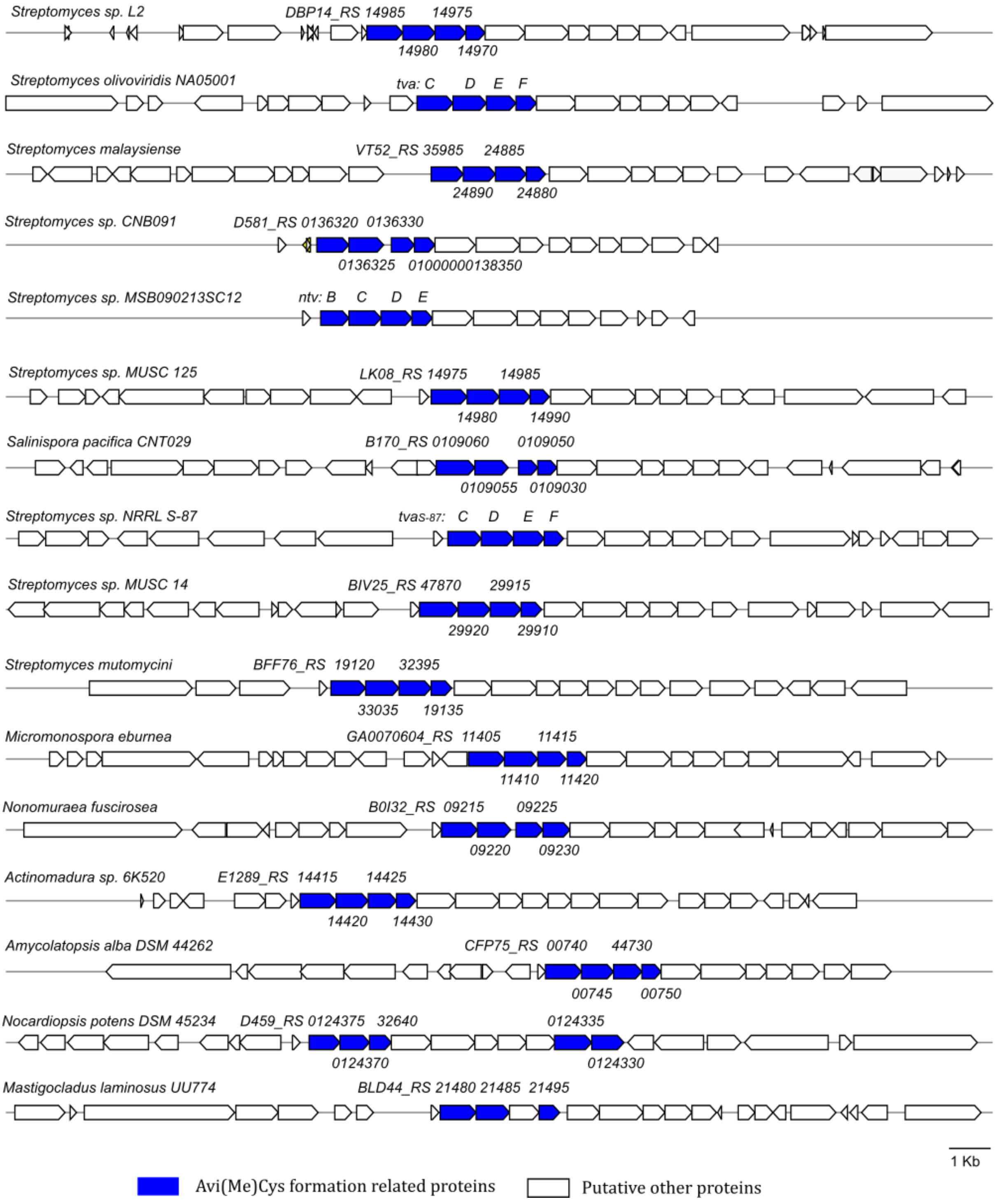

**Figure S25.** NMR spectra of TVA-YJ-1.

[1]HNMR spectrum (600 MHz, DMSO-d6)

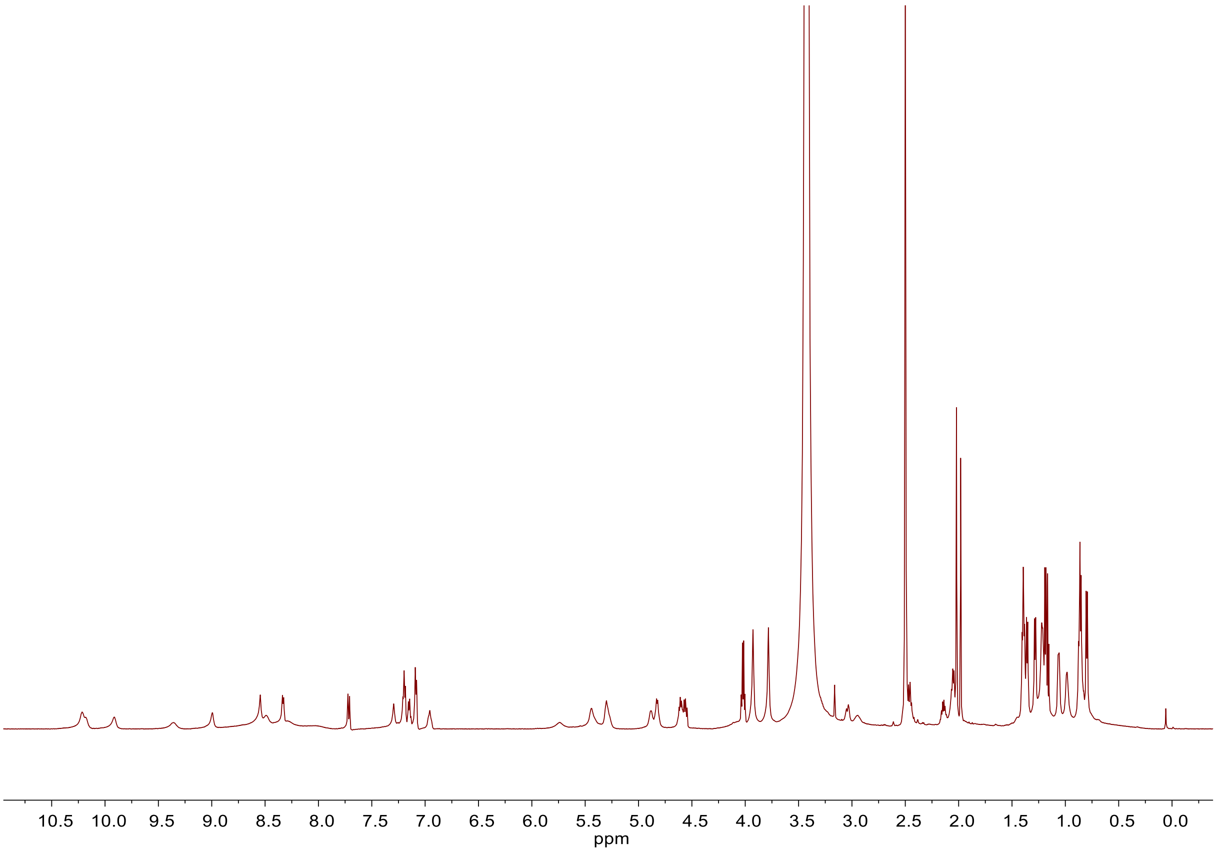

$^{13}$C NMR spectrum (600 MHz, DMSO-d6)

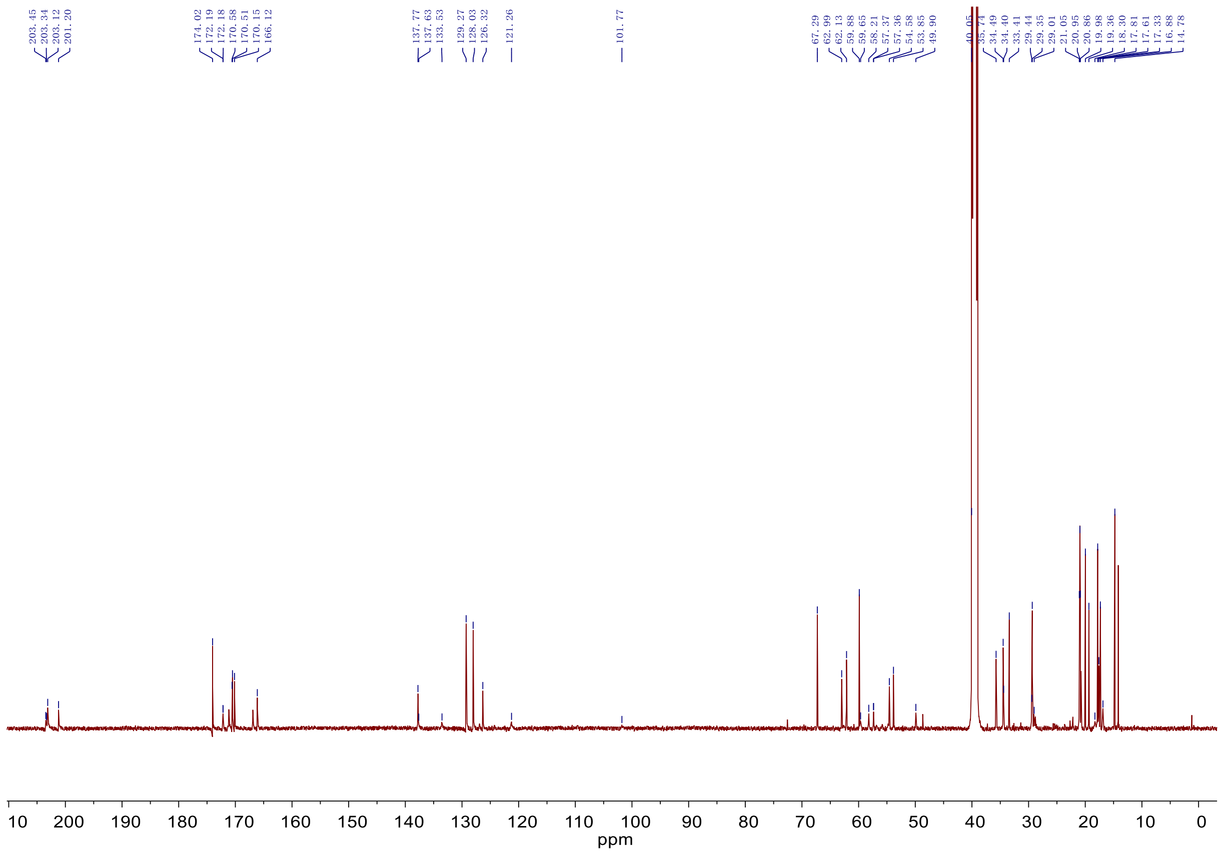

HMBC spectrum (600 MHz, DMSO-d6)

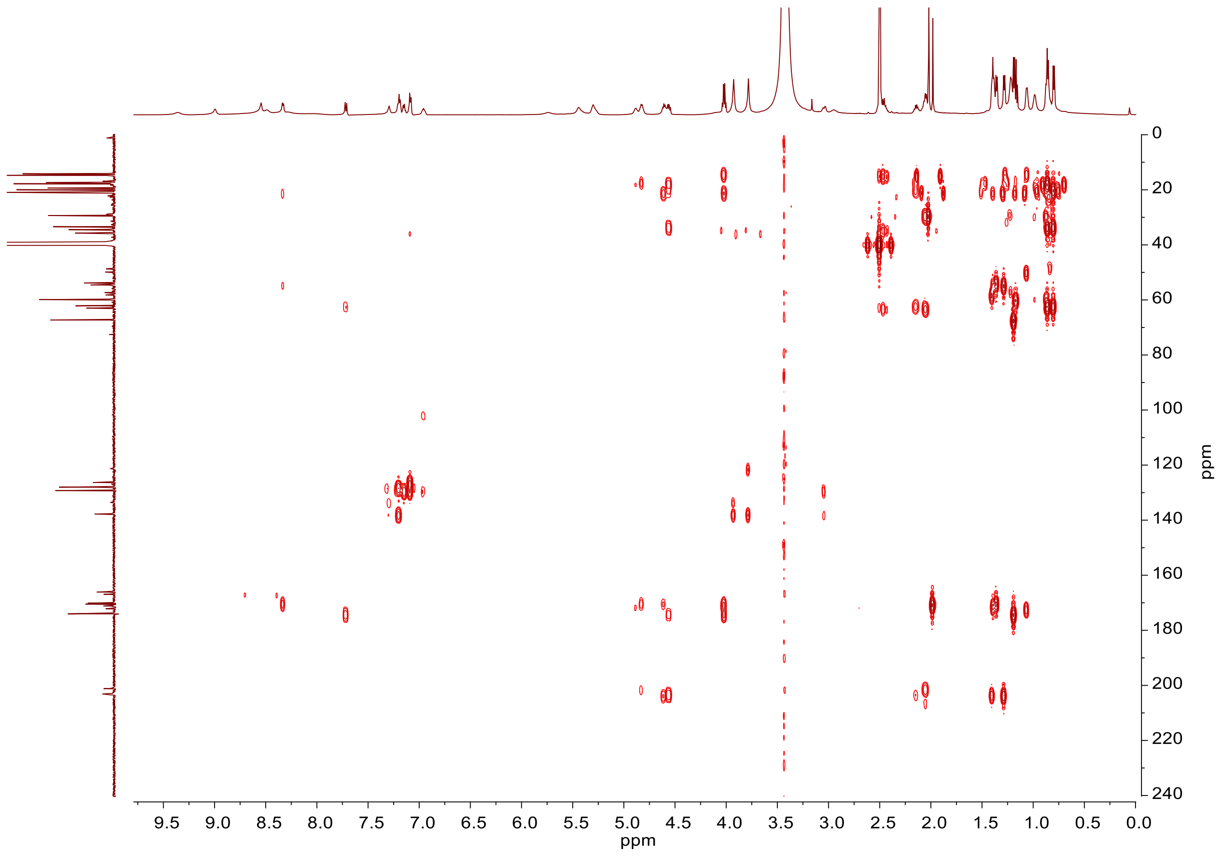

ROESY spectrum (600 MHz, DMSO-d6)

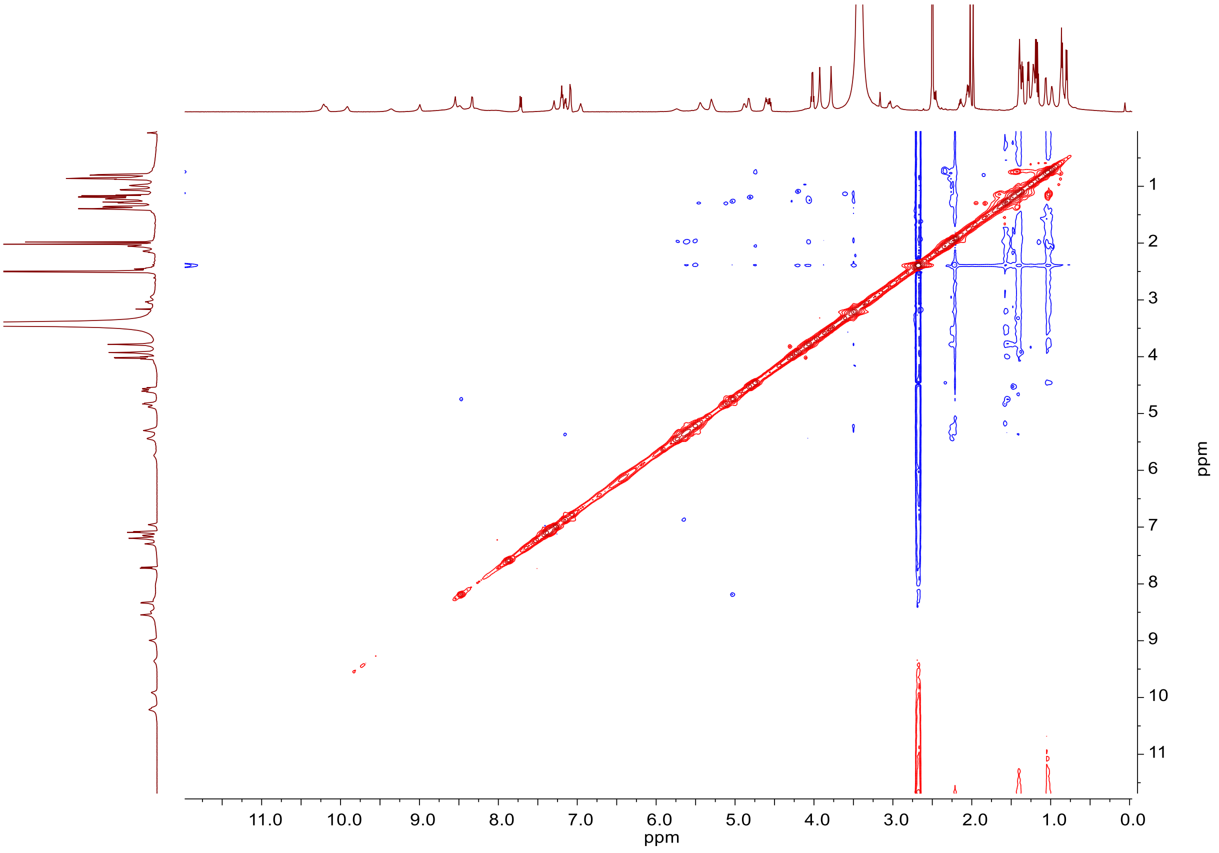

COSY spectrum (600 MHz, DMSO-d6)

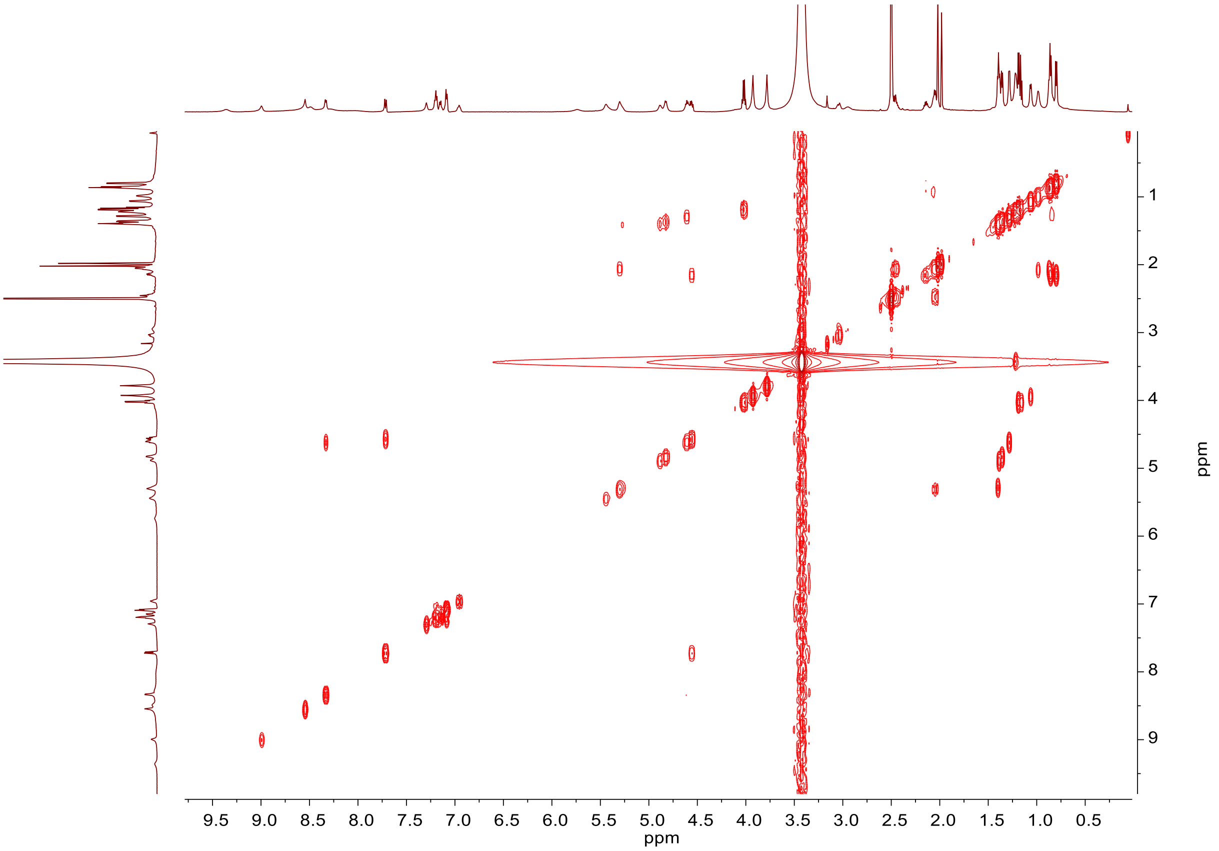

HSQC spectrum (600 MHz, DMSO-$d_6$)

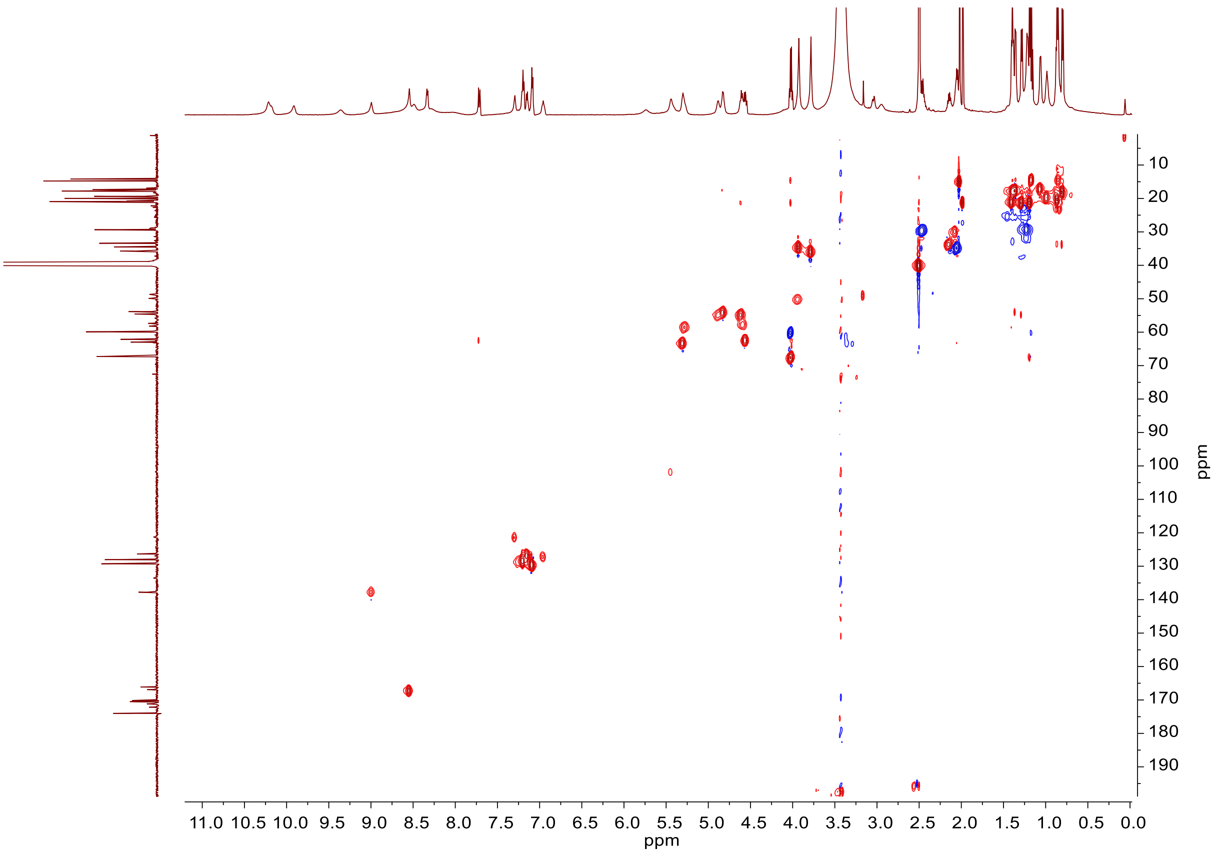

### 3. Supplementary Tables

**Table S1.** NMR data (600MHz, DMSO-d6) for TVA-YJ-1.

| | $\delta_H$,mult., ($J$ in Hz) | $\delta_C$ | HMBC | ROESY |
|---|---|---|---|---|
| **hdmHis** | | | | |
| 1-CO | | 166.1 | | |
| 2-CH$_\alpha$ | 4.57, dd (11.16,8.16) | 57.3 | | 8,36 |
| 3-CH$_\beta$ | n.a. | n.a. | | |
| 3-OH | n.a. | | | |
| 4 | | 133.5 | | |
| N$^5$-CH$_3$ | 3.94, s | 34.4 | 4,6 | 3 |
| 6 | 8.99, s | 137.6 | | |
| N$^7$-CH$_3$ | 3.79, s | 35.7 | 6,8 | 8 |
| 8 | 7.30, s | 121.3 | | 2,7 |
| 9-NH | 8.54, br s | | | 36 |
| **Phe** | | | | |
| 10-CO | | 172.2 | | |
| 11-CH$_\alpha$ | n.a. | n.a. | | |
| 12-CH$_{\beta,1}$ | 3.05, m | 40.2 | 13,14 | 14 |
| CH$_{\beta,2}$ | | | | |
| 13 | 3.04, m | 137.7 | | |
| 14,18 | 7.19, m | 128.0 | 13 | 3,12,22,15 |
| 15,17 | 7.09, m | 129.2 | | 14, |
| 16 | 7.15, m | 126.3 | | |
| 19-NH | n.a. | | | |
| **Ala** | | | | |
| 20-CO | | 170.5 | | |
| 21-CH$_\alpha$ | 4.88, br s | 54.6 | | |
| 22-CH$_\beta$ | 1.39, d (6.55) | 17.6 | 20,21 | 18 |

| | | | | |
|---|---|---|---|---|
| 23-NH | n.a. | | | |
| **Val** | | | | |
| 24-CO | | 170.5 | | |
| 25-$CH_{\alpha}$ | 4.02, m | 59.6 | | |
| 26-$CH_{\beta}$ | 2.08, m | 29.4 | | |
| 27-$CH_{\gamma,1}$ | 0.99, d (3.0) | 19.3 | 25,26,28 | |
| 28-$CH_{\gamma,2}$ | 0.88, d (6.8) | 18.3 | 26,27 | |
| 29-NH | n.a. | | | |
| **AviMeCys** | | | | |
| 30-CO | | 171.1. | | |
| 31-$CH_{\alpha}$ | 4.57, br s | 57.3 | | |
| 32-$CH_{\beta}$ | n.a. | n.a. | | |
| 33-$CH_{\gamma}$ | 1.23, d (6.8) | 29.0 | 31 | |
| 34 | 5.44 | 101.7 | | |
| 35 | 6.95 | 126.8 | 1,34 | |
| 36-NH | 9.35, br | | | |
| 37-NH | 8.48, m | | | 31 |
| **Ala** | | | | |
| 38-CO | | 172.2 | | |
| 39-$CH_{\alpha}$ | 3.94, m | 49.8 | | |
| 40-$CH_{\beta}$ | 1.07, d (6.5) | 16.8 | 38,39 | |
| 41-NH | n.a. | | | |
| **Ala’** | | | | |
| 42-CS | | 203.3 | | |
| 43-$CH_{\alpha}$ | 5.29, m | 58.2 | 42 | |
| 44-$CH_{\beta}$ | 1.41, d (6.6) | 20.8 | 42,43 | |
| 45-NH | 9.91, br s | | | |
| **Ala’** | | | | |
| 46-CS | | 203.5 | | |

| | | | | |
|---|---|---|---|---|
| 47-$CH_{\alpha}$ | 4.63, dd (13.5,6.7) | 54.5 | 46,48 | |
| 48-$CH_{\beta}$ | 1.28, d (6.8) | 21.0 | | |
| 49-NH | 8.33, d (6.8) | | 47,48,50 | 51 |
| **Ala** | | | | |
| 50-CO | | 170.1 | | |
| 51-$CH_{\alpha}$ | 4.83, q (7.1) | 53.8 | 50,52 | 49 |
| 52-$CH_{\beta}$ | 1.36, d (7.1) | 17.3 | | |
| 53-NH | 10.21, br s | | | |
| **Met'** | | | | |
| 54-CS | | 201.2 | | |
| 55-$CH_{\alpha}$ | 5.30, m. | 62.9 | 60 | |
| 56-$CH_{\beta}$ | 2.05, m | 34.3 | 54 | |
| 57-$CH_{\gamma}$ | 2.46, m | 29.3 | 58 | |
| 58-$CH_{\varepsilon}$ | 2.02, s | 14.7 | | |
| 59-NH | 10.20, br s | | | |
| **Val'** | | | | |
| 60-CS | | 203.2 | | |
| 61-$CH_{\alpha}$ | 4.56, dd (9.3,6.8) | 62.1 | 60,62,63,64,66 | |
| 62-$CH_{\beta}$ | 2.15, q (6.8) | 33.4 | 63,64 | |
| 63-$CH_{\gamma,1}$ | 0.87, d (6.9) | 19.9 | | |
| 64-$CH_{\gamma,2}$ | 0.81, d (7.7) | 17.8 | | |
| 65-NH | 7.72, d (9.4) | | 66 | |
| **LA** | | | | |
| 66-CO | | 174.0 | | |
| 67-$CH_{\alpha}$ | 4.02, q(6.6) | 67.2 | 66,69 | |
| 68-OH | n.a. | | | |
| 69-$CH_3$ | 1.19, d(5.7) | 20.9 | | |

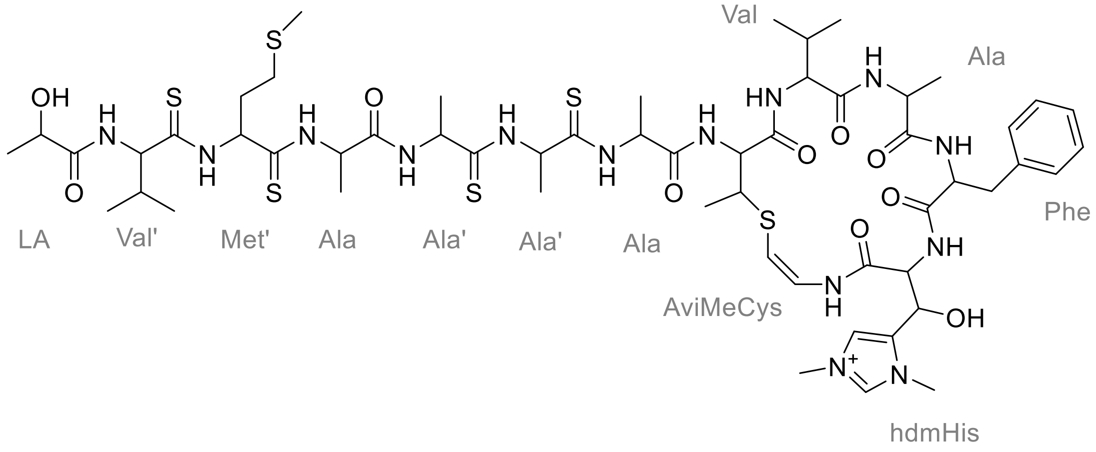
OH
S
O
S
O
S
S
O
HN
O
O
NH
O
NH
O
OH
LA
Val'
Met'
Ala
Ala'
Ala'
Ala
Val
Ala
Phe
AviMeCys
hdmHis

**Table S2.** Lan synthase homologs in the genome of *Streptomyces sp.* NRRL S-87 and *Streptomyces laurentii*.

| **Strain** | **LanKC/LanL** | **Accession number** |
|---|---|---|
| *Streptomyces sp.* NRRL S-87 | LanKC1 | WP_037887254.1 |
| | LanKC2 | WP_051795233.1 |
| *Streptomyces laurentii* | LanL | BAU81016.1 |

**Table S3.** Related bacterial strains and plasmids used in this study.

| Strains/Plasmids | Characteristic(s) | Sources/References |
|---|---|---|
| ***Streptomyces* sp.** | | |
| **NRRL S-87** | Wild type strain, TVA-YJ-1/2-producing strain | NRRL |
| YJ101 | The *tvaA*$_{S\text{-}87}$ in-frame deletion mutant of *S*. sp. NRRL S-87 | This study |
| YJ102 | The *tvaC*$_{S\text{-}87}$ in-frame deletion mutant of *S*. sp. NRRL S-87 | This study |
| YJ103 | The *tvaD*$_{S\text{-}87}$ in-frame deletion mutant of *S*. sp. NRRL S-87 | This study |
| YJ104 | The *tvaE*$_{S\text{-}87}$ in-frame deletion mutant of *S*. sp. NRRL S-87 | This study |
| YJ105 | The *tvaF*$_{S\text{-}87}$ in-frame deletion mutant of *S*. sp. NRRL S-87 | This study |
| YJ106 | The YJ102 derivative carrying *tvaC*$_{S\text{-}87}$ *in trans* | This study |
| YJ107 | The YJ102 derivative carrying *tvaC*$_{S\text{-}87}$ -N241A&D258A *in trans* | This study |
| YJ108 | The YJ104 derivative carrying *PermE** in upstream of *tva*$_{S\text{-}87}$ gene cluster | This study |
| YJ109 | The YJ102 derivative carrying *spaKC in trans* | This study |
| YJ110 | The YJ103 derivative carrying *spaKC in trans* | This study |
| YJ111 | The YJ104 derivative carrying *spaKC in trans* | This study |
| ***Streptomyces laurentii*** | | |
| **ATCC 31255** | Wild type strain, thiostrepton-producing strain | ATCC |
| YJ112 | TVA-YJ-1/2-producing strain, *S. laurentii* derivative | This study |
| YJ113 | The *lanL*$_{SL}$ in-frame deletion mutant of *S. laurentii* | This study |
| YJ114 | TVA-YJ-1/2-producing strain, YJ113 derivative | This study |
| ***Escherichia coli*** | | |
| DH5α | Host for general cloning | Transgen |
| BL21 (DE3) | Host for protein expression | Transgen |

| | | |
|---|---|---|
| ET12567/pUZ8002 | Donor strain for conjugation between *E. coli* and *Streptomyces* | ref. 15 |
| **Plasmids** | | |
| pMD19-T | *E. coli* subcloning vector | Takara |
| pOJ260 | *E.coli-Streptomyces* shuttle vector for gene inactivation, replicating in *E. coli* but not in *Streptomyces* | ref. 7 |
| pWHU2653 | *E.coli-Streptomyces* shuttle vector for gene inactivation with CRISPR/Cas9 editing system | ref. 8 |
| pJTU2554 | *E.coli-Streptomyces* shuttle vector for heterogeneous expression | Shanghai Jiaotong University |
| pSET152 | *E.coli-Streptomyces* shuttle vector for gene complementation | ref. 7 |
| pRSFDeut-1 | Protein expression vector used in *E.coli*, encoding N-terminal His-tag, kanamycin resistance | Novagen |
| pET28a(+) | Protein expression vector used in *E.coli*, encoding N-terminal His-tag, kanamycin resistance | Novagen |
| pGEX-4T-1 | Protein expression vector used in *E.coli*, encoding N-terminal GST-tag, ampicillin resistance | Novagen |
| pYJ1001 | pMD19-T derivative containing partial $tvaA_{S\text{-}87}$ fragment | This study |
| pYJ1002 | pMD19-T derivative containing partial $tvaA_{S\text{-}87}$ fragment | This study |
| pYJ1003 | pOJ260 derivative for $tvaA_{S\text{-}87}$ in-frame deletion | This study |
| pYJ1004 | pWHU2653 derivative containing the *sg* sequence of $tvaC_{S\text{-}87}$ | This study |
| pYJ1005 | pWHU2653 derivative containing the *sg* sequence of $tvaD_{S\text{-}87}$ | This study |
| pYJ1006 | pWHU2653 derivative containing the *sg* sequence of $tvaE_{S\text{-}87}$ | This study |
| pYJ1007 | pWHU2653 derivative containing the *sg* sequence of $tvaF_{S\text{-}}$ | This study |

| | | |
|---|---|---|
| | *87* | |
| pYJ1008 | pWHU2653 derivative for *tvaC*$_{S\text{-}87}$ in-frame deletion | This study |
| pYJ1009 | pWHU2653 derivative for *tvaD*$_{S\text{-}87}$ in-frame deletion | This study |
| pYJ1010 | pWHU2653 derivative for *tvaE*$_{S\text{-}87}$ in-frame deletion | This study |
| pYJ1011 | pWHU2653 derivative for *tvaF*$_{S\text{-}87}$ in-frame deletion | This study |
| pYJ1012 | pSET152 derivative containing *tvaC*$_{S\text{-}87}$ under the control of *PermE** | This study |
| pYJ1013 | pSET152 derivative containing *tvaC*$_{S\text{-}87}$- *N241A&D258A* under the control of *PermE** | This study |
| pYJ1014 | pOJ260 derivative containing *PermE** | This study |
| pYJ1015 | pSET152 derivative containing *spaKC* under the control of *PermE** | This study |
| pYJ1017 | pOJ260 derivative for *lanL*$_{SL}$ in-frame deletion | This study |
| pYJ1016 | pJTU2554 derivative containing biosynthetic gene cluster of TVA-YJ-1/2 under the control of *PermE** | This study |

**Table S4.** Primers used in this study.

| **Primers** | **Sequence (from 5' to 3')** |
|---|---|
| *tvaA*$_{S\text{-}87}$-R-for | AAACTCGAG TTCCACTGTTGATCTGAGTCG |
| *tvaA*$_{S\text{-}87}$-R-rev | ACGTCTAGA GCGTCGATGTGGTCGCGGTG |
| *tvaA*$_{S\text{-}87}$-L-for | CGCAAGCTTAGCCGAGGAGGGAGGTGGCCATGG |
| *tvaA*$_{S\text{-}87}$-L-rev | AAACTCGAG GGCGGCCTGGATCTTACTGACGAGG |
| sg-for | accaccaccaccacCACTGAGCTAGCTTCAGACGTG |
| *tvaC*$_{S\text{-}87}$-sg-rev | gccgtctctccgactcaggcGCTGGATCCTACCAACCGGC |
| *tvaC*$_{S\text{-}87}$-sg-for | gcctgagtcggagagacggcGTTTTAGAGCTAGAAATAGC |
| sg-rev | gactagaggatccccgggtATCTAGAAAAAAAACCCCGCC |
| *tvaD*$_{S\text{-}87}$-sg-rev | gggggaacctgacactcaccGCTGGATCCTACCAACCGGC |
| *tvaD*$_{S\text{-}87}$-sg-for | ggtgagtgtcaggttcccccGTTTTAGAGCTAGAAATAGC |
| *tvaE*$_{S\text{-}87}$-sg-rev | catggccgatctcgcgttccGCTGGATCCTACCAACCGGC |
| *tvaE*$_{S\text{-}87}$-sg-for | ggaacgcgagatcggccatgGTTTTAGAGCTAGAAATAGC |
| *tvaF*$_{S\text{-}87}$-sg-rev | gcaagaccaccagcacgtccGCTGGATCCTACCAACCGGC |
| *tvaF*$_{S\text{-}87}$-sg-for | ggacgtgctggtggtcttgcGTTTTAGAGCTAGAAATAGC |
| *tvaC*$_{S\text{-}87}$-L-for | gacctgcaggcatgcaagcttGACCGCTGAACAGGTACGCG |
| *tvaC*$_{S\text{-}87}$-L-rev | tacatcagggtggaCAGCATGGTCTCCTCGTTACCC |
| *tvaC*$_{S\text{-}87}$-R-for | atgctgTCCACCCTGATGTACGCGG |
| *tvaC*$_{S\text{-}87}$-R-rev, | gcttgcggcagcgtgaagcttGCCGACCAGGCGCGCAGG |
| *tvaD*$_{S\text{-}87}$-L-for | gacctgcaggcatgcaagcttGCCTGCACAGCCAGCAGG |
| *tvaD*$_{S\text{-}87}$-L-rev | acagcggtgGCGGGACAGGTGAGGCAG |
| *tvaD*$_{S\text{-}87}$-R-for | acctgtcccgcCACCGCTGTCTGCTGGTCG |
| *tvaD*$_{S\text{-}87}$-R-rev, | gcttgcggcagcgtgaagctttCTCGGACCGCTTGCTGG |
| *tvaE*$_{S\text{-}87}$-L-for | gacctgcaggcatgcaagcttacccgtggaccacatcatcgc |
| *tvaE*$_{S\text{-}87}$-L-rev | acgccgaccaggcgcgCTGTGGCGCCGCTTGCAG |
| *tvaE*$_{S\text{-}87}$-R-for | acagCGCGCCTGGTCGGCGTCC |
| *tvaE*$_{S\text{-}87}$-R-rev, | gcttgcggcagcgtgaagcttCCGGTGATCTTGGGGACG |

| | |
|---|---|
| *tvaF<sub>S-87</sub>*-L-for | gacctgcaggcatgcaagCTTTGGGGCGAGCTCCGGCCC |
| *tvaF<sub>S-87</sub>*-L-rev | tcttcgcggagctgatcGACGCGCGGCCCCATGAGGGC |
| *tvaF<sub>S-87</sub>*-R-for | GTCGATCAGCTCCGCGAAGAC |
| *tvaF<sub>S-87</sub>*-R-rev, | gcttgcggcagcgtgaagCTTACAGGGCGTGGCAGATGG |
| *tvaC<sub>S-87</sub>*-C-for | gaaatcgataagcttggatccGCCACGGTGGCCTTCCAC |
| *tvaC<sub>S-87</sub>*-C-rev | gggctgcaggtcgactctagaTGAGGCAGGACGGGGAGG |
| *tvaC<sub>S-87</sub>*-N241A-for | GACGACGCAATCCTGCTCAAGGACGCCGACAGTAGG |
| *tvaC<sub>S-87</sub>*-N241A-rev | GTCGGCGTCCTTGAGCAGGATGTTGTCTGCTCGCAG |
| *tvaC<sub>S-87</sub>*-D258A-for | CTGGTCGCATGGGAGCTGGCCGGGTTCGGC |
| *tvaC<sub>S-87</sub>*-D258A-rev | CTCCCATGCGACCAGTGCAACCGTGGGATG |
| *PermE*-for | ctatgacatgattacgaattcGGTACCAGCCCGACCCGAGCACGC |
| *2kb*-rev | gacggccagtgccaagcttCCGCGGCCGCGGAGTCCCGGG |
| *spaKC*-for | gaaatcgataagcttggatccTCCTTGAGGGCTCGTCAGG |
| *spaKC*-rev | gggctgcaggtcgactctagaGGCTGTTCTCGAGCTCCTGC |
| *lanL<sub>SL</sub>*-L-for | caggtcgactctagaggatccAGATACAGCATCGGGTGCAGC |
| *lanL<sub>SL</sub>*-L-rev | ttctcgcgcacGAACTGGCCCGTCTCCTCG |
| *lanL<sub>SL</sub>*-R-for | ggccagttcGTGCGCGAGAACCGCCGG |
| *lanL<sub>SL</sub>*-R-rev | ctatgacatgattacgaattcCGCGGGCCTCCAGGAGCT |
| YJ-1-for | CCGGGGATCCTAAGGCATCCATGACATAAACGGAAC |
| YJ-1-rev | GTCATGTCCTGCTCCAGATCCACC |
| YJ-2-for | GGTGGATCTGGAGCAGGACATGAC |
| YJ-2-rev | ATCGAGCGCCCGGTTCGAGGCCGG |
| *PermE2*-for | GGTACCAGCCCGACCCGAGCACGC |
| *PermE2*-rev | GGATGCCTTAGGATCCCCGGGTACCGAGCTCTC |
| YJ-3-for | cctcgaaccgggcgctcgatTCTAGAGTCGACCTGCAGCCCAA |
| YJ-3-rev | gctcgggtcgggctggtaccACTAGTAGCAGCGGGCCGCGTTG |
| TvaD-H24A-R26A-F | GCCTTCGCACACGCAGAGTGGGGCGAGCTCCGGCCC |
| TvaD-H24A-R26A-R | CCACTCTGCGTGTGCGAAGGCCGACTCGCTCAGGAC |
| CDF-TvaE-NcoI-F | TAATAAGGAGATATACCATGGGCATGACCCAATCCGCCG |

| | |
|---|---|
| CDF-TvaE-HindIII-R | GCATTATGCGGCCGCAAGCTTTCATCGGCCGGTCACCTC |
| TvaEF-F | GGTGCGGCGGCAGTGGCGCTGGGCACCTGGGCGGAT |
| TvaEF-R | CAGCGCCACTGCCGCCGCACCGCCACCCTTGTGGTC |
| ET-TrxDC-NcoI-F | TAAGAAGGAGATATACCATGGGCATGGGCATGAGCGATAAAATTAT |
| ET-TrxDC-XhoI-R | GGTTTCTTTACCAGACTCGAGTCACGCCTGCGCCTCCCG |
| RSF-SA-NcoI-F | TAATAAGGAGATATACCATGGGCCATCATCATCATCATCACAGCAGC |
| RSF-SA-HindIII-R | GCATTATGCGGCCGCAAGCTTTCAACAGTGGAAGGCCACCG |
| DC-241N/A-F | CGAGACGACGCCATCCTGCTCAAGGACGCCGACAGT |
| DC-241N/A-R | GAGCAGGATGGCGTCGTCTCGCAGGTCGAAGTGGAC |
| LP-For | CAGATTGGTGGATCCATGTCTGAAGCAATGGCT |
| Lp-Rev | GAATTCGAATTCTCAGCCCTGCGCCTCCTCGTC |

(11) Tao, M.; Xu, M.; Zhang, F.; Cheng, Z.; Bashiri, G.; Wang, J.; Hong, J.; Wang, Y.; Xu, L.; Chen, X.; Huang, S. X.; Lin, S.; Deng, Z., Functional genome mining reveals a novel class V lanthipeptide

containing a D-amino acid introduced by an F420H2-dependent reductase. *Angew. Chem. Int. Ed. Engl.* **2020**. DOI:10.1002/anie.202008035.

(12) (a) Wiebach, V.; Mainz, A.; Siegert, M. J.; Jungmann, N. A.; Lesquame, G.; Tirat, S.; Dreux-Zigha, A.; Aszodi, J.; Le Beller, D.; Sussmuth, R. D., The anti-staphylococcal lipolanthines are ribosomally synthesized lipopeptides. *Nat. Chem. Biol.* **2018,** *14*, 652; (b) Wiebach, V.; Mainz, A.; Schnegotzki, R.; Siegert, M. J.; Hugelland, M.; Pliszka, N.; Sussmuth, R., An amphipathic alpha-helix guides maturation of the ribosomally-synthesized lipolanthines. *Angew. Chem. Int. Ed. Engl.* **2020**. DOI:10.1002/anie.202003804.

(13) Lu, J.; Wu, Y.; Li, J.;Li, Y.; Zhang, Y.; Bai, Z.; Zheng, J.; Zhu, J.; Wang, H., Lanthipeptide Synthetases Participate the Biosynthesis of 2-Aminovinyl-Cysteine Motifs in Thioamitides. Unpublished Manuscript.

(14) Soding, J.; Biegert, A.; Lupas, A. N., The HHpred interactive server for protein homology detection and structure prediction. *Nucleic Acids Res.* **2005,** *33*, W244.

(15) MacNeil, D. J.; Gewain, K. M.; Ruby, C. L.; Dezeny, G.; Gibbons, P. H.; MacNeil, T., Analysis of Streptomyces avermitilis genes required for avermectin biosynthesis utilizing a novel integration vector. *Gene* **1992,** *111*, 61.
